# The Late-Stage Steps of *Burkholderia cenocepacia* Protein *O*-Linked Glycan Biosynthesis Are Conditionally Essential

**DOI:** 10.1101/2024.11.21.624715

**Authors:** Leila Jebeli, Taylor A. McDaniels, Duncan T. T. Ho, Hamza Tahir, Nicholas L. Kai-Ming, Molli Mcgaw, Kristian I. Karlic, Jessica M. Lewis, Nichollas E. Scott

## Abstract

Periplasmic *O*-linked protein glycosylation is a highly conserved process observed across the *Burkholderia* genus. Within Burkholderia, protein glycosylation requires the five gene cluster known as the *<u>O</u>*-<u>g</u>lycosylation <u>c</u>luster (OGC, *ogcXABEI*) which facilitates the construction of the *O*-linked trisaccharide attached to periplasmic proteins. Previous studies have reported conflicting results regarding the essentiality of *ogcA*, predicted to be responsible for the addition of the final carbohydrate of the *O*-linked trisaccharide and *ogcX,* the putative *O*-linked glycan flippase. Within this work, we aimed to dissect the impact of the loss of *ogcA* and *ogcX* on *Burkholderia cenocepacia* viability. We demonstrate that the loss of either *ogcA* or *ogcX* are detrimental if glycosylation is initiated leading to marked phenotypic effects. Proteomic analysis supports that the loss of *ogcA*/*ogcX* both blocks glycosylation and drives pleotropic effects in the membrane proteome, resulting in the loss of membrane integrity. Consistent with this, strains lacking *ogcA* and *ogcX* exhibit increased sensitivity to membrane stressors including antibiotics and demonstrate marked changes in membrane permeability. These effects are consistent with fouling of the undecaprenyl pool due to dead-end *O*-linked glycan intermediates, and consistent with this, we show that modulation of the undecaprenyl pool through the overexpression of undecaprenyl pyrophosphate synthase (UppS) or the OGC flippase (OgcX) restores viability while expression of early-stage OGC biosynthesis genes (*ogcI* and *ogcB*) reduce *B. cenocepacia* viability. These findings demonstrate disrupting *O*-linked glycan biosynthesis or transport appears to dramatically impact *B. cenocepacia* viability, supporting the assignment of *ogcA* and *ogcX* as conditionally essential.

**Importance:** Protein glycosylation, a conserved process in *Burkholderia* species, utilizes glycans generated by the *O*-glycosylation cluster (OGC), which is composed of five genes (*ogcX, ogcA*, *ogcB*, *ogcE*, and *ogcI*). In this study, we demonstrate that the loss of *ogcA* or *ogcX* significantly affects the physiology of *Burkholderia cenocepacia*. Using complementary genetic approaches and proteomic techniques, we show that the loss of *ogcA* or *ogcX* blocks glycosylation, alters the cell membrane, and sensitizes cells to stressors such as antibiotics. This increased sensitivity to membrane stress is consistent with the accumulation of dead-end *O*-linked glycan intermediates, which sequester the limited and essential undecaprenyl pool within *B. cenocepacia*. These findings highlight that *ogcA* and *ogcX* are conditionally essential for *B. cenocepacia* survival and provides mechanistic insight into how protein glycosylation fidelity—the use of specific glycans for protein glycosylation—is controlled in *Burkholderia* species.

## Introduction

Across bacterial species many surface polysaccharides are synthesized utilising the essential polyisoprenoid lipid undecaprenyl phosphate (Und-P) ^1,2^. Functioning as a chemical carrier, Und-P provides an assembly point for the stepwise construction of diverse lipid-linked oligosaccharide units at the inner face of the cytoplasmic membrane within Gram-negative and Gram-positive species ^1,2^. By providing a tethering point to enable glycan elongation, Und_-_P allows the construction of a range of glycans, which upon translocation across the cytoplasmic membrane can be integrated into glycoconjugates including peptidoglycan, cell wall teichoic acids, O-antigens, capsules, and exopolysaccharides ^3,4^. While Und-P and its phosphorylated form undecaprenyl pyrophosphate (Und-PP) are important for the construction of surface carbohydrates, these molecules account for less than 1% of the total lipid pool ^5^, resulting in competition for available Und-P and the need for rapid flux of oligosaccharide units through the Und-P/Und-PP cycle ^6,7^. Under normal physiological conditions, *de novo* synthesis and recycling of Und-P/Und-PP are able to meet physiological demands ^6,7^, yet it is increasingly recognised that genetic manipulations that result in Und-PP-glycan intermediates that sequester the Und-P/Und-PP pool can dramatically impact cellular viability, morphology and growth ^6–11^. Understanding Und-PP-glycan intermediates with the capacity to foul the Und-P/Und-PP pool has the potential to not only aid antimicrobial development ^12,13^ but also explain the mechanisms underpinning the homogeneity observed within the glycan structures of glycoconjugates ^14–16^.

Ensuring only glycans possessing specific structural configurations, herein referred to as glycan fidelity, are translocated across the cytoplasmic membrane ensures homogeneity within oligosaccharide units constructed utilising the Und-P/Und-PP pool ^3,4^. For many glycoconjugates generated using Und-PP linked oligosaccharides, the addition of specific glycan components is a pre-requisite for effective translocation across the cytoplasmic membrane by the actions of dedicated translocases known as flippases ^17–19^. Of the known flippases the Wzx enzymes represent a highly diverse family of enzymes ^19,20^ responsible for the translocation of a range of carbohydrates including O-antigens ^21^, capsules ^22,23^, exopolysaccharides ^24^, and the enterobacterial common antigen (ECA) ^10^. While Wzx flippases were initially thought to possess limited capacity for the recognition of glycan structures ^25,26^, it has become clear multiple features of their cognate glycans, such as specific branching or terminal sugars in addition to the initiating sugar influence translocation efficiency when Wzx enzymes are expressed under native-like conditions ^3,15,27,28^. This specificity results in discrete Wzx flippases being observed genetically linked to their cognate glycan clusters ^15,22,23,27^ with Wzx enzymes suggested to function as quality control mechanisms ^16^, ensuring only “complete” glycans are effectively translocated across the cytoplasmic membrane. The specificity for complete glycans by Wzx flippases in concert with the extension of glycans beyond key points commit Und-PP-linked oligosaccharides to the generation of specific glycans ^11,25,29–32^. In cases where Und-PP-linked intermediates are unable to be completed this drives detrimental phenotypes due to the sequestration of the Und-P/Und-PP pool including growth defects, enhanced sensitivity to membrane stresses and alterations in cellular morphology ^6,7,11^. Importantly the profound effects of Und-P/Und-PP sequestration is known to rapidly lead to the emergence of suppressor mutations, which complicates the analysis of Und-P/Und-PP glycan biosynthesis pathways by obfuscating the role of specific glycan biosynthesis genes ^9,33,34^. While these potential confounding effects are now well appreciated for the O-antigens ^6,11^, capsule ^35^ and ECA ^7^, this phenomenon has not been explored within glycans used for other glycoconjugates such as protein glycosylation.

Protein glycosylation is widespread across bacterial species ^36,37^ including members of the *Campylobacter* ^38,39^, *Neisseria* ^40–44^, and *Burkholderia* ^45–47^ genera. To date, several Und-P/Und-PP dependent glycosylation systems have been described in Gram-negative species, which utilise the Und-P lipid pool to enable periplasmic glycosylation leading to glycosylation of tens to hundreds of proteins ^37,48–50^. While similarities within the biosynthetic clusters used for protein glycosylation systems and lipooligosaccharide biosynthetic clusters have been previously highlighted ^36,37^, a notable difference within well-characterised glycosylation systems is the use of ABC transporter-based flippases ^36,37^ as opposed to Wzx flippases. ABC transporter-based flippases, such as PglK (also known as WlaB^51^) within *Campylobacter* sp. ^52,53^ and PglF within *Neisseria* sp. ^54–56^ are dispensable for glycosylation ^53,56^ as well as possess relaxed glycan specificities as demonstrated using heterogenous expression studies for PglK ^53^ and the observation of glycan microheterogeneity within *Neisseria* ^57,58^. The tolerance of these glycosylation pathways to both glycan alterations, as well as the loss of their flippases, is in stark contrast to the strict specificity and essentiality seen within other Und-P/Und-PP dependent systems, yet not all Und-P/Und-PP dependent protein glycosylation systems possess ABC transporter flippases.

Within *Burkholderia* species, periplasmic glycosylation is highly conserved ^45–47^, leading to nearly exclusive modification of serine residues ^45^ with a trisaccharide corresponding to β-Gal-(1,3)–α-GalNAc-(1,3)–β-GalNAc ^47^. *Burkholderia O*-linked glycan assembly is mediated by a single gene cluster known as the *<u>O</u>*-linked <u>G</u>lycosylation <u>C</u>luster (*OGC*) consisting of five genes (*ogcX*, *A*, *B*, *E* and *I* corresponding to BCAL3114 to BCAL3118 within *Burkholderia cenocepacia* respectively) ^47^. Our current model of glycan assembly (**Figure 1A**) supports OgcE (BCAL3117) as a UDP-Glc / UDP-GlcNAc epimerase, OgcI (BCAL3118) as the initiating glycotransferase for the *O*-linked glycan, OgcB (BCAL3116) and OgcA (BCAL3115) work as extending glycosyltransferases responsible for the addition of the second GalNAc and final Gal residues respectively, and the putative Wzx flippase OgcX (BCAL3114), responsible for the translocation of the glycan across the cytoplasmic membrane ^47^ where it can then be transferred to glycoproteins by the oligosaccharyltransferase PglL (BCAL0960) ^47,59^. Previous studies have provided experimental evidence for the function of OgcB, OgcE and OgcI ^47^ yet contradictory results have been obtained for the enzymes OgcA and OgcX. While it has been noted that despite multiple attempts an *ogcA* knockout has been unable to be generated ^47^, the loss of *ogcX* has been previously reported to be tolerated within *B. cenocepacia* ^47^. Similarly, saturation transposon studies from several *Burkholderia* species ^60–62^ have revealed variability in the ability to disrupt *ogcA* and *ogcX,* with some studies concluding these enzymes are essential ^62^ while others suggest that at least for *ogcA* disruptions within this gene may be tolerated ^61^. These observations support that unlike the glycosylation systems of the *Campylobacter* and *Neisseria* genera, disruptions in glycan biosynthesis steps may not be tolerated within *Burkholderia.* These contradictory results support further assessments are required to dissect the impact of loss of *ogcA* and *ogcX* on viability and protein glycosylation.

**Figure 1.**
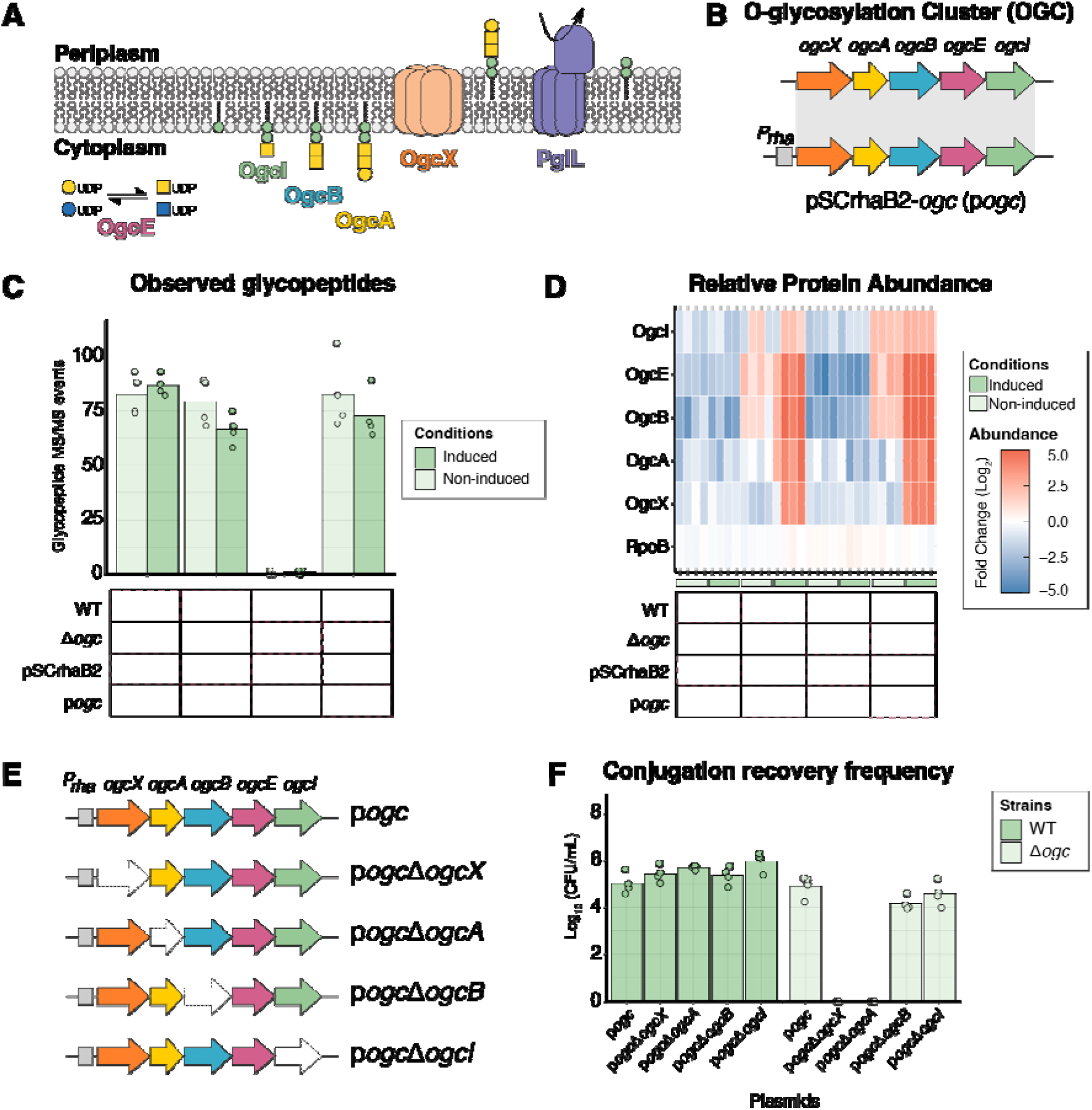
Loss of *ogc*X and *ogc*A impacts viability of *B. cenocepacia* Δ*ogc*. **A)** Diagram of the *O*-glycosylation biosynthesis pathway in *Burkholderia* species ^47^. *O*-glycosylation biosynthesis involves the stepwise construction of a conserved trisaccharide composed of Gal-GalNAc_2_ by the glycosyltransferases, OgcI, OgcB and OgcA, with the nucleotide-sugar donors UDP-GalNAc and UDP-Gal generated by the epimerase OgcE. The resulting trisaccharide is then thought to be translocated across the inner membrane by the putative flippase, OgcX. Once in the periplasm, the glycans can be transferred to proteins by the oligosaccharyltransferase, PglL. **B)** Graphic representation of the genetic organization of the *ogc* cluster (*ogc*X, A, B, E and I assigned as BCAL3114–BCAL3118 within *B. cenocepacia* J2315) and its integration into the rhamnose-inducible vector pSCrhaB2, resulting in the generation of the pSCrhaB2-*ogc* (p*ogc*). **C-D)**Quantification of glycopeptide and protein levels encoded by the *ogc* cluster identified from whole cell proteomic analysis of *B. cenocepacia* WT and Δ*ogc* containing either pSCrhaB2 or p*ogc* with and without induction with 1% rhamnose (n=4). **E)** Graphic representation of p*ogc* mutagenesis removing *ogc*X, *ogc*A, *ogc*B and *ogc*I. **F)** Conjugation of p*ogc* derivatives into *B. cenocepacia* K56-2 WT and Δ*ogc* reveals differences in recovery rates of p*ogc*Δ*ogc*X and p*ogc*Δ*ogc*A within *B. cenocepacia* Δ*ogc* (n=4).

Within this work, we sought to assess the requirement of *ogcA* and *ogcX* for protein glycosylation and the impact of the loss of these *ogc* components on *B. cenocepacia.* Utilising complementary genetic approaches, including *ogc* complementation within strains lacking the *ogc* cluster ^47,63^ as well as glycosylation inducible strains containing the initiating transferase, *ogcI,* under rhamnose control ^64^, we assess the proteomic and physiological impacts of the loss of *ogcA* and *ogcX.* We demonstrate that in the absence of glycosylation initiation, *ogcA* and *ogcX* are dispensable, yet upon glycosylation initiation, loss of these genes leads to growth defects, widespread proteomic impacts, changes in the membrane integrity and the absence of observable glycosylation. While these changes can be complemented and compensated by the modulation of enzymes associated with *de novo* synthesis of Und-P as well as the overexpression of the flippase OgcX, we observe that the modulation of early steps in the *O*-linked glycan biosynthesis pathways can also drive changes in viability. This data supports that the construction of the *O*-linked glycan in *B. cenocepacia* is finely tuned with *ogcA* and *ogcX* conditionally detrimental if glycosylation is initiated.

## Methods

### Bacterial strains and growth conditions

Bacterial strains used in this study are listed in **Supplementary Table 1**. Strains were cultured in Lysogeny Broth (LB) or on LB agar (1.5–2% w/v), prepared in accordance with the manufacturer’s instructions containing 0.5% NaCl (BD™, New Jersey, USA). Liquid cultures were incubated overnight at 37°C with shaking, while agar plates were incubated at 37°C overnight for *Escherichia coli* and 24–72 hours for

*B. cenocepacia*. When required, LB was supplemented with 200 μg/mL diaminopimelic acid (DAP; Sigma) to support growth of *E. coli* RHO3 ^65^. Antibiotics were added to cultures to select or maintain plasmids/ transconjugants at a final concentration of 50 μg/mL trimethoprim for *E. coli* and 100 μg/mL for *B. cenocepacia*, 20 μg/mL tetracycline for *E. coli* and 150 μg/mL for *B. cenocepacia* and 50 μg/mL kanamycin for *E. coli*. To counter-select helper and donor *E. coli* strains during triparental and quadparental mating, ampicillin at 100 μg/mL and polymyxin B at 25 µg/mL were added ^66^. Culturing of strains containing the temperature-sensitive plasmid pFlptet (see **Supplementary Table 2**) was undertaken at 30°C with cultures shifted to 37°C to enable plasmid curing ^67^. Curing of pDAI-SceI-SacB was achieved by sucrose counterselection as previously described with 5% sucrose in LB lacking NaCl ^66^. Induction of *B. cenocepacia* strains was performed by either adding 20% filter-sterilised L-rhamnose monohydrate (Sigma) or 1 mM 4-Isopropylbenzoic acid (cumate, Sigma) to LB broth or LB agar at a concentration of 0.05%-1% for rhamnose and 100 μM for cumate. For non-induced controls, an equivalent volume of solvent (sterile water for rhamnose and ethanol for cumate) was added to the culture media.

### Recombinant DNA methods

Oligonucleotides used in this study are listed in **Supplementary Table 3**, with all PCR amplifications for cloning carried out using Q5 DNA polymerase (New England Biolabs) with the addition of 2% DMSO for the amplification of *B. cenocepacia* DNA, due to its high GC content. Genomic DNA isolations were performed using an EZ-10 Spin Column Bacterial Genomic DNA Mini-Preps Kit (Bio Basic), while PCR clean up and restriction digest purifications were performed with a Zymoclean Gel DNA Recovery Kit (ZymoResearch). pGPI-SceI ^68^ mutagenesis plasmids were constructed using Gibson assembly ^69^ with DNA fragments flanking genes of interest amplified by Q5-based PCR. *Sma*I-linearized pGPI-SceI and Q5-amplified fragments were assembled using the NEBuilder® HiFi DNA master mix according to the manufacturer’s instructions (New England Biolabs). pSCrhaB2 ^64^ and pMLBAD ^70^ based expression vectors containing genes of interest were generated using Gibson Assembly or restriction endonuclease-based cloning. For Gibson Assembly, PCR-amplified fragments were introduced into *Nde*I/*Xba*I-linearized pSCrhaB2 or PCR-derived pMLBAD vector backbones using the NEBuilder® HiFi DNA master mix. For restriction endonuclease-based cloning, PCR-amplified fragments were digested with *Nde*I/*Xba*I and then ligated into *Nde*I/*Xba*I-linearized pSCrhaB2 overnight at 16°C using T4 Ligase (New England Biolabs). Mutagenesis of pSCrhaB2-ogc to remove *ogc*A, *ogc*B, *ogc*I and *ogc*X was achieved by assembling PCR-amplified fragments into *Nde*I/*Hind*III-linearized pSCrhaB2 using the NEBuilder® HiFi DNA master mix.

Prior to Gibson Assembly and restriction cloning all DNA fragments were assessed for correctness based on size using agarose gel electrophoresis. Ligated/assembled plasmid mixtures were introduced into chemically competent *E. coli* PIR2 cells via heat shock transformation ^71^, plated on selective media and screened using colony PCR with GoTaq® Green Master Mix (Promega). All plasmids used in this study have been confirmed by either Sanger sequencing using the Australian Genome Research Facility (Melbourne, Australia) or nanopore plasmid sequencing using Plasmidsaurus (SNPsaurus LLC, Eugene, OR). Donor plasmids for biparental and quadparental mating were further isolated and introduced into *E. coli* RHO3 ^65^ via electroporation ^71^ to improve conjugation efficiency into *B. cenocepacia*. A complete list of the plasmids used in this study, along with a summary of their construction, is provided in **Supplementary Table 2**.

### Conjugation of plasmids into *B. cenocepacia*

Plasmids were introduced into *B. cenocepacia* K56-2 via one of three approaches: I) biparental mating using the conjugative diaminopimelic acid auxotroph *E. coli* RHO3 ^65^ for the introduction of pFLPtet or pDAI-SceI-SacB; II) triparental mating ^66^ using *E. coli* PIR2 containing pRK2013 (helper plasmid) ^72^ and *E. coli* PIR2 carrying pGPI-SceI-derived vectors or pSCrhaB2/pMLBAD expression vectors (donor plasmid); or III) quadparental mating using *E. coli* PIR2 containing pRK2013 ^72^, *E. coli* PIR2 containing pTNS3 (Tn7 transposase plasmid) ^73^ and *E. coli* RHO3 carrying pUC18T-mini-Tn7T-Tp-rha-ogcI (donor plasmid) for the chromosomal integration of miniTn7-rha-ogcI. Conjugations were allowed to proceed for 24 hours, then successful transconjugants were selected with trimethoprim, and the introduction of plasmids of interest confirmed using PCR-based screening. For the generation of unmarked, non-polar deletions using pGPI-SceI and the removal of the trimethoprim resistance marker within the integrated miniTn7, additional rounds of conjugation were undertaken to introduce pDAI-SceI-SacB ^68,74^ or pFLPtet^67^ respectively using tetracycline based selection. pGPI-SceI/pDAI-SceI-SacB-based mutagenesis as well as FLP-based removal of the trimethoprim resistance marker within Tn7 were confirmed using screening oligonucleotides (**Supplementary Table 3**) by colony PCR using GoTaq® Green Master Mix. Numeration of transconjugants derived from the introduction of pSCrhaB2-*ogc* and its derivatives into *B. cenocepacia* wildtype (WT) or *B. cenocepacia* Δ*ogc* were assessed across three independent conjugations.

### Spot plate assays

Colony size morphology and membrane stress tolerance assessments were performed as previously described ^75–77^ using spot plate assays. For spot plate assays, strains of interest were grown overnight in LB broth without induction, normalised to an OD_600_ of 1.0 and then serially diluted from 10^-1^ to 10^-6^. 10 μL of diluted cultures were manually spotted onto LB plates with and without inducers (1% Rhamnose /100 μM cumate). 0.01% SDS or 2% NaCl were added to plates to assess membrane integrity and the impact of osmotic stress on viability with and without inducers. For assays of *B. cenocepacia* strains containing pSCrhaB2-or pMLBAD-based expression vectors, trimethoprim was added to plates to ensure the maintenance of plasmids. For antimicrobial resistance, spot assays concentrations corresponding to at or below the broth dilution defined MIC assays (**Supplementary Figure 10**) were used for trimethoprim (8 μg/mL), tetracycline (16 μg/mL), chlorhexidine (4 μg/mL), ceftazidime (4 μg/mL), and rifampicin (32 μg/mL) on cation-adjusted Mueller–Hinton agar. Spots were allowed to dry, and plates were incubated for 48 hours at 37°C before being imaged. All spot assays were assessed using at least three independent replicates. Colony area measurements were undertaken using Fiji ^78^, assessing the size of 20 colonies per spot assay with three independent assays conducted with t-tests used to determine significant changes in colony size as well as viable counts between induced and non-induced conditions.

### Microtiter plate-based viability and growth kinetics assays

To assess viability and growth kinetics, Microtiter 96-well-based growth assays were undertaken as previously described ^63^. Bacterial strains were grown overnight in LB, normalised to an OD_600_ of 1.0 and diluted 1 in 100 with LB supplemented with or without combinations of inducers and/or membrane stressors (0.01% SDS or 2% NaCl). 200 μL of diluted cultures were added to 96-well flat-bottom plates and incubated at 37°C with shaking at 200 rpm in a CLARIOstar plate reader (BMG LABTECH, Inc.), measuring the optical density at 600 nm every 10 minutes over 24 hours. Growth assays were assessed in at least technical triplicate across three independent biological replicates.

### Fluorescence Detection of the permeability of inner and outer membranes

Permeability of the inner membrane was assessed using Hoechst 33342 (Abcam) as previously described ^79–81^. Briefly, bacterial strains were grown overnight in LB (with and without 1% Rhamnose), normalised to an OD_600_ of 2.0, washed three times with PBS and dispensed at 100 µL aliquots into 96-well black clear-bottomed plates. 100 µL of PBS and heat-killed bacteria were added to the plates as controls. 100 µL of 2.5 µM Hoechst 33342 in PBS was added to wells to generate bacterial suspensions with an OD_600_ of 1.0 and a final concentration of 1.25 µM Hoechst 33342. Fluorescence was measured in a CLARIOstar plate reader using an excitation of 361nm and emission detection at 486nm. The fluorescence intensity was first normalised by assigning the value of 100 to the heat-killed and 0 to the non-induced parental strain. To assess outer membrane permeability, 1-*N*-phenylnaphthylamine (NPN, Thermo Fisher Scientific) as previously described ^82–84^. Cultures were normalised as above, washed three times with 5 mM HEPES, pH 7.4, 10 mM sodium azide and dispensed at 100 µL aliquots into 96-well black clear-bottomed plates. 100 µL of 5 mM HEPES, pH 7.4, 10 mM sodium azide was added to the plates as a blank. 100 µL of 20 µM NPN in 5 mM HEPES, pH 7.4, 10 mM sodium azide was added to wells to generate bacterial suspensions with an OD_600_ of 1.0 and a final concentration of 100 µM NPN. Fluorescence was measured in a CLARIOstar plate reader using an excitation of 350nm and emission detection at 420nm. Hoechst 33342 and NPN assays were assessed in technical triplicate across five independent biological replicates comparing changes in fluorescence intensity to the non-induced parental strain. To assess viability and ensure comparable cell numbers, the OD-adjusted bacterial suspensions were serially diluted and plated on LB and viable cells numerated (**Supplementary Figure 11**). The fluorescence intensity from both assays was normalised against the determined cell number.

### Preparation of proteomic samples

*B. cenocepacia* cultures for proteomic analysis were grown overnight with or without induction with rhamnose and cumate as outlined above with shaking at 180 rpm. Overnight cultures were normalised to an OD_600_ of 1.0 and then collected by centrifugation at 10,000 x *g* at 4°C for 10 minutes, washed 3 times with ice-cold PBS and then snap frozen at-80°C until processing. Frozen whole cell samples were prepared for analysis using sodium deoxycholate (SDC) based lysis and the in-StageTip preparation approach as previously described ^85^. OgcX complementation assays were undertaken using SDS-based lysis and S-trap sample preparation to improve the detection of OgcX. For in-StageTip preparation, cells were resuspended in 4% SDC, 100 mM Tris pH 8.0 and boiled at 95°C with shaking (2000 rpm) for 10 minutes to solubilise the proteome. Samples were allowed to cool for 10 minutes and then boiled for a further 10 minutes (95°C, 2000 rpm) before the protein concentrations were determined by bicinchoninic acid assays (Thermo Fisher Scientific). Samples were reduced/alkylated with the addition of Tris-2-carboxyethyl phosphine and chloroacetamide (final concentration 10 mM and 40 mM, respectively), and samples were incubated in the dark for 0.5-1 hour at 45°C. Following reduction/alkylation, samples were digested overnight with Trypsin (1/50 w/w Solu-trypsin, Sigma) at 37°C with shaking at 1000rpm. Digests were then quenched with the addition of 1.25 volumes of isopropanol before being cleaned up using SDB-RPS (Sigma) StageTips.

For S-trap-based preparation, cells were resuspended in 4% sodium dodecyl sulphate (SDS), 100 mM Tris pH 8.0, and boiled at 95°C with shaking (2000 rpm) for 10 minutes; the protein concentrations were then determined by bicinchoninic acid assays, and samples were reduced/alkylated as above. Samples were acidified to 1.2% phosphoric acid and diluted with seven volumes of S-trap wash buffer (90% methanol, 100mM tetraethylammonium bromide pH 7.1) before being loaded onto S-trap mini spin columns (Protifi) and washed 3 times with 400 µL of S-trap wash buffer. Samples were then digested with 2 µg of Trypsin (a 1:50 protease/protein ratio) in 100 mM tetraethylammonium bromide overnight at 37 °C before being collected by centrifugation with washes of 100 mM tetraethylammonium bromide, followed by 0.2% formic acid followed by 0.2% formic acid / 50% acetonitrile before being dried by vacuum centrifugation at room temperature and stored at-20°C.

SDB-RPS StageTip-based clean-up of digests was undertaken by adjusting or resuspending samples in 50% isopropanol, 1% trifluoroacetic acid and then loading them on in-house created SDB-RPS StageTips, which were prepared according to previously described protocols ^85–87^. Briefly, five frits of SDB-RPS were excised using a blunt 16-gauge Hamilton needle and loaded into 200 μL tips. SDB-RPS StageTips were placed in a Spin96 tip holder ^86^ to enable batch-based spinning of samples and tips conditioned with 100% acetonitrile, followed by 30% methanol, 1% trifluoroacetic acid, followed by 90% isopropanol, 1% trifluoroacetic acid with each wash spun through the column at 1000 x *g* for 3 minutes. Acidified isopropanol/peptide mixtures were loaded onto the SDB-RPS columns and spun through tips before being washed with 90% isopropanol, 1% trifluoroacetic acid, followed by 90% ethyl acetate, 1% trifluoroacetic acid, followed by 1% trifluoroacetic acid in Milli-Q water. Peptide samples were eluted with 80% acetonitrile, 5% ammonium hydroxide and dried by vacuum centrifugation at room temperature and stored at-20°C.

### LC-MS analysis of DDA samples

Cleaned-up peptide samples were re-suspended in Buffer A* (2% acetonitrile, 0.1% trifluoroacetic acid in Milli-Q water) and separated using a two-column chromatography set-up on a Dionex Ultimate 3000 UPLC composed of a PepMap100 C18 20 mm x 75 μm trap and a PepMap C18 500 mm x 75 μm analytical column (Thermo Fisher Scientific) coupled to a Orbitrap Fusion™ Lumos™ Tribrid™ Mass Spectrometer (Thermo Fisher Scientific) with datasets collected with and without the use of the FAIMS Pro interface (Thermo Fisher Scientific). 145-minute gradients were run for each sample, with samples loaded onto the trap column with 98% Buffer A (2% acetonitrile, 0.1% formic acid in Milli-Q water) and 2% Buffer B (80% acetonitrile, 0.1% formic acid) with peptides separated by altering the buffer composition from 2% Buffer B to 28% B over 126 minutes, then from 28% B to 40% B over 9 minutes, then from 40% B to 80% B over 3 minutes, the composition was held at 80% B for 2 minutes, and then dropped to 2% B over 2 minutes and held at 2% B for another 3 minutes. For datasets collected without the use of the FAIMS Pro interface, the Lumos™ Mass Spectrometer was operated in a data-dependent mode, switching between the collection of a single Orbitrap MS scan (450-2000 m/z, maximal injection time of 50 ms, an Automatic Gain Control (AGC) of maximum of 4*10^5^ ions and a resolution of 60k) acquired every 3 seconds followed by Orbitrap MS/MS HCD scans of precursors (NCE 30%, maximal injection time of 80 ms, an AGC set to a maximum of 1.25*10^5^ ions and a resolution of 30k). For datasets collected with the FAIMS Pro interface, a data-dependent stepped FAIMS approach was utilised with three different FAIMS CVs-25,-45 and-65, as previously described ^88^. For each FAIMS CV a single Orbitrap MS scan (450-2000 m/z, maximal injection time of 50 ms, an AGC of maximum of 4*10^5^ ions and a resolution of 60k) was acquired every 1.2 seconds followed by Orbitrap MS/MS HCD scans of precursors (NCE 30%, maximal injection time of 80 ms, an AGC set to a maximum of 1.25*10^5^ ions and a resolution of 30k). HCD scans containing HexNAc-associated oxonium ions (204.0867, 138.0545 and 366.1396 m/z) triggered two additional product-dependent MS/MS scans ^89^ of potential glycopeptides; an Orbitrap EThcD scan (NCE 15%, maximal injection time of 150 ms, AGC set to a maximum of 2*10^5^ ions with a resolution of 30k using the extended mass range setting to improve the detection of high mass glycopeptide fragment ions ^90^) and a stepped collision energy HCD scan (using NCE 35% with 5% Stepping, maximal injection time of 150 ms, an AGC set to a maximum of 2*10^5^ ions and a resolution of 30k).

### LC-MS analysis of DIA samples

Cleaned-up peptide samples were resuspended in Buffer A* and separated using a two-column chromatography set-up on a Dionex Ultimate 3000 UPLC composed of a PepMap100 C18 20 mm x 75 μm trap and a PepMap C18 500 mm x 75 μm analytical column. Samples were concentrated onto the trap column at 5 μL/min for 5 minutes with 98% Buffer A and 2% Buffer B, then infused into an Orbitrap Fusion™ Lumos™ Tribrid™ Mass Spectrometer at 300 nL/minute via the analytical column. 125-minute analytical runs were undertaken by altering the buffer composition from 3% Buffer B to 25% B over 112 minutes, then from 25% B to 40% B over 4 minutes, then from 40% B to 80% B over 1 minute. The composition was held at 80% B for 3 minutes and then dropped to 3% B over 1 minute before being held at 3% B for another 4 minutes. Data was collected in a Data-independent manner with a single MS1 event (120k resolution, AGC 1*10^6^, 350-1400 m/z) and 50 MS2 scans (NCE 30%, 30k resolution, AGC 1*10^6^, 200-2000 m/z and a maximal injection time of 55 ms) of a width of 13.7 m/z collected over the mass range of 360 to 1033.5 m/z.

### DDA-based proteomic analysis

DDA datasets were analysed using MSFragger (versions 18.0, 19.0, 20.0 or 22.0) ^91–93^. Samples were searched with a Tryptic specificity, allowing a maximum of two missed cleavage events and Carbamidomethyl set as a fixed modification of Cysteine while oxidation of Methionine allowed as a variable modification. For searches to identify canonical Bukrholderia glycosylation the *Burkholderia* glycans HexHexNAc_2_ (elemental composition: C_22_O_15_H_36_N_2_, mass: 568.2115 Da) and Suc-HexHexNAc_2_ (elemental composition: C_26_O_18_H_40_N_2_, mass: 668.2276 Da) were included as variable modifications at Serine in line with the strong preference for PglL glycosylation at Serine residues ^45^. To allow the assessment of glycoforms within strains lacking *ogc* genes the *Burkholderia* glycans HexHexNAc_2_, Suc-HexHexNAc_2_, HexNAc_2_ (elemental composition: C_16_O_10_H_26_N_2_, mass: 406.1488 Da) and HexNAc (elemental composition: C_8_O_5_H_13_N, mass: 203.0794 Da) were included as variable modifications at Serine. For all glycopeptide searches the glycan fragment ions were defined as 204.0866, 186.0760 168.0655, 366.1395, 144.0656, 138.0550, 466.1555 and 407.1594. A maximum mass precursor tolerance of 20 ppm was allowed at both the MS1 and MS2 levels. Samples were searched against the *B. cenocepacia* reference proteome J2315 (Uniprot accession: UP000001035, 6,993 proteins, downloaded July 24^th^ 2020) ^94^ supplemented with the proteins RhaS (HTH-type transcriptional activator RhaS, P09377), RhaR (HTH-type transcriptional activator RhaR, P09378), TmpR (Dihydrofolate reductase resistance marker, P00384) and CymR (HTH-type transcriptional regulator CymR, O33453). Assessments of changes in glycosylation were undertaken based on spectral counting of glycopeptide identification events with data visualization undertaken using ggplot2 ^95^ in R. To confirm the identity of glycoforms representative HCD and EThcD spectra were annotated with the aid of the Interactive Peptide Spectral Annotator tool (http://www.interactivepeptidespectralannotator.com/PeptideAnnotator.html) ^96^.

### DIA-based proteomic analysis

DIA Datasets were searched using DIA-library free analysis within Spectronaut (version 17.6). Data files were searched against the *B. cenocepacia* K56-2 proteome (Uniprot accession: UP000011196, 7467 proteins, downloaded October 13^th^ 2020) ^97^ and *B. cenocepacia* strain J2315 (Uniprot accession: UP000001035) ^94^, merging these proteomes enabling the matching of proteins to both J2315 and K56-2 accessions. Oxidation of Methionine was allowed as a variable modification, Carbamidomethyl was set as a fixed modification of Cysteine, and protease specificity was set to Trypsin. Protein quantitation was undertaken using MaxLFQ ^98^ based analysis. The precursor PEP was altered to 0.01 from the default 0.2 to improve quantitative accuracy, with all single peptide protein matches excluded. Statistical analysis was undertaken using Perseus ^99^ with missing values imputed based on the total observed protein intensities with a range of 0.3 σ and a downshift of 2.5 σ. Biological replicates were grouped together, and student t-tests were used to compare individual groups with a minimum fold change of +/-1 considered for further analysis. Multiple hypothesis correction was undertaken using a permutation-based FDR approach allowing an FDR of 5%. Enrichment analysis using Fisher exact tests was undertaken in Perseus using Gene Ontology (GO) terms obtained from Uniprot (*B. cenocepacia* strain J2315 proteome: UP000001035) as well as against proteins previously reported as altered within K56-2 Δ*pglL,* K56-2 Δ*ogc or* K56-2 Δ*pglL*Δ*ogc* compared to K56-2 WT by Oppy *et al.* ^63^. Data visualization was undertaken using ggplot2 ^95^ in R.

## Data availability

All mass spectrometry data (RAW files, FragPipe outputs, Spectronaut outputs / experiment files, Rmarkdown scripts, and output tables) have been deposited into the PRIDE ProteomeXchange repository ^100,101^. All PRIDE accession numbers, descriptions of the associated experiments and experiment types (DIA or DDA) are provided in **Supplementary Table 4**.

## Results

### Loss of *ogcA* and *ogcX* impacts viability of *B. cenocepacia*

To dissect the impact of individual *ogc* genes on viability, we established a rhamnose-inducible vector pSCrhaB2 ^64^ containing the *ogc* cluster (pSCrhaB2-*ogc*) herein referred to as p*ogc* (**Figure 1B**), and assessed its ability to restore glycosylation within *B. cenocepacia* Δ*ogc* ^47^. Using mass spectrometry-based proteomics, we confirmed the restoration of glycosylation regardless of rhamnose induction and the absence of glycosylation within Δ*ogc* containing pSCrhaB2 empty vector alone (**Figure 1C, Supplementary Table 6**). Notably, we observed that the introduction of p*ogc* did not impact glycosylation levels within *B. cenocepacia* WT regardless of induction state (**Figure 1C**). While glycosylation was observed at comparable levels within strain/plasmid combinations containing endogenous or plasmid complemented *ogc,* proteomic analysis demonstrates OgcX and OgcA were below the limit of detection in the absence of rhamnose induction yet detectible upon induction (**Figure 1D, Supplementary table 7**). With the identification that p*ogc* restored glycosylation, we then sought to remove *ogcX* and *ogcA* within p*ogc* to assess the impact on viability with *ogcI* / *ogcB* removed as additional controls (**Figure 1E**). To assess the impact of *ogc* deletions on viability, we introduced plasmids into *B. cenocepacia* WT and Δ*ogc* by conjugation assessing the recovery of viable colonies. Conjugation efficiency was not impacted by the removal of *ogcA, B, I* or *X* within *B. cenocepacia* WT with comparable cfu/mL observed; however, within Δ*ogc* conjugations of p*ogc*Δ*ogcX* and p*ogc*Δ*ogcA* demonstrated dramatic reductions in recoverable cfu/mL (**Figure 1F**). These observations support that the loss of either *ogcX* and *ogcA* within p*ogc* impacts viability in the absence of chromosomally encoded *ogcX* and *ogcA*.

### Deleterious effects due to the loss of *ogcX* o r *ogcA* are dependent on glycosylation initiation

The absence of effects on viability in response to the loss of the *ogc* ^47,63^, yet the requirement of *ogcA* / *ogcX* alone supports that if *O*-linked glycan biosynthesis is allowed to proceed, it is deleterious after the addition of the second GalNAc, mediated by the OgcB ^47^. If true, we reasoned that controlling the initiation of glycosylation via placing OgcI under inducible control should render Δ*ogcA and* Δ*ogcX* viable. To assess this, we removed *ogcI* and chromosomally re-introduced *ogcI* under rhamnose-inducible control within a miniTn7 integrative system (**Figure 2A, Supplementary Figure 1A and 1B**), generating Δ*ogcI* Tn7-*ogcI.* Proteomic and glycoproteomic analysis confirmed the inducible control of *ogcI* as well as the appearance of glycosylation upon the addition of rhamnose in this background (**Supplementary Figure 2B and 2C**). Following verification of Δ*ogcI* Tn7-*ogcI,* we generated viable mutations in *ogcX and ogcA* (Δ*ogcI*Δ*ogcX* Tn7-*ogcI* and Δ*ogcI*Δ*ogcA* Tn7-*ogcI*) as well as *ogcB and ogcAB (*Δ*ogcI*Δ*ogcB* Tn7-*ogcI* and Δ*ogcI*Δ*ogcAB* Tn7-*ogcI*, **Supplementary Figure 3**). Consistent with the requirement of glycosylation initiation to drive deleterious effects, spot plate assays in the presence of rhamnose revealed reduced growth within Δ*ogcI*Δ*ogcA* Tn7-*ogcI* and Δ*ogcI*Δ*ogcX* Tn7-*ogcI* yet did not impact Δ*ogcI*Δ*ogcAB* Tn7-ogcI, Δ*ogcI*Δ*ogcB* Tn7-ogcI or Δ*ogcI* Tn7-*ogcI* growth (**Figure 2B**). Quantification of colony size within strains lacking *ogcX* or *ogcA* revealed a reduction in size to 10% of Δ*ogcI* Tn7-*ogcI*, Δ*ogcI*Δ*ogcB* Tn7-*ogcI* and Δ*ogcI*Δ*ogcAB* Tn7-*ogcI* strains in response to induction (**Figure 2C**). Consistent with these results, microtiter plate-based growth assays also revealed marked delays in the growth kinetics of strains lacking *ogcX* or *ogcA* upon induction compared to Δ*ogcI* Tn7-*ogcI* (**Supplementary Figure 4**). Importantly, across strains, quantification of glycosylation using glycoproteomics supports that while rhamnose induction led to the restoration of glycosylation within Δ*ogcI* Tn7-*ogcI,* no glycosylation was observable within strains lacking *ogcA, B, X or AB* upon rhamnose induction (**Figure 2D, Supplementary Table 8**).

**Figure 2.**
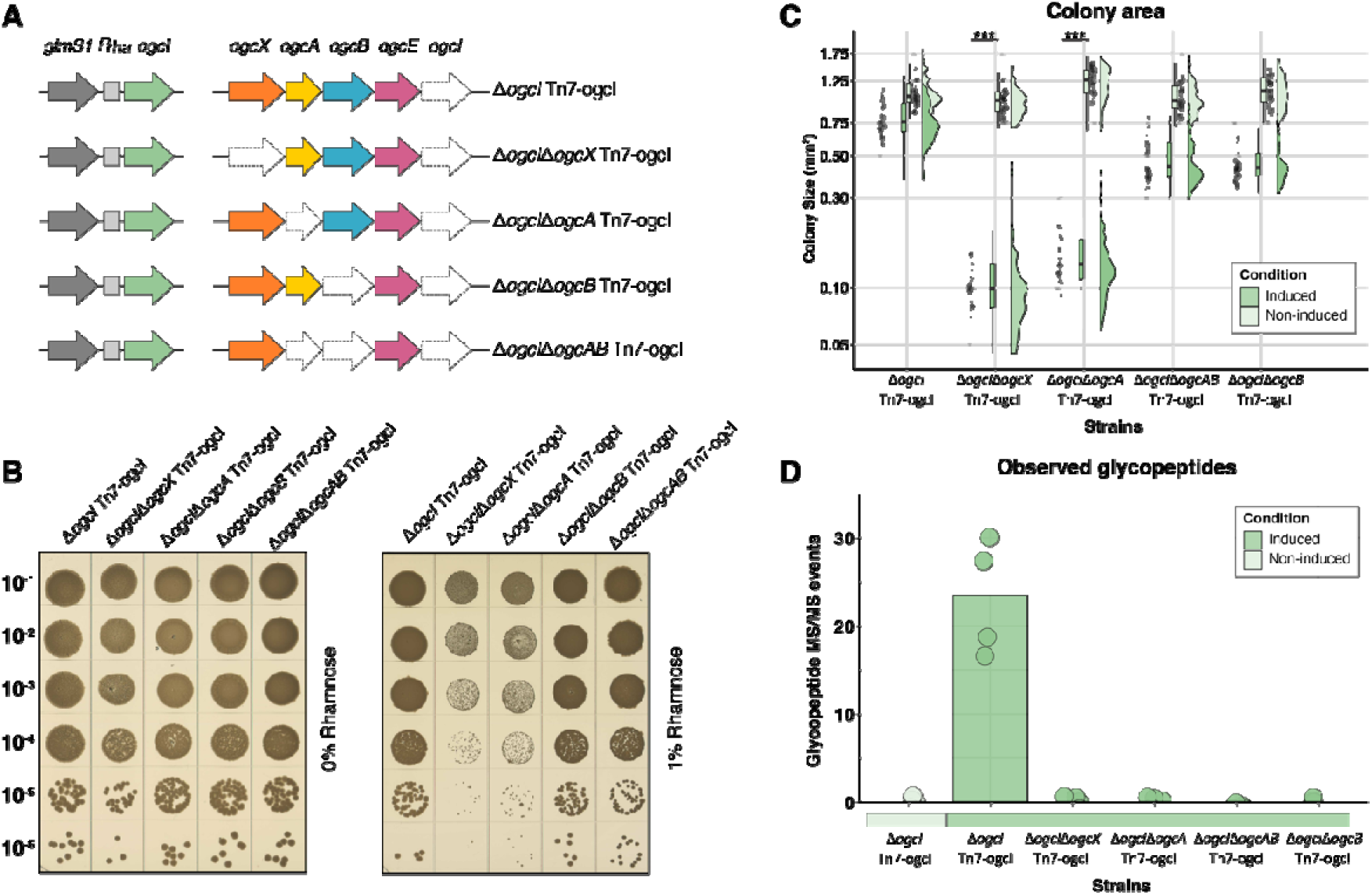
Loss of *ogcX* or *ogcA* leads to reduced growth when glycosylation is initiated. **A)** Graphic representation of strains Δ*ogcX,* Δ*ogcA,* Δ*ogcB and* Δ*ogcAB* generated within the chromosomal inducible background *B. cenocepacia* Δ*ogcI* Tn7-*ogcI*. **B-C)** Spot plate assays and half-violin box plots assessing growth of strains with and without 1% rhamnose induction. Induction results in reduced colony size of Δ*ogcX* and Δ*ogcA* strains to 10% of the size observed in *B. cenocepacia* Δ*ogcI* Tn7-*ogcI* and strains without induction (n=4). **D)** Quantification of glycopeptides identified from whole-cell proteomic analysis of strains. Glycopeptides were assessed in the absence (light green) or presence (dark green) of rhamnose induction (n=4). Induction of *ogcI* expression by 1% rhamnose led to the restoration of glycosylation only in *B. cenocepacia* Δ*ogcI* Tn7-*ogcI* in contrast, no glycosylation was observed in the Δ*ogcX*, Δ*ogcB*, and Δ*ogcAB* strains.

To support that the observed growth defects were *ogc*X and *ogc*A dependent, plasmid-based complementation was undertaken. For OgcA complementation, two rhamnose inducible vectors denoted as pSCrhaB2-*ogcA*-Met1 and pSCrhaB2-*ogcA*-Met2 corresponding to variants differing in the two potential start codons of *ogcA* (**Supplementary Figure 5**), herein referred to as p*ogcA*_Met1_ and p*ogcA*_Met2_ respectively, were generated. Upon induction both p*ogcA*_Met1_ and p*ogcA*_Met2_ within Δ*ogcI*Δ*ogcA* Tn7-*ogcI* restored colony size to comparable levels observed within Δ*ogcI* Tn7-*ogcI* (**Figure 3A**). Interestingly, Δ*ogcI*Δ*ogcA* Tn7-*ogcI* carrying the empty vector pSCrhaB2 was unable to grow in the presence of plasmid antibiotic selection conditions upon induction even with reduced rhamnose induction levels (**Supplementary Figure 6A**). Proteomic and glycoproteomic analysis confirms the induction of OgcA and restoration of glycosylation within Δ*ogcI*Δ*ogcA* Tn7-*ogcI* containing either p*ogcA*_Met1_ or p*ogcA*_Met2_ in a rhamnose dependent manner (**Figure 3B-C, Supplementary Table 9 and 10**). Consistent with spot plate assays proteomic and glycoproteomic analysis of Δ*ogcI*Δ*ogcA* Tn7-*ogcI* containing pSCrhaB2 was unable to be assessed under rhamnose induction due to loss of viability. To complement *ogcX,* we generated a modified pMLBAD vector ^70^ placing *ogcX* expression under cumate inducible control generating pCumate-*ogcX-his*, herein referred to as p*ogcX* as well as a control plasmid containing GFP, pCumate-sfGFP here in referred to as p*sfGFP*. Akin to *ogcA* complementation, the introduction of p*ogcX* into Δ*ogcI*Δ*ogcX* Tn7-*ogcI* restored colony size compared to p*sfGFP* following rhamnose induction with the presence of p*ogcX,* regardless of cumate induction, sufficient to relieve the deleterious effects of glycosylation initiation (**Figure 3D**). As with Δ*ogcI*Δ*ogcA* Tn7-*ogcI* carrying pSCrhaB2, loss of viability of Δ*ogcI*Δ*ogcX* Tn7-*ogcI* carrying p*sfGFP* was observed in the presence of plasmid antibiotic selection conditions and rhamnose induction (**Supplementary Figure 6B**). Glycoproteomic analysis confirmed the restoration of glycosylation of Δ*ogcI*Δ*ogcX* Tn7-*ogcI* containing p*ogcX* in a rhamnose-dependent manner (**Figure 3E, Supplementary Table 11**) with proteomic analysis supporting the production of OgcX in the absence of cumate (**Figure 3F, Supplementary Table 12**) consistent with low level constitutive expression of OgcX from p*ogcX*. As observed within *ogcA* complementation, proteomic/glycoproteomic analysis of Δ*ogcI*Δ*ogcX* Tn7-*ogcI* containing p*sfGFP* was unable to be assessed under rhamnose induction due to loss of viability (**Figure 3E/F**). Combined, these complementation assays support that the loss of *ogcA* and *ogcX* drive the deleterious effects on the viability of *B. cenocepacia* upon glycosylation initiation.

**Figure 3.**
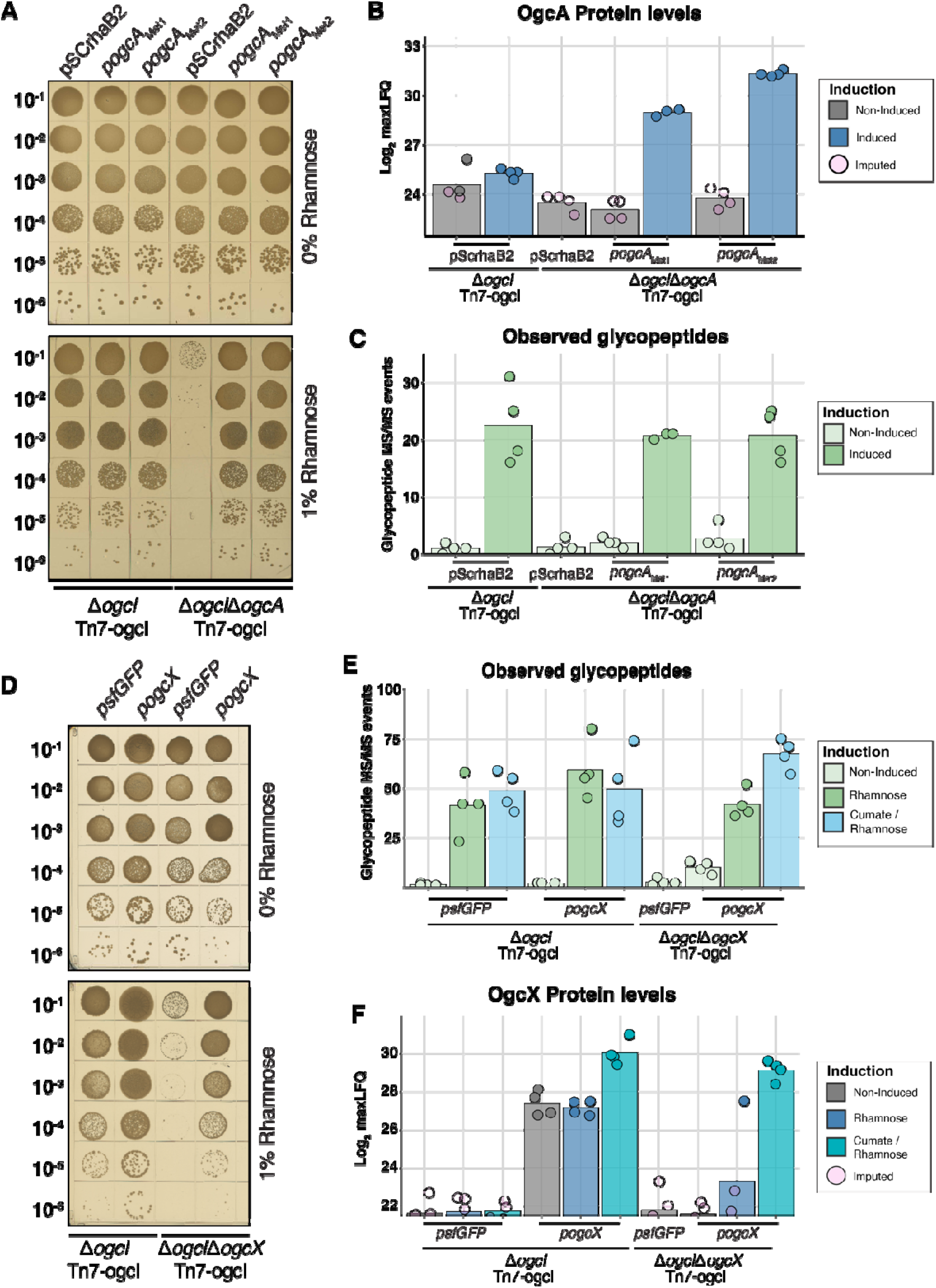
Complementation of Δ *ogcA* and Δ *ogcX* restores growth and glycosylation. **A)** Spot plate assays of *B. cenocepacia* Δ*ogcI* Tn7-*ogcI* and *B. cenocepacia* Δ*ogcI*Δ*ogcA* Tn7-*ogcI* containing pSCrhaB2 (EV), pSCrhaB2-*ogcA*-Met1 (*ogcA*-Met1), or pSCrhaB2-*ogcA*-Met2 (*ogcA*-Met2) with and without 1% rhamnose induction revealing complementation of Δ*ogcA* with *ogcA* variants restores viability and colony size. **B-C)** Quantification of glycopeptides and OgcA levels from whole-cell proteomic analysis of *B. cenocepacia* Δ*ogcI* Tn7-*ogcI* and *B. cenocepacia* Δ*ogcI*Δ*ogcA* Tn7-*ogcI* strains carrying EV, *ogcA*_Met1_, or *ogcA*_Met2_. Induction with 1% rhamnose results in the detection of OgcA within *B. cenocepacia* Δ*ogcI*Δ*ogcA* Tn7-*ogcI* and the restoration of glycosylation (n=4, Δ*ogcI* Δ*ogcA* Tn7-*ogcI ogcA*-Met1 induced group n=3). **D)** Spot plate assays of *B. cenocepacia* Δ*ogcI* Tn7-*ogcI* and *B. cenocepacia* Δ*ogcI*Δ*ogcX* Tn7-*ogcI* carrying pcumate-sfGFP (sfGFP) and pcumate-*ogcX-his* (p*ogc*X) with and without 1% rhamnose induction revealing complemented Δ*ogcX* restores viability and colony size. **E-F)** Quantification of glycopeptides and OgcX levels from whole-cell proteomic analysis of *B. cenocepacia* Δ*ogcI* Tn7-*ogcI* and *B. cenocepacia* Δ*ogcI*Δ*ogcX* Tn7-*ogcI* strains carrying sfGFP or *ogcX* plasmids. Induction results in the detection of OgcX within *B. cenocepacia* Δ*ogcI*Δ*ogcX* Tn7-*ogcI* and the restoration of glycosylation (n=4).

### Proteomic analysis reveals loss of *ogcX* or *ogcA* drives alterations within the membrane proteome

Our initial proteomic characterisations (**Figure 1D, 3A, 3F**) revealed variability in tracking proteins of the OGC; thus, to improve the depth and reproducibility of protein quantification of these proteins, we turned to the use of Data-Independent Acquisition (DIA) based proteomics ^102,103^. To understand the proteomic changes uniquely driven by loss of *ogcX* and *ogcA* we compared the proteome of four strains Δ*ogcI* Tn7-*ogcI,* Δ*ogcI*Δ*ogcA* Tn7-*ogcI,* Δ*ogcI*Δ*ogcB* Tn7-*ogcI and* Δ*ogcI*Δ*ogcX* Tn7-*ogcI* under both rhamnose induced and non-induced conditions. Using DIA proteomic analysis, all five proteins of the OGC were observed allowing the confirmation of the expected loss of OGC proteins (**Supplementary Figure 7, Supplementary Table 13**). Across strains, a total of 3748 proteins were identified, with ∼90% identified with >2 unique peptides in each biological replicate (**Supplementary Figure 8A**). Principal component analysis revealed distinct clustering for each biological group, with the Δ*ogcI*Δ*ogcA* Tn7-*ogcI* and Δ*ogcI*Δ*ogcX* Tn7-*ogcI* strains separating along PC1 upon induction supporting the loss of these genes leading to similar overall proteomic effects when glycosylation is initiated (**Figure 4A**). Examination of the protein alterations between non-induced and induced conditions revealed that induction resulted in significant changes (defined as >2-fold difference and-log10(p-value) > 2) within strains lacking *ogcX* and *ogcA* while relatively few alterations were observed within Δ*ogcI*Δ*ogcB* Tn7-*ogcI* and Δ*ogcI* Tn7-*ogcI* (**Figure 4B**). Across the alterations observed in Δ*ogcI*Δ*ogcX* Tn7-*ogcI* and Δ*ogcI*Δ*ogcA* Tn7-*ogcI*, 722 proteins were differentially impacted in both, with Δ*ogcI*Δ*ogcX* Tn7-*ogcI* exhibiting the highest number of overall protein alterations (**Figure 4C**). Notably, the protein alterations upon induction within Δ*ogcI*Δ*ogcX* Tn7-*ogcI* and Δ*ogcI*Δ*ogcA* Tn7-*ogcI* were highly similar, demonstrating significant co-enrichment (Fisher exact test Benjamini-Hochberg corrected *p*-value = 8.20*10^-204^, **Supplementary Table 14 and Supplementary Figure 8B**) with these alterations also being highly correlative (Pearson correlation: 0.81, **Figure 4E**). To assess the overall patterns within the observed protein alterations, Gene Ontology (GO) enrichment was undertaken on the proteome alterations observed within Δ*ogcI*Δ*ogcX* Tn7-*ogcI,* revealing an over-representation of membrane-associated protein classes (**Figure 4D, Supplementary Table 14**). This over-representation corresponds to the increased abundance of the observable membrane proteome in both Δ*ogc*IΔ*ogc*X Tn7-ogcI and Δ*ogc*IΔ*ogc*A Tn7-ogcI upon induction (**Figure 4E**). Examination of the increased membrane proteins (**Figure 4F**) confirms the induction of OgcI (BCAL3118), as well as increases in functionally diverse membrane proteins, including BCAM1996 (a homolog of the cell envelope integrity protein CreD), transporters such as BCAL1079/BCAL1081 and BCAL1511/BCAL1512, outer membrane efflux proteins such as BCAL3514, BCAS0014, and BCAL1456, as well as membrane proteins of unknown functions, including BCAL2191 and BCAM1247. It should be noted that the proteomic alterations observed during induction within Δ*ogc*IΔ*ogc*X Tn7-ogcI and Δ*ogc*IΔ*ogc*A Tn7-ogcI demonstrate a partial overlap within proteins previously reported to be impacted by the loss of glycosylation (**Supplementary Figure 9**) ^63^. Combined, DIA proteomic analysis supports disruptions within *ogc*X and *ogc*A leads to widespread proteomic alterations, characterized by a dramatic increase in the membrane proteome when glycosylation is initiated.

**Figure 4.**
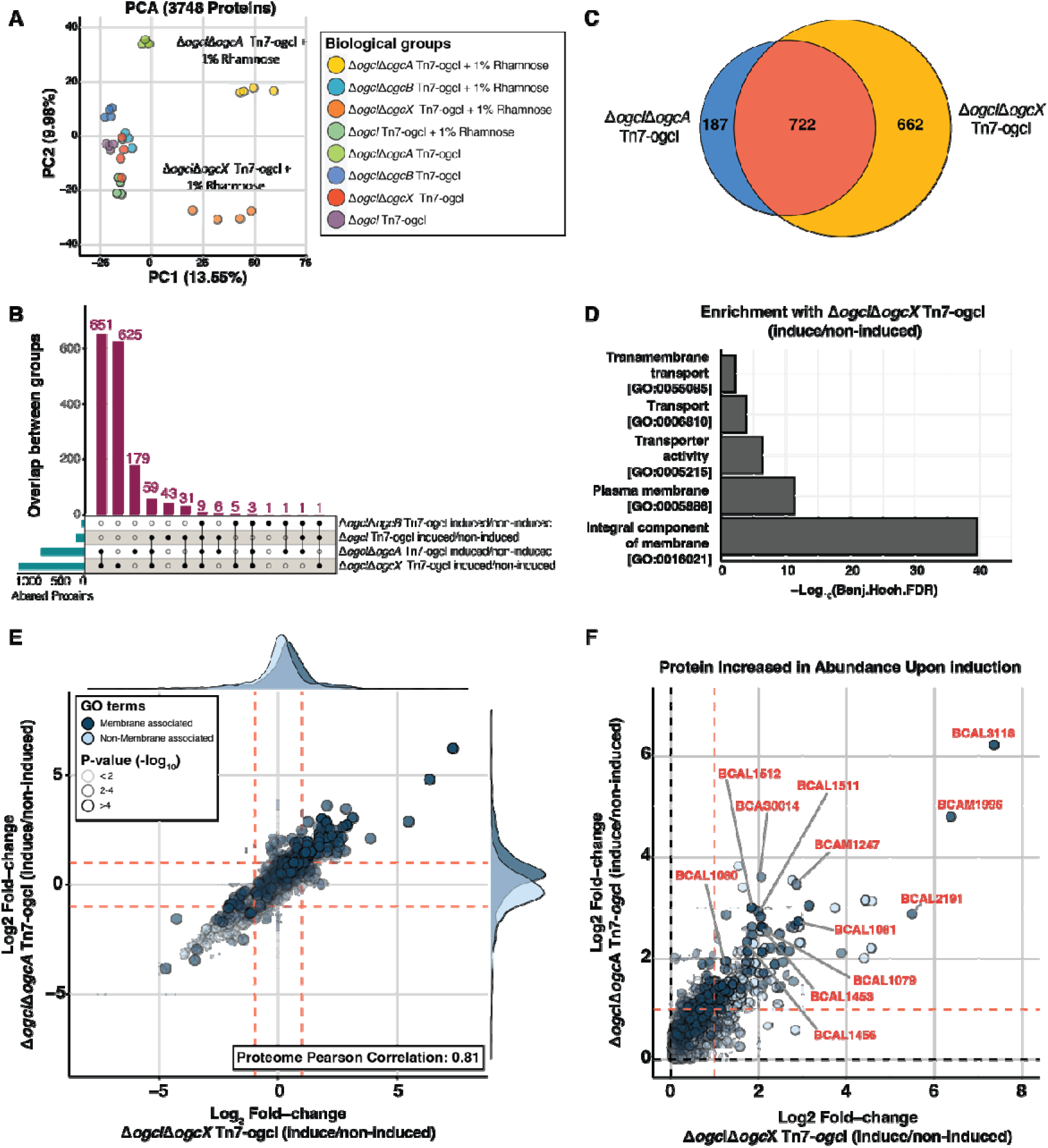
Proteomic analysis of strains in response to glycosylation initiation. DIA proteomic analysis of Δ*ogcI* Tn7-ogcI, Δ*ogcI*Δ*ogcA* Tn7-*ogcI*, Δ*ogcI*Δ*ogcB* Tn7-*ogcI*, and Δ*ogcI*Δ*ogcX* Tn7-*ogcI* strains with and without rhamnose induction (n=4). **A)** Principal component analysis (PCA) of Δ*ogcI* Tn7-*ogcI* strains under induced and non-induced conditions. Biological groups are observed to form distinct clusters, with Δ*ogcI*Δ*ogcA* Tn7-*ogcI* and Δ*ogcI*Δ*ogcX* Tn7-*ogcI* strains separating along PC1 upon induction. **B-C)** Upset plot and Venn diagram of proteomic alterations, defined as >2-fold-change and-log_10_(p-value) > 2 observed across strains, reveal that induction results in hundreds of protein alterations within Δ*ogcI*Δ*ogcA* Tn7-*ogcI* and Δ*ogcI*Δ*ogcX* Tn7-*ogcI* strains compared to Δ*ogcI*Δ*ogcB* Tn7-*ogcI* or Δ*ogcI* Tn7-ogcI. **D)** Enrichment analysis of GO terms associated with the altered proteins in Δ*ogcI*Δ*ogcX* Tn7-*ogcI* strain reveals an over-representation of proteomic changes in membrane-associated protein classes. **E)** 2D scatter plots comparing proteome changes observed within Δ*ogcI*Δ*ogcA* Tn7-*ogcI* and Δ*ogcI*Δ*ogcX* Tn7-*ogcI* in response to glycosylation initiation. Proteins associated with membrane GO-terms are color-coded dark blue with the opacity corresponding to the average p-values observed across both comparisons. **F)** Zoomed in 2D scatter plots of proteins observed to increase upon induction within Δ*ogcI*Δ*ogcA* Tn7-*ogcI* and Δ*ogcI*Δ*ogcX* Tn7-*ogcI* with membrane proteins of note highlighted.

### Loss of *ogcA* and *ogcX* impacts membrane integrity and permeability of *B. cenocepacia*

Given the dramatic membrane proteomic alterations in response to the loss of *ogcX* and *ogcA* upon induction, we hypothesised that these alterations suggests the loss of membrane integrity as noted during the inhibition of several Wzx-dependent glycan pathways ^4,6,7,34,104^. To assess membrane integrity, we assayed the impact of membrane stress (0.01% SDS) using spot plate assays revealing increased sensitivity of Δ*ogcI*Δ*ogcA* Tn7-*ogcI* and Δ*ogcI*Δ*ogcX* Tn7-*ogcI* upon glycosylation initiation compared to Δ*ogcI* Tn7-*ogcI,* Δ*ogcI*Δ*ogcB* Tn7-*ogcI or* Δ*ogcI*Δ*ogcAB* Tn7-*ogcI* strains (**Figure 5A**). In contrast, osmotic stress (2% NaCl) spot plate assays revealed adverse impacts on all strains in the absence of induction, yet only *ogcX* and *ogcA* deletion strains failed to show improved viability following glycosylation initiation (**Figure 5A**). Microtiter plate-based growth assays recapitulated these observations with the strains lacking *ogcX* and *ogcA* demonstrating alterations in growth kinetics in the presence of 0.01% SDS or 2% NaCl (**Supplementary Figure 10**). Consistent with the loss of viability being driven by the absence of *ogcX* and *ogcA* complementation ameliorated the growth defects in the presence of 0.01% SDS or 2% NaCl (**Figure 5B & C)**. These observations support the loss of membrane integrity within Δ*ogcX* and Δ*ogcA* backgrounds yet to directly assess membrane permeability in response to glycosylation initiation, we undertook Hoechst 33342 and N-phenyl-1-naphthylamine (NPN) fluorescent-based permeability assays. Utilising Hoechst 33342, a fluorescent DNA binding dye, we evaluated DNA binding revealing enhanced fluorescence within Δ*ogcX* and Δ*ogcA* backgrounds upon glycosylation initiation, yet minimal alterations in Δ*ogcI* Tn7-*ogcI,* Δ*ogcI*Δ*ogcB* Tn7-*ogcI or* Δ*ogcI*Δ*ogcAB* Tn7-*ogcI* (**Figure 5D**). Similarly, NPN uptake assays, which monitor dye partitioning into membranes ^83^, revealed increased fluorescence in Δ*ogcI*Δ*ogcX* Tn7-*ogcI* and Δ*ogcI*Δ*ogcA* Tn7-*ogcI* upon induction, supporting increased membrane permeability (**Figure 5E**, **Supplementary Figure 11**) with complementation of Δ*ogcI*Δ*ogcX* Tn7-*ogcI* and Δ*ogcI*Δ*ogcA* Tn7-*ogcI* reversing membrane permeability (**Supplementary Figure 12**). Combined, these findings support the absence of *ogcX* and *ogcA* upon glycosylation initiation compromises the *B. cenocepacia* envelope, leading to sensitisation to stressors and membrane permeability.

**Figure 5.**
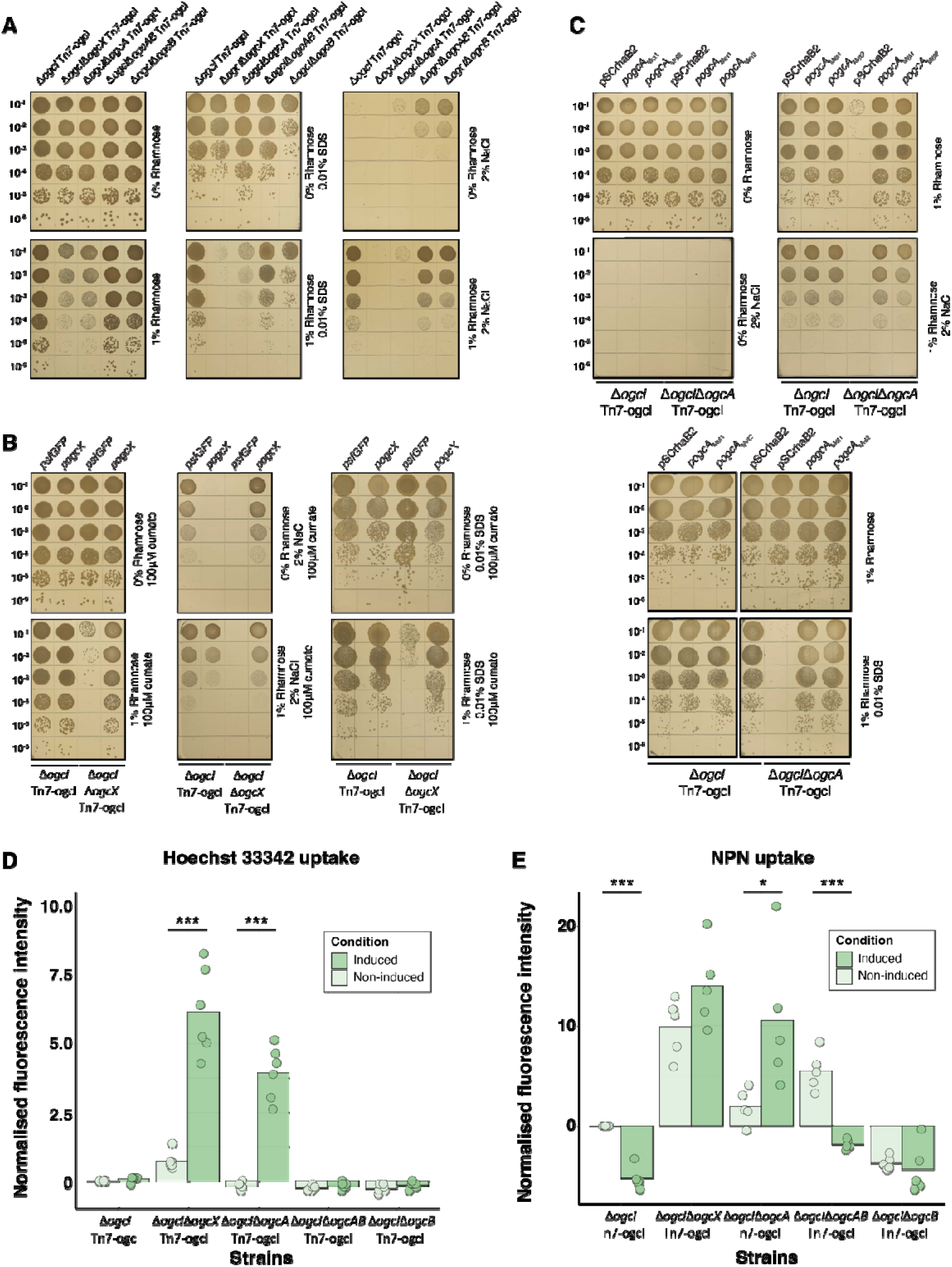
Loss of *ogcA* and *ogcX* impacts the membrane integrity of *B. cenocepacia*. **A)** Spot plate assays of strains in response to membrane (0.01% SDS) and osmotic (2% NaCl) stress with and without 1% rhamnose induction (n=4). The deletion of ogcX and *ogcA* increases sensitivity to 0.01% SDS when glycosylation is initiated, while both mutants appear sensitive to osmotic stress (2% NaCl) regardless of induction. **B-C)** Spot plate assays of complemented Δ*ogc*IΔ*ogc*A Tn7-*ogc*I and Δ*ogc*IΔ*ogc*X Tn7-*ogc*I strains in response to membrane (0.01% SDS) and osmotic (2% NaCl) stress with and without 1% rhamnose induction demonstrating the restoration of growth within complemented strains (n=4). **D-E)** Hoechst 33342 and NPN uptake assays of strains reveal loss of *ogcA* and *ogcX* leads to increased fluorescence upon induction, supporting enhanced membrane permeability (n=6 for Hoechst 33342, n=5 for NPN). The fluorescence intensities have been normalised against bacterial viability counts (**Supplementary Figure 11**).

### Loss of *ogcA* and *ogcX* enhances sensitivity of *B. cenocepacia* to antimicrobials

The observations that the loss of *ogcX* and *ogcA* leads to widespread impacts on the *B. cenocepacia* membrane highlight the potential for these effects to be exploited to potentiate antimicrobial agents. In line with this, recent transposon studies have suggested that mutations within the *ogc* alter susceptibility to β-lactam antibiotics; however, validation of these effects using CRISPRi resulted in only modest recapitulation of sensitization ^61^. Given the growth defects observed in response to induction under plasmid selection conditions (**Figure 3A, 3D and Supplementary Figure 6**), this suggests the enhanced sensitivity of strains lacking *ogcX* and *ogcA* to trimethoprim. Thus, we assessed the impact of antimicrobial agents including trimethoprim, tetracycline, ceftazidime, chlorhexidine and rifampicin on glycosylation inducible strains. Utilising broth dilution assays, changes in antimicrobial sensitivity were observed in strains lacking *ogcX* and *ogcA* upon induction yet were modest in magnitude corresponding to a 1-to 2-fold decrease in the minimal inhibitory concentrations (MIC) of trimethoprim and ceftazidime (**Supplementary Figure 13 and Supplementary Table 5**). To improve the detection of alterations in antimicrobial sensitivity, we assessed the impact of antimicrobial agents at or below the broth dilution-defined MICs using spot assays. Utilising this approach, upon induction, strains lacking *ogcX* and *ogcA* were observed to show enhanced sensitivity to tetracycline, rifampicin, trimethoprim and ceftazidime compared to Δ*ogcI*Δ*ogcB* Tn7-*ogcI* and Δ*ogcI* Tn7-*ogcI* (**Figure 6**). Interestingly, for both trimethoprim and ceftazidime, all strains showed reduced growth without induction, yet the growth of Δ*ogcI*Δ*ogcB* Tn7-*ogcI* and Δ*ogcI* Tn7-*ogcI* was improved upon induction (**Figure 6**). Combined, these findings support the loss of *OGC* genes sensitise *B. cenocepacia* to several antimicrobial agents with these impacts most pronounced for strains lacking *ogcX* or *ogcA* upon initiation of glycosylation.

**Figure 6.**
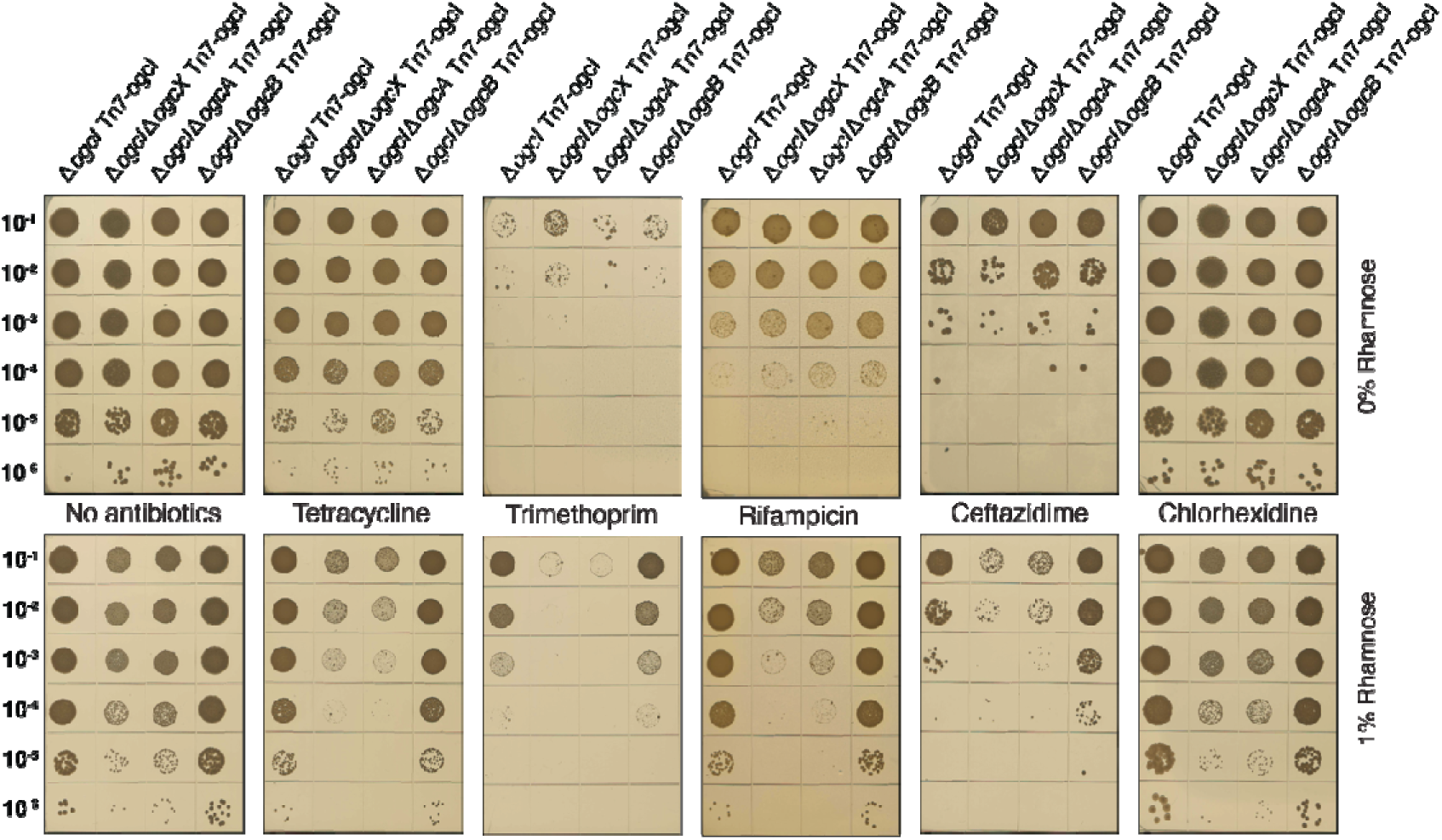
Loss of *ogcA* and *ogcX* sensitises *B. cenocepacia* to antimicrobials. Spot plate assays of *B. cenocepacia* strains, including Δ*ogcI* Tn7-*ogcI*, Δ*ogcI*Δ*ogcX* Tn7-*ogcI*, Δ*ogcI*Δ*ogcA* Tn7-*ogcI*, and Δ*ogcI*Δ*ogcB* Tn7-*ogcI* on CAMHA containing various antibiotics, with or without induction with 1% rhamnose. Rhamnose-induced glycosylation resulted in increased susceptibility to tetracycline, rifampicin, trimethoprim and ceftazidime in strains lacking *ogcA* and *ogcX*. Data representative of four biological replicates per strain per antimicrobial.

### Expression of UppS and OgcX improves the viability of *B. cenocepacia* in response to *ogc* biosynthesis blockages

As the deletion of *ogcA* and *ogcX* leads to deleterious effects upon glycosylation initiation, we sought to further support the role of Und-P/Und-PP sequestration in driving these effects. We reasoned that modulating Und-P precursor levels or enhancing glycan intermediate translocation would counteract Und-P/Und-PP sequestration. To probe this, we overexpressed undecaprenyl pyrophosphate synthase (UppS), the *de novo* synthase responsible for generating Und-P, which has been previously shown to relieve Und-P sequestration (**Figure 7A**) ^7,105^. Utilising cumate-inducible vectors containing the two putative annotated UppS enzymes of *B. cenocepacia* K56-2, BCAL2087 and BCAM2067 within the vectors p*BCAL2087* and p*BCAM2067* respectively, we assessed the impact of UppS overexpression in the absence of *ogcA* and *ogcX* with glycosylation initiation. Within Δ*ogcI*Δ*ogcA* Tn7-*ogcI* and Δ*ogcI*Δ*ogcX* Tn7-*ogcI*, the induction of p*BCAL2087* partially alleviated reductions observed within Δ*ogcA* and Δ*ogcX*, while p*BCAM2067* only restored viability within Δ*ogcI*Δ*ogcA* Tn7-*ogcI* (**Figure 7B**). These observations support the partial repression of growth defects upon induction of UppS. To assess if enhancing glycan translocation restores growth, we introduced *pogcX* into Δ*ogcI*Δ*ogcA* Tn7-*ogcI* which improved growth upon the initiation of glycosylation (**Figure 7C**). To probe the impact of OgcX overexpression on membrane integrity spot assays were undertaken revealing the amelioration of growth defects within Δ*ogcI*Δ*ogcA* Tn7-*ogcI* compared to the control vector p*sfGFP* (**Figure 7D/E**). Surprisingly, while *pogcX* improved resistance to membrane stresses in Δ*ogcI*Δ*ogcA* Tn7-*ogcI* in response to 0.01% SDS (**Figure 7D**) we noted reduced viability within Δ*ogcI*Δ*ogcB* Tn7-*ogcI* in response to salt and SDS stress compared to the control vector p*sfGFP* (**Figure 7D/E**). To confirm the overexpression of OgcX allowed the translocation of incomplete *O*-linked glycans we undertook glycoproteomic analysis confirming both enhanced glycosylation within inducible strains and the appearance of truncated glycans within Δ*ogcI*Δ*ogcA* Tn7-*ogcI* and Δ*ogcI*Δ*ogcB* Tn7-*ogcI* (**Figure 7F, Supplementary Table 15 and 16**). Manual inspection of glycopeptide assignments within Δ*ogcI*Δ*ogcA* Tn7-*ogcI* confirms the identity of these glycosylation events as corresponding to the modification of peptides with HexNAc_2_ glycans (**Figure 7G**). Of note, while glycopeptide analysis suggested the presence of both HexNAc_2_ and HexNAc modified peptides within Δ*ogcI*Δ*ogcB* Tn7-*ogcI* (**Figure 7F**) manual inspection of glycopeptide spectra demonstrate the miss assignment of glycopeptides possessing multiple individual glycosylation events as HexNAc_2_ events (**Supplementary Figure 14**). Combined, these data demonstrate that both modulating UppS activity and enhancing glycan translocation of HexNAc_2,_ but not HexNAc, alleviates the growth defects associated with initiating glycosylation in strains defective in completing the Burkholderia O-linked glycan.

**Figure 7.**
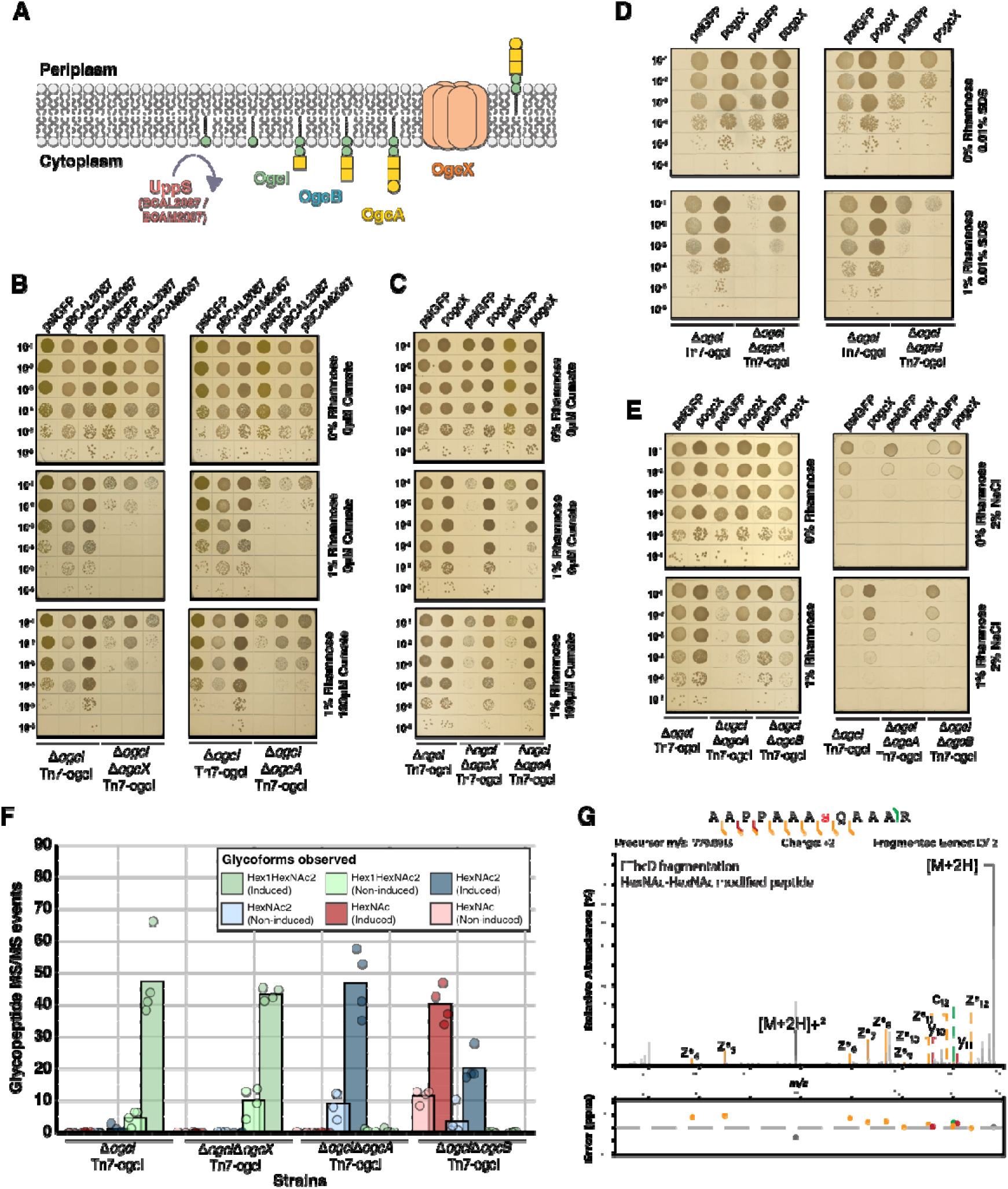
Overexpression of UppS or OgcX modulates *ogc* associated defects in *B. cenocepacia*. **A)** Proposed enzymes responsible for the *de novo* synthesis of the undecaprenyl pool and steps inferred within *ogc* biosynthesis. Within *B. cenocepacia*, two putative undecaprenyl pyrophosphate synthases (UppS) are assigned corresponding to BCAL2087 and BCAM2067, while OgcI is responsible for *OGC* initiation and OgcB the addition of the second GalNAc. The functions of OgcA, as the glycotransferase responsible for the addition of the final Gal and OgcX, as the flippase responsible for glycan translocation have only been inferred to date. **B)** Spot plate assays of the *B. cenocepacia* Δ*ogcI* Tn7-*ogcI*, Δ*ogcI*Δ*ogcX* Tn7-*ogcI* and Δ*ogcI*Δ*ogcA* Tn7-*ogcI* containing pcumate-sfGFP, pcumate-BCAL2087 or pcumate-BCAM2067 reveal the initiation of glycosylation reduces viability while induction of putative UppS partially restores the defect observed within Δ*ogcA* and Δ*ogcX* strains (n=4). **C)** Spot plate assays of the *B. cenocepacia* Δ*ogcI* Tn7-*ogcI*, Δ*ogcI*Δ*ogcX* Tn7-*ogcI* and Δ*ogcI*Δ*ogcA* Tn7-*ogcI* containing pcumate-sfGFP (*psfGFP*) and pcumate-ogcX (*pogcX*) demonstrating the complementation of Δ*ogcI*Δ*ogcA* Tn7-*ogcI* with *pogcX* restores growth (n=4). **D and E)** Spot plate assays of the *B. cenocepacia* Δ*ogcI* Tn7-*ogcI*, Δ*ogcI*Δ*ogcA* Tn7-*ogcI* and Δ*ogcI*Δ*ogcB* Tn7-*ogcI* containing pcumate-sfGFP (*psfGFP*) and pcumate-ogcX (*pogcX*) in response to membrane (0.01% SDS) and osmotic (2% NaCl) stresses with glycosylation initiation demonstrating that p*ogcX* restores resistant to stress in Δ*ogcI*Δ*ogcA* Tn7-*ogcI* yet does not suppress the defect observed in Δ*ogcI*Δ*ogcB* Tn7-*ogcI* (n=4). **F)** Quantification of glycopeptides from whole-cell proteomic analysis of *B. cenocepacia* Tn7-*ogcI* strains carrying p*ogcX*. Induction results in the detection of glycosylation within strains with varying glycoforms observed across strains (n=4). **G)** EThcD fragmentation of the HexNAc2 modified peptide ^397^AAPPAAASQAAAR^409^ (BCAL1674 Uniprot: B4E8U6) confirming the localization of the HexNAc_2_ glycosylation event residue S^404^.

### Modulation of early steps in *ogc* biosynthesis are also detrimental to *B. cenocepacia* viability

Finally, as changes driven by *ogc* perturbations were detrimental we questioned if shifting the equilibrium of the Und-P/Und-PP pool associated with the *ogc* alone would also impact viability. To probe this, we overexpressed components of the *ogc* within the parental *B. cenocepacia* K56-2 background and assess viability utilising spot assays. Introduction of rhamnose inducible vectors containing *ogcI* (p*ogcI*), *ogcB* (p*ogcB*), *ogcA* (p*ogcA*_Met1_) as well as the vector p*ogcAB*_Met1_*, which* allows the expression of both *ogcA* and *ogcB,* revealed induction of p*ogcI* and p*ogcB* resulted in reduced viability, while the overexpression of *ogcA*(p*ogcA*), *ogcAB* (p*ogcAB*), and the complete *ogc* (p*ogc*) showed no impact on viability (**Figure 8**). Critically, complementation of *B. cenocepacia* Δ*ogcAB,* Δ*ogcB* and Δ*ogcI* with p*ogcAB*_Met1_, p*ogcB* and p*ogcI* induced with 0.05% rhamnose restored glycosylation, confirming the functionality of these constructs (**Supplementary Figure 15 to 17, Supplementary Table 17 to 22**). These observations support that the overexpression of early steps in *O*-linked glycan biosynthesis, OgcI and OgcB, appear to impact viability, yet these effects can be alleviated in the case of *ogcB* by co-expression of *ogcA*. Combined, these observations support alterations in early steps in *O*-linked glycan biosynthesis modulate *ogc*-associated viability supporting the sensitivity of the Und-P/Und-PP pool to sequestration from non-physiological expression levels of *ogc* proteins.

**Figure 8.**
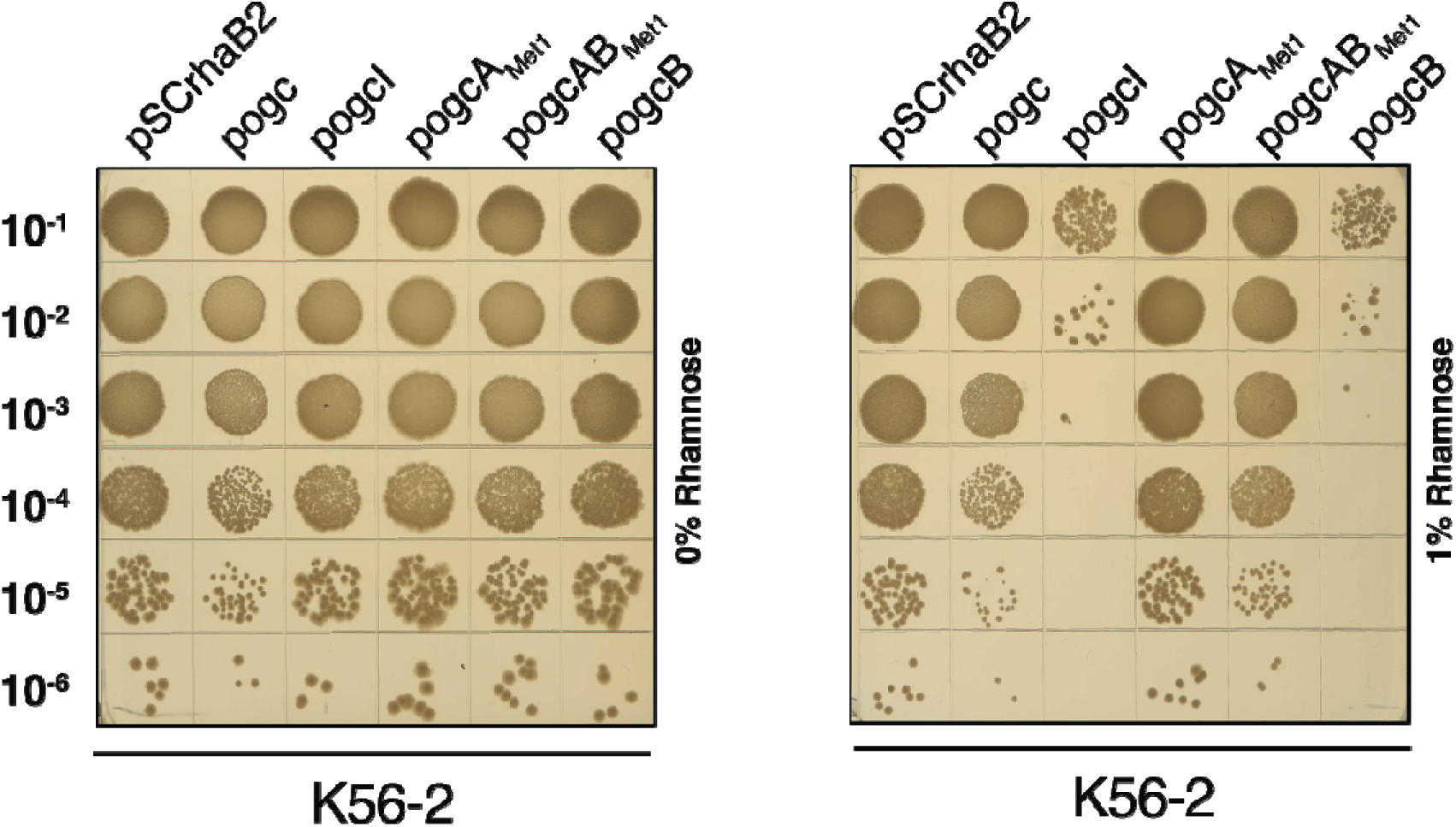
Modulation of early steps in *ogc* biosynthesis impacts *B. cenocepacia* viability. Spot plate assays of *B. cenocepacia* K56-2 (WT) containing the expression vectors pSCrhaB2, pSCrhaB2-*ogc (* p*ogc)*, pSCrhaB2-*ogcI (* p*ogcI)*, pSCrhaB2-*ogcA-Met1 (* p*ogcA*_Met1_), pSCrhaB2-*ogcB* (p*ogcB*), pSCrhaB2-*ogcAB-Met1* (p*ogcAB*_Met1_). Induction of *ogc*I and *ogc*B results in reduced viability, while the expression of *ogc*A and *ogcA B*demonstrated no reduction in viability (n=4).

## Discussion

*Burkholderia O*-linked glycosylation, mediated by the *ogc* and *pglL* ^47,59^, are highly conserved at the pathway and glycan composition levels ^45–47^. While previous work demonstrated the requirement of the *ogc* for the generation of the trisaccharide used for glycosylation ^47^, a deeper understanding of how glycan fidelity is controlled, as well as the impact of *ogcA* and *ogcX* on *B. cenocepacia* has not been assessed. In line with the growing recognition of the impact of Und-P/ Und-PP sequestration by dead-end intermediates of O-antigens ^6,11^, capsule ^35^, cell wall teichoic acids ^106,107^ and the ECA ^7,10^ glycans, we demostrate that blocking *ogc* also lead to sequestration of the Und-P/ Und-PP pool. Critically, this work highlights that in line with previous studies on Wzx-dependent glycan pathways ^3,4,10,27^, the *B. cenocepacia O*-linked glycosylation pathway appears highly selective for its complete cognate glycan. This observation sheds light on the near-exclusive presence of the observed Gal–GalNAc_2_ structure decorating glycoproteins within all *Burkholderia* glycoproteomic studies to date ^46,47,59,88,108,109^ and the mechanism responsible for glycan selectivity.

The conservation of the *Burkholderia O*-linked glycan makes this glycosylation system unique among characterised bacterial protein glycosylation systems to date, including the *O*-linked glycosylation systems of *Acinetobacter* ^110–112^ and *Neisseria* ^40,58,113^, as well as the *N*-linked glycosylation systems of *Campylobacter* ^38,39^ where extensive glycan diversity has been observed. While the conservation of glycans in the context of protein glycosylation systems appears atypical, the conservation of glycan biosynthesis across species, or even genera, has previously been noted, as in the case of the ECA ^114,115^. For the ECA, it has been suggested that the invariance of this oligosaccharide underpins its importance to Enterobacterales physiology ^115^. While the function of *Burkholderia O*-linked glycosylation is still unknown, this work raises a critical question on the importance of the cognate glycan in glycosylation. Despite the identification of *B. cenocepacia* protein glycosylation over a decade ago, we have yet to identify a specific function for glycosylation, nor have individual glycosylation events been identified that require glycosylation for functionality ^59,63,109^. As demonstrated for other Wzx, non-native substrates can be translocated under heightened levels of flippase expression ^14–16^ and consistent with this we demonstrate that overexpression of OgcX enhances the translocation of non-cognate *ogc* structures (**Figure 7F/G**). These overexpression studies support the role of OgcX as a flippase and confirms the function of OgcA as the glycosyltransferase responsible for adding the final Gal residue of the *O*-linked trisaccharide. Importantly this ability to enhance translocation within Δ*ogcI*Δ*ogcA* Tn7-*ogcI* allows us to demonstrate that poor translocation efficiency appears to underpin the detrimental effects observed from the loss of OgcA (**Figure 7B-E**). However, we observe that translocation of a single HexNAc, as seen in the case of Δ*ogc*IΔ*ogc*B Tn7-*ogcI* appears to be detrimental to *B. cenocepacia* in response to salt and SDS stressors (**Figure 7D/E**) suggesting that glycosylation with HexNAc alone may be insufficient for optimal *B. cenocepacia* physiology in response to membrane stressors. Thus, this work provides experimental confirmation of the roles of OgcX as well as OgcA, highlights how *O*-linked glycan fidelity is controlled and supports the importance of glycosylation with specific glycan structures within *B. cenocepacia* for membrane stress resistance.

Our observations of deleterious effects in response to the loss of *ogcX* (**Figure 1, 2 & 3**) suggest the previously reported Δ*ogcX* isolate within *B. cenocepacia* ^47^ likely possessed an uncharacterised suppressor mutation. As noted previously, suppressor mutations can rapidly emerge upon Und-P/Und-PP sequestration ^9,33,34^ with the emergence of a potential suppressor mutation highlighted as the potential source of confounding results associated with defining the function of the WzxE flippase within *E. coli* K-12 ^10^. While we were unable to obtain the previously reported Δ*ogcX* isolate the observation that low-level glycosylation of a truncated glycan of a single HexNAc residue was observed within this strain ^47^, suggests a potential disruption within the *O-*linked glycosylation system. In contrast to previous results, our observation of the requirement of OgcX for *O*-glycosylation supports that this flippase is indispensable within *B. cenocepacia* for translocation of the *O*-linked glycan into the periplasmic space. Interestingly, not all flippases are indispensable as has been noted within *E. coli* K-12, where some redundancy exists in the ability of ECA to be mobilised to the periplasm by the O-antigen (Wzx_O16_) or colanic acid (WzxC) flippases ^116^. Our analysis of glycosylation in the absence of *ogcX* (**Figure 2D**) is consistent with a lack of functional redundancy for the cognate *O*-linked glycan by the remaining Wzx flippases of *B. cenocepacia.* Within *B. cenocepacia* K56-2, previous analysis has highlighted the presence of at least five putative Wzx flippases with similar functional indispensability observed for the peptidoglycan flippase murJ_Bc_ (BCAL2764) ^117^. Thus, these findings support that OgcX is the sole flippase required for translocation of the *O*-linked glycan, and alternative flippases are not functionally redundant for this glycan in *B. cenocepacia*.

We have previously reported the inability to generate Δ*ogcA* within *B. cenocepacia* ^47^, yet disruptions of *ogcA* were recently suggested to be tolerated ^61^. Our data assessing the loss of *ogcA* using plasmid-based complementation (**Figure 1**) as well as strains with inducible control of glycosylation (**Figure 2 to 7**) supports that, akin to *ogcX*, the loss of *ogcA* is deleterious within *B. cenocepacia*. Previously, putative Δ*ogcA* candidates were observed within high-density transposon libraries yet *ogcA* CRISPRi silencing failed to recapitulate the resulting phenotypes associated with Δ*ogcA* ^61^. As such, we suggest that the Δ*ogcA* reported within prior work may contain unexpected secondary mutations or that the presence of transposons within *ogcA* may have adversely affected the expression of neighbouring genes of the *ogc*. While transposon libraries coupled with sequencing provide a robust means to assess gene function at a genomic scale ^118^, it is recognised that this approach can lead to inactivation or polar effects in neighbouring genes ^119,120^. Furthermore, the enrichment of transposon junctions prior to sequencing ^118^ does not allow the assessment of unlinked potential suppressor or inactivation events, which may restore variability even in the presence of insertions into conditionally essential genes. Of note within our work, the utilisation of proteomics allowed the assessment of *ogcA*, ogc*X,* and ogc*B* mutagenesis, revealing neighbouring *ogc* proteins are unaffected by the removal of these genes (**Supplementary Figure 7**). Considering our findings, in addition to prior transposons studies of *B. cenocepacia* and *Burkholderia pseudomallei* that assign *ogcA* as essential ^60,62^, this supports the assignment of *ogcA* as conditionally essential.

The observation that both *ogcA* and *ogcX* appear conditionally essential is consistent with a growing body of work highlighting the detrimental impacts of blocking Und-P/PP biosynthetic pathways ^6,7,11^. As with previous studies on the deleterious impacts of Und-P/PP sequestration, the use of inducible systems ^10,11,14^ provided a tractable way to explore the roles of *ogcA*/*ogcX* while limiting potential confounding effects from the emergence of suppressors. The inducible deleterious effects from the loss of *ogcA* and *ogcX* (**Figure 2**) support fouling of the Und-P/PP pool with complementation confirming the restoration of glycosylation as well as suppression of growth defects by the re-introduction of *ogcX* / *ogcA* (**Figure 3 an d).5**Interestingly, within this work, we noted that while the introduction of plasmid variants lacking *ogcA/X* were unable to be recovered from *B. cenocepacia* Δ*ogc* rhamnose inducible strains lacking *ogcA/X* were viable although exhibited reduced growth rates (**Figure 2, Supplementary Figure 4**). While the viability of inducible Δ*ogc*IΔ*ogc*A Tn7-ogcI *and* Δ*ogc*IΔ*ogc*X Tn7-ogcI strains was essential for allowing proteomic (**Figure 4**) and phenotypic assessments (**Figure 5, 6 and 7**), this does suggest the overall restoration of glycosylation mediated by Tn7-rha-*ogcI* may be reduced compared to wild type *B. cenocepacia,* reducing the magnitude of the effect of sequestration on the Und-P/PP pool. In line with this, we noted higher overall glycosylation levels, as determined by identified glycopeptides, within wildtype *B. cenocepacia* (**Figure 1C**) compared to the observed levels within glycosylation inducible strains (**Figure 2D, 3C, 3F and 7**) **D**consistent with an overall reduced glycosylation capacity. Regardless, the ability to demonstrate changes in membrane permeability and stress sensitivity (**Figures 5 and 6**) highlights the importance of *O*-linked glycan completion for *B. cenocepacia* physiology as well as allowed the assessment of antibiotic sensitisation previously suggested for Δ*ogcA* ^61^. Our work demonstrates that upon initiation of glycosylation both Δ*ogcA and* Δ*ogcX* become sensitised to clinically important antibiotics including tetracycline, rifampicin, trimethoprim as well as ceftazidime (**Figure 6**). This enhanced sensitivity also reconciles the differences we observe between experiments undertaken using plasmids (**Figure 1F, 3A, 3D, 5B, 5C, 7B, 7D and Supplementary Figure 6**) compared to inducible strains alone (**Figure 2B and 5A**) supporting that the antibiotic selection required to maintain plasmids exacerbates the observed viability defect within Δ*ogcA and* Δ*ogcX strains*. It should be noted that the observation that inhibiting *ogcA* or *ogcX* can modulate antibiotic sensitisation suggests targeting the *ogc* may be a potent way to sensitise *Burkholderia* to antibiotics. The concept of modulating the Und-P/PP pool to sensitise *B. cenocepacia* has been previously explored by the depletion of *dxr* (BCAL2085) ^121^, which directly contributes to the precursors required for UppS-mediated *de novo* synthesis of Und-P ^1,122^. Thus, fouling of the Und-P/PP pool through inhibition of OgcX or OgcA may be an attractive approach for drug re-sensitisation.

Previous studies have noted that the inhibition of peptidoglycan biosynthesis is likely the key defect responsible for the deleterious effects of Und-P/PP fouling ^6,7,105^. Consistent with this our DIA proteomic analysis reveals an increase in several proteins associated with peptidoglycan upon initiation of glycosylation within Δ*ogcA/X* strains including BCAL0310 (*murA*), BCAL3412 (*mtgA*), BCAL3463 (*fts*W) and BCAL2777 *a* putative N-acetylmuramoyl-L-alanine amidase (**Supplementary Table 13, Supplementary Figure 18**). Interestingly, among the altered membrane proteins, one of the most significant changes observed within Δ*ogcA* and Δ*ogcX* corresponds to a >30-fold increase in BCAM1996 (CreD, **Figure 4F**). Within *Pseudomonas aeruginosa* CreD is known to be associated with sensing changes in peptidoglycan synthesis being potently transcribed in response to perturbations in peptidoglycan binding proteins ^123^ and suggested to be important for peptidoglycan Und-P/Und-PP recycling ^124^. Similarly, within *Stenotrophomonas maltophilia* CreD has also been shown to be critical for envelope integrity and maintaining cell morphology ^125^. The increased abundance of CreD in response to glycosylation initiation within Δ*ogc*IΔ*ogc*A Tn7-ogcI *and* Δ*ogc*IΔ*ogc*X Tn7-ogcI may suggest a more general role for CreD in Und-P/Und-PP recycling. Of the proteomic alterations observed it is notable that the proteome changes observed within Δ*ogc*IΔ*ogc*A Tn7-ogcI and Δ*ogc*IΔ*ogc*X Tn7-ogcI are similar yet more extensive than the proteomic changes observed from the loss of glycosylation alone (**Supplementary Figure 9**) ^63^. These observations support that blocking *ogc* biosynthesis induces extensive and pleiotropic effects, beyond the changes seen from the loss of protein glycosylation alone, including evidence of changes in peptidoglycan biosynthesis which are consistent with Und-P/PP fouling.

In this study, it is noteworthy that for the overexpression of the putative UppS enzymes only BCAL2087 was able to alleviate the growth defects associated with glycosylation initiation in both the Δ*ogcA* and Δ*ogcX* backgrounds yet BCAM2067 only partially restored the viability of Δ*ogcI*Δ*ogcA* Tn7-*ogcI* (**Figure 7B**). While the exact cause of this difference is unclear, it is important to note that the Und-P/PP pool is not a homogeneous pool of polyisoprenyl lipids ^126^. It is possible that the assigned UppS enzymes contribute different polyisoprenyl lipids, which influence the Und-P/PP pool and lead to changes in tolerance to Und-PP intermediates, however, this is speculative, and the exact cause of this difference is unknown. Consistent with changes in the Und-P/PP pool impacting viability our analysis of individual *ogc* components reveals OgcI and OgcB alone as sufficient to inhibit growth when overexpressed within *B. cenocepacia* (**Figure 8**). These proteins do not themselves impact growth within *E. coli* (**Supplementary Figure 19**) and when induced with 0.05% rhamnose restore glycosylation within *B. cenocepacia* Δ*ogcB* and Δ*ogcI* (**Supplementary Figure 16 and 17**). Thus, this reduced viability observed upon induction of OgcI and OgcB supports that increasing the proportion of early-stage intermediates in *ogc* biosynthesis leads to the sequestration of the Und-PP/P pool. In line with this model, the overexpression of OgcA with OgcB was found to relieve toxicity, supporting the importance of adequate enzyme capacities in meeting the demands during *ogc* biosynthesis. It should be noted a caveat of our model is the assumption that the products of the *ogc* solely supply glycan units used for protein glycosylation, yet whether this is the case is unknown. As noted within other glycan biosynthesis pathways both glycan intermediates as well as complete glycan units may be incorporated into different glycoconjugates ^114,115^. Previous studies of *O*-linked glycosylation systems have demonstrated that glycans used for glycosylation may be derived from several Und-P-dependent biosynthesis pathways, such as the O-antigen in *Pseudomonas aeruginosa* ^127,128^ and capsule pathways within the *Moraxellaceae* ^129,130^. In light of this, while the *ogc* is sufficient and required for protein glycosylation ^47^ and we show the modulation of *ogc* intermediates impact viability, it is currently unclear if this pathway contributes solely to protein glycosylation, which impacts the interpretation of these findings.

In conclusion, this work demonstrates that the loss of *ogcA*/*X* within *B. cenocepacia* results in detrimental impacts on viability, membrane permeability and the proteome, which appears to sensitise strains to membrane stresses including clinically useful antibiotics. This work highlights that akin to the impacts of glycan biosynthesis blockages observed in O-antigens ^6,9^ and the ECA ^7,10^, modulating or blocking the generation of *ogc* intermediates appears to lead to sequestration of the Und-P/PP pool within *B. cenocepacia.* These findings not only advance our mechanistic understanding of how glycan fidelity is maintained for Burkholderia *O*-linked protein glycosylation but the therapeutic potential of targeting steps in this pathway.

## Acknowledgements

N.E.S was supported by an Australian Research Council (ARC) Future Fellowship (FT200100270), an ARC Discovery Project Grant (DP210100362) and a National Health and Medical Research Council Ideas grant (2018980). We thank the Melbourne Mass Spectrometry and Proteomics Facility of The Bio21 Molecular Science and Biotechnology Institute for access to MS instrumentation. We thank Nicholas Coleman for sharing the plasmid pUS250-sfGFP and Erin Garcia for sharing the plasmid pFlptet.

## AUTHOR CONTRIBUTION STATEMENT

**Leila Jebeli:** Investigation, Validation, Visualization, Writing - Original Draft, Formal analysis; **Taylor A. McDaniels:** Investigation, Visualization, Formal analysis, Supervision; **Duncan T. T. Ho:** Investigation, Visualization, Formal analysis; **Hamza Tahir:** Investigation, Resources; **Nicholas L. Kai-Ming:** Investigation, Visualization; **Molli Mcgaw:** Investigation, Resources; **Kristian I. Karlic:** Software, Formal analysis, Visualization; **Jessica M. Lewis:** Supervision, Investigation, Visualization; **Nichollas E. Scott:** Conceptualization, Visualization, Investigation, Formal analysis, Supervision, Writing - Review & Editing, Funding acquisition.

## References

1 Touz, E. T. & Mengin-Lecreulx, D. Undecaprenyl Phosphate Synthesis. EcoSal Plus 3 (2008). 10.1128/ecosalplus.4.7.1.7

2 Workman, S. D. & Strynadka, N. C. J. A Slippery Scaffold: Synthesis and Recycling of the Bacterial Cell Wall Carrier Lipid. J Mol Biol 432, 4964–4982 (2020). 10.1016/j.jmb.2020.03.025

3 Hong, Y., Cunneen, M. M. & Reeves, P. R. The Wzx translocases for Salmonella enterica O-antigen processing have unexpected serotype specificity. Molecular microbiology 84, 620–630 (2012). 10.1111/j.1365-2958.2012.08048.x

4 Liu, M. A., Stent, T. L., Hong, Y. & Reeves, P. R. Inefficient translocation of a truncated O unit by a Salmonella Wzx affects both O-antigen production and cell growth. FEMS Microbiol Lett 362 (2015). 10.1093/femsle/fnv053

5 Barreteau, H. et al. Quantitative high-performance liquid chromatography analysis of the pool levels of undecaprenyl phosphate and its derivatives in bacterial membranes. J Chromatogr B Analyt Technol Biomed Life Sci 877, 213–220 (2009). 10.1016/j.jchromb.2008.12.010

6 Jorgenson, M. A. & Young, K. D. Interrupting biosynthesis of O-antigen or the lipopolysaccharide core produces morphological defects in Escherichia coli by sequestering undecaprenyl phosphate. Journal of bacteriology (2016). 10.1128/JB.00550-16

7 Jorgenson, M. A., Kannan, S., Laubacher, M. E. & Young, K. D. Dead-end intermediates in the enterobacterial common antigen pathway induce morphological defects in Escherichia coli by competing for undecaprenyl phosphate. Molecular microbiology 100, 1–14 (2016). 10.1111/mmi.13284

8 Tatar, L. D., Marolda, C. L., Polischuk, A. N., van Leeuwen, D. & Valvano, M. A. An Escherichia coli undecaprenyl-pyrophosphate phosphatase implicated in undecaprenyl phosphate recycling. Microbiology (Reading) 153, 2518–2529 (2007). 10.1099/mic.0.2007/006312-0

9 Yuasa, R., Levinthal, M. & Nikaido, H. Biosynthesis of cell wall lipopolysaccharide in mutants of Salmonella. V. A mutant of Salmonella typhimurium defective in the synthesis of cytidine diphosphoabequose. Journal of bacteriology 100, 433–444 (1969). 10.1128/jb.100.1.433-444.1969

10 Rick, P. D. et al. Evidence that the wzxE gene of Escherichia coli K-12 encodes a protein involved in the transbilayer movement of a trisaccharide-lipid intermediate in the assembly of enterobacterial common antigen. The Journal of biological chemistry 278, 16534–16542 (2003). 10.1074/jbc.M301750200

11 Qin, J., Hong, Y. & Totsika, M. Determining glycosyltransferase functional order via lethality due to accumulated O-antigen intermediates, exemplified with Shigella flexneri O-antigen biosynthesis. Applied and environmental microbiology 90, e0220323 (2024). 10.1128/aem.02203-23

12 Swoboda, J. G. et al. Discovery of a small molecule that blocks wall teichoic acid biosynthesis in Staphylococcus aureus. ACS Chem Biol 4, 875–883 (2009). 10.1021/cb900151k

13 Muscato, J. D. et al. Rapid Inhibitor Discovery by Exploiting Synthetic Lethality. J Am Chem Soc 144, 3696–3705 (2022). 10.1021/jacs.1c12697

14 Sham, L. T., Zheng, S., Yakhnina, A. A., Kruse, A. C. & Bernhardt, T. G. Loss of specificity variants of WzxC suggest that substrate recognition is coupled with transporter opening in MOP-family flippases. Molecular microbiology 109, 633–641 (2018). 10.1111/mmi.14002

15 Chua, W. Z. et al. High-Throughput Mutagenesis and Cross-Complementation Experiments Reveal Substrate Preference and Critical Residues of the Capsule Transporters in Streptococcus pneumoniae. mBio 12, e0261521 (2021). 10.1128/mBio.02615-21

16 Su, T. et al. Rewiring the pneumococcal capsule pathway for investigating glycosyltransferase specificity and genetic glycoengineering. Sci Adv 9, eadi8157 (2023). 10.1126/sciadv.adi8157

17 Ruiz, N. Lipid Flippases for Bacterial Peptidoglycan Biosynthesis. Lipid Insights 8, 21–31 (2015). 10.4137/LPI.S31783

18 Cuthbertson, L., Kos, V. & Whitfield, C. ABC transporters involved in export of cell surface glycoconjugates. Microbiol Mol Biol Rev 74, 341–362 (2010). 10.1128/MMBR.00009-10

19 Kuk, A. C. Y., Hao, A. & Lee, S. Y. Structure and Mechanism of the Lipid Flippase MurJ. Annu Rev Biochem 91, 705–729 (2022). 10.1146/annurev-biochem-040320-105145

20 Hong, Y., Liu, M. A. & Reeves, P. R. Progress in Our Understanding of Wzx Flippase for Translocation of Bacterial Membrane Lipid-Linked Oligosaccharide. Journal of bacteriology 200 (2018). 10.1128/JB.00154-17

21 Liu, D., Cole, R. A. & Reeves, P. R. An O-antigen processing function for Wzx (RfbX): a promising candidate for O-unit flippase. Journal of bacteriology 178, 2102–2107 (1996). 10.1128/jb.178.7.2102-2107.1996

22 Kenyon, J. J. & Hall, R. M. Variation in the complex carbohydrate biosynthesis loci of Acinetobacter baumannii genomes. PloS one 8, e62160 (2013). 10.1371/journal.pone.0062160

23 Paton, J. C. & Trappetti, C. Streptococcus pneumoniae Capsular Polysaccharide. Microbiol Spectr 7 (2019). 10.1128/microbiolspec.GPP3-0019-2018

24 Becker, A. Challenges and perspectives in combinatorial assembly of novel exopolysaccharide biosynthesis pathways. Front Microbiol 6, 687 (2015). 10.3389/fmicb.2015.00687

25 Feldman, M. F. et al. The activity of a putative polyisoprenol-linked sugar translocase (Wzx) involved in Escherichia coli O antigen assembly is independent of the chemical structure of the O repeat. The Journal of biological chemistry 274, 35129–35138 (1999).

26 Marolda, C. L., Vicarioli, J. & Valvano, M. A. Wzx proteins involved in biosynthesis of O antigen function in association with the first sugar of the O-specific lipopolysaccharide subunit. Microbiology (Reading) 150, 4095–4105 (2004). 10.1099/mic.0.27456-0

27 Hong, Y. & Reeves, P. R. Diversity of o-antigen repeat unit structures can account for the substantial sequence variation of wzx translocases. Journal of bacteriology 196, 1713–1722 (2014). 10.1128/JB.01323-13

28 Liu, M. A., Morris, P. & Reeves, P. R. Wzx flippases exhibiting complex O-unit preferences require a new model for Wzx-substrate interactions. Microbiologyopen 8, e00655 (2019). 10.1002/mbo3.655

29 Liu, D. & Reeves, P. R. Escherichia coli K12 regains its O antigen. Microbiology (Reading) 140 **(Pt** **1**), 49–57 (1994). 10.1099/13500872-140-1-49

30 Stevenson, G. et al. Structure of the O antigen of Escherichia coli K-12 and the sequence of its rfb gene cluster. Journal of bacteriology 176, 4144–4156 (1994). 10.1128/jb.176.13.4144-4156.1994

31 Barr, K. & Rick, P. D. Biosynthesis of enterobacterial common antigen in Escherichia coli. In vitro synthesis of lipid-linked intermediates. The Journal of biological chemistry 262, 7142–7150 (1987).

32 Barr, K., Ward, S., Meier-Dieter, U., Mayer, H. & Rick, P. D. Characterization of an Escherichia coli rff mutant defective in transfer of N-acetylmannosaminuronic acid (ManNAcA) from UDP-ManNAcA to a lipid-linked intermediate involved in enterobacterial common antigen synthesis. Journal of bacteriology 170, 228–233 (1988). 10.1128/jb.170.1.228-233.1988

33 Burrows, L. L. & Lam, J. S. Effect of wzx (rfbX) mutations on A-band and B-band lipopolysaccharide biosynthesis in Pseudomonas aeruginosa O5. Journal of bacteriology 181, 973–980 (1999). 10.1128/JB.181.3.973-980.1999

34 Xayarath, B. & Yother, J. Mutations blocking side chain assembly, polymerization, or transport of a Wzy-dependent Streptococcus pneumoniae capsule are lethal in the absence of suppressor mutations and can affect polymer transfer to the cell wall. Journal of bacteriology 189, 3369–3381 (2007). 10.1128/JB.01938-06

35 Bai, J. et al. Essential Gene Analysis in Acinetobacter baumannii by High-Density Transposon Mutagenesis and CRISPR Interference. Journal of bacteriology 203, e0056520 (2021). 10.1128/JB.00565-20

36 Koomey, M. O-linked protein glycosylation in bacteria: snapshots and current perspectives. Curr Opin Struct Biol 56, 198–203 (2019). 10.1016/j.sbi.2019.03.020

37 Nothaft, H. & Szymanski, C. M. Protein glycosylation in bacteria: sweeter than ever. Nature reviews. Microbiology 8, 765–778 (2010). 10.1038/nrmicro2383

38 Nothaft, H. et al. Diversity in the protein N-glycosylation pathways within the Campylobacter genus. Molecular & cellular proteomics: MCP 11, 1203–1219 (2012). 10.1074/mcp.M112.021519

39 Jervis, A. J. et al. Characterization of the structurally diverse N-linked glycans of Campylobacter species. Journal of bacteriology 194, 2355–2362 (2012). 10.1128/JB.00042-12

40 Hadjineophytou, C. et al. Genetic determinants of genus-Level glycan diversity in a bacterial protein glycosylation system. PLoS Genet 15, e1008532 (2019). 10.1371/journal.pgen.1008532

41 Anonsen, J. H. et al. Characterization of a Unique Tetrasaccharide and Distinct Glycoproteome in the O-Linked Protein Glycosylation System of Neisseria elongata subsp. glycolytica. Journal of bacteriology 198, 256–267 (2016). 10.1128/JB.00620-15

42 Vik, A. et al. Broad spectrum O-linked protein glycosylation in the human pathogen Neisseria gonorrhoeae. Proceedings of the National Academy of Sciences of the United States of America 106, 4447–4452 (2009). 10.1073/pnas.0809504106

43 Borud, B. et al. Genetic, structural, and antigenic analyses of glycan diversity in the O-linked protein glycosylation systems of human Neisseria species. Journal of bacteriology 192, 2816–2829 (2010). 10.1128/JB.00101-10

44 Hadjineophytou, C. et al. Sculpting the Bacterial O-Glycoproteome: Functional Analyses of Orthologous Oligosaccharyltransferases with Diverse Targeting Specificities. mBio, e0379721 (2022). 10.1128/mbio.03797-21

45 Hayes, A. J., Lewis, J. M., Davies, M. R. & Scott, N. E. Burkholderia PglL enzymes are Serine preferring oligosaccharyltransferases which target conserved proteins across the Burkholderia genus. Commun Biol 4, 1045 (2021). 10.1038/s42003-021-02588-y

46 Ahmad Izaham, A. R. & Scott, N. E. Open Database Searching Enables the Identification and Comparison of Bacterial Glycoproteomes without Defining Glycan Compositions Prior to Searching. Molecular & cellular proteomics: MCP 19, 1561–1574 (2020). 10.1074/mcp.TIR120.002100

47 Fathy Mohamed, Y., et al. A general protein O-glycosylation machinery conserved in Burkholderia species improves bacterial fitness and elicits glycan immunogenicity in humans. The Journal of biological chemistry 294, 13248–13268 (2019). 10.1074/jbc.RA119.009671

48 Iwashkiw, J. A., Vozza, N. F., Kinsella, R. L. & Feldman, M. F. Pour some sugar on it: the expanding world of bacterial protein O-linked glycosylation. Molecular microbiology 89, 14–28 (2013). 10.1111/mmi.12265

49 Nothaft, H. & Szymanski, C. M. New discoveries in bacterial N-glycosylation to expand the synthetic biology toolbox. Curr Opin Chem Biol 53, 16–24 (2019). 10.1016/j.cbpa.2019.05.032

50 Borud, B. & Koomey, M. Sweet complexity: O-linked protein glycosylation in pathogenic Neisseria. Frontiers in cellular and infection microbiology 14, 1407863 (2024). 10.3389/fcimb.2024.1407863

51 Linton, D. et al. Functional analysis of the Campylobacter jejuni N-linked protein glycosylation pathway. Molecular microbiology 55, 1695–1703 (2005). 10.1111/j.1365-2958.2005.04519.x

52 Perez, C. et al. Structure and mechanism of an active lipid-linked oligosaccharide flippase. Nature 524, 433–438 (2015). 10.1038/nature14953

53 Alaimo, C. et al. Two distinct but interchangeable mechanisms for flipping of lipid-linked oligosaccharides. EMBO J 25, 967–976 (2006). 10.1038/sj.emboj.7601024

54 Kahler, C. M. et al. Polymorphisms in pilin glycosylation Locus of Neisseria meningitidis expressing class II pili. Infection and immunity 69, 3597–3604 (2001). 10.1128/IAI.69.6.3597-3604.2001

55 Power, P. M. et al. Genetic characterization of pilin glycosylation in Neisseria meningitidis. Microbiology 146 **(Pt** **4**), 967–979 (2000). 10.1099/00221287-146-4-967

56 Aas, F. E., Vik, A., Vedde, J., Koomey, M. & Egge-Jacobsen, W. Neisseria gonorrhoeae O-linked pilin glycosylation: functional analyses define both the biosynthetic pathway and glycan structure. Molecular microbiology 65, 607–624 (2007). 10.1111/j.1365-2958.2007.05806.x

57 Johannessen, C., Koomey, M. & Borud, B. Hypomorphic glycosyltransferase alleles and recoding at contingency loci influence glycan microheterogeneity in the protein glycosylation system of Neisseria species. Journal of bacteriology 194, 5034–5043 (2012). 10.1128/JB.00950-12

58 Wang, N. et al. Allelic polymorphisms in a glycosyltransferase gene shape glycan repertoire in the O-linked protein glycosylation system of Neisseria. Glycobiology 31, 477–491 (2021). 10.1093/glycob/cwaa073

59 Lithgow, K. V. et al. A general protein O-glycosylation system within the Burkholderia cepacia complex is involved in motility and virulence. Molecular microbiology 92, 116–137 (2014). 10.1111/mmi.12540

60 Gislason, A. S., Turner, K., Domaratzki, M. & Cardona, S. T. Comparative analysis of the Burkholderia cenocepacia K56-2 essential genome reveals cell envelope functions that are uniquely required for survival in species of the genus Burkholderia. Microb Genom 3 (2017). 10.1099/mgen.0.000140

61 Hogan, A. M. et al. Profiling cell envelope-antibiotic interactions reveals vulnerabilities to beta-lactams in a multidrug-resistant bacterium. Nat Commun 14, 4815 (2023). 10.1038/s41467-023-40494-5

62 Moule, M. G. et al. Genome-wide saturation mutagenesis of Burkholderia pseudomallei K96243 predicts essential genes and novel targets for antimicrobial development. MBio 5, e00926–00913 (2014). 10.1128/mBio.00926-13

63 Oppy, C. C. et al. Loss of O-linked protein glycosylation in Burkholderia cenocepacia impairs biofilm formation, siderophore activity and alters transcriptional regulators mSphere 4, e00660–00619 (2019).

64 Cardona, S. T. & Valvano, M. A. An expression vector containing a rhamnose-inducible promoter provides tightly regulated gene expression in Burkholderia cenocepacia. Plasmid 54, 219–228 (2005). 10.1016/j.plasmid.2005.03.004

65 Lopez, C. M., Rholl, D. A., Trunck, L. A. & Schweizer, H. P. Versatile dual-technology system for markerless allele replacement in Burkholderia pseudomallei. Applied and environmental microbiology 75, 6496–6503 (2009). 10.1128/AEM.01669-09

66 Aubert, D. F., Hamad, M. A. & Valvano, M. A. A markerless deletion method for genetic manipulation of Burkholderia cenocepacia and other multidrug-resistant gram-negative bacteria. Methods in molecular biology 1197, 311–327 (2014). 10.1007/978-1-4939-1261-2_18

67 Garcia, E. C., Anderson, M. S., Hagar, J. A. & Cotter, P. A. Burkholderia BcpA mediates biofilm formation independently of interbacterial contact-dependent growth inhibition. Molecular microbiology 89, 1213–1225 (2013). 10.1111/mmi.12339

68 Flannagan, R. S., Linn, T. & Valvano, M. A. A system for the construction of targeted unmarked gene deletions in the genus Burkholderia. Environ Microbiol 10, 1652–1660 (2008). 10.1111/j.1462-2920.2008.01576.x

69 Gibson, D. G. et al. Enzymatic assembly of DNA molecules up to several hundred kilobases. Nature methods 6, 343–345 (2009). 10.1038/nmeth.1318

70 Lefebre, M. D. & Valvano, M. A. Construction and evaluation of plasmid vectors optimized for constitutive and regulated gene expression in Burkholderia cepacia complex isolates. Applied and environmental microbiology 68, 5956–5964 (2002).

71 Sambrook, J., E. F. Fritsch, and T. Maniatis. *Molecular cloning: a laboratory manual*. (Cold Spring Harbor Laboratory, Cold Spring Harbor, 1990).

72 Figurski, D. H. & Helinski, D. R. Replication of an origin-containing derivative of plasmid RK2 dependent on a plasmid function provided in trans. Proceedings of the National Academy of Sciences of the United States of America 76, 1648–1652 (1979). 10.1073/pnas.76.4.1648

73 Choi, K. H., DeShazer, D. & Schweizer, H. P. mini-Tn7 insertion in bacteria with multiple glmS-linked attTn7 sites: example Burkholderia mallei ATCC 23344. Nature protocols 1, 162–169 (2006). 10.1038/nprot.2006.25

74 Hamad, M. A., Di Lorenzo, F., Molinaro, A. & Valvano, M. A. Aminoarabinose is essential for lipopolysaccharide export and intrinsic antimicrobial peptide resistance in Burkholderia cenocepacia(dagger). Molecular microbiology 85, 962–974 (2012). 10.1111/j.1365-2958.2012.08154.x

75 Bernier, S. P., Son, S. & Surette, M. G. The Mla Pathway Plays an Essential Role in the Intrinsic Resistance of Burkholderia cepacia Complex Species to Antimicrobials and Host Innate Components. Journal of bacteriology 200 (2018). 10.1128/JB.00156-18

76 Malinverni, J. C. & Silhavy, T. J. An ABC transport system that maintains lipid asymmetry in the gram-negative outer membrane. Proceedings of the National Academy of Sciences of the United States of America 106, 8009–8014 (2009). 10.1073/pnas.0903229106

77 Mychack, A. et al. A synergistic role for two predicted inner membrane proteins of Escherichia coli in cell envelope integrity. Molecular microbiology 111, 317–337 (2019). 10.1111/mmi.14157

78. Schindelin, J. et al. Fiji: an open-source platform for biological-image analysis. Nature methods 9, 676-682 (2012). 10.1038/nmeth.2019

79 Jassem, A. N. et al. In vitro susceptibility of Burkholderia vietnamiensis to aminoglycosides. Antimicrobial agents and chemotherapy 55, 2256–2264 (2011). 10.1128/AAC.01434-10

80 Richmond, G. E., Chua, K. L. & Piddock, L. J. Efflux in Acinetobacter baumannii can be determined by measuring accumulation of H33342 (bis-benzamide). J Antimicrob Chemother 68, 1594–1600 (2013). 10.1093/jac/dkt052

81 Krishnamoorthy, G. et al. Efflux Pumps of Burkholderia thailandensis Control the Permeability Barrier of the Outer Membrane. Antimicrobial agents and chemotherapy 63 (2019). 10.1128/AAC.00956-19

82 Helander, I. M. & Mattila-Sandholm, T. Fluorometric assessment of gram-negative bacterial permeabilization. J Appl Microbiol 88, 213–219 (2000). 10.1046/j.1365-2672.2000.00971.x

83 Loh, B., Grant, C. & Hancock, R. E. Use of the fluorescent probe 1-N-phenylnaphthylamine to study the interactions of aminoglycoside antibiotics with the outer membrane of Pseudomonas aeruginosa. Antimicrobial agents and chemotherapy 26, 546–551 (1984). 10.1128/AAC.26.4.546

84 Malott, R. J., Steen-Kinnaird, B. R., Lee, T. D. & Speert, D. P. Identification of hopanoid biosynthesis genes involved in polymyxin resistance in Burkholderia multivorans. Antimicrobial agents and chemotherapy 56, 464–471 (2012). 10.1128/AAC.00602-11

85 Kulak, N. A., Pichler, G., Paron, I., Nagaraj, N. & Mann, M. Minimal, encapsulated proteomic-sample processing applied to copy-number estimation in eukaryotic cells. Nature methods 11, 319–324 (2014). 10.1038/nmeth.2834

86 Harney, D. J. et al. Proteomic Analysis of Human Plasma during Intermittent Fasting. Journal of proteome research 18, 2228–2240 (2019). 10.1021/acs.jproteome.9b00090

87 Rappsilber, J., Mann, M. & Ishihama, Y. Protocol for micro-purification, enrichment, pre-fractionation and storage of peptides for proteomics using StageTips. Nature protocols 2, 1896–1906 (2007). 10.1038/nprot.2007.261

88 Ahmad Izaham, A. R., et al. What Are We Missing by Using Hydrophilic Enrichment? Improving Bacterial Glycoproteome Coverage Using Total Proteome and FAIMS Analyses. Journal of proteome research 20, 599–612 (2021). 10.1021/acs.jproteome.0c00565

89 Saba, J., Dutta, S., Hemenway, E. & Viner, R. Increasing the productivity of glycopeptides analysis by using higher-energy collision dissociation-accurate mass-product-dependent electron transfer dissociation. Int J Proteomics 2012, 560391 (2012). 10.1155/2012/560391

90 Caval, T., Zhu, J. & Heck, A. J. R. Simply Extending the Mass Range in Electron Transfer Higher Energy Collisional Dissociation Increases Confidence in N-Glycopeptide Identification. Anal Chem 91, 10401–10406 (2019). 10.1021/acs.analchem.9b02125

91 Polasky, D. A., Geiszler, D. J., Yu, F. & Nesvizhskii, A. I. Multi-attribute Glycan Identification and FDR Control for Glycoproteomics. Molecular & cellular proteomics: MCP, 100205 (2022). 10.1016/j.mcpro.2022.100205

92 Kong, A. T., Leprevost, F. V., Avtonomov, D. M., Mellacheruvu, D. & Nesvizhskii, A. I. MSFragger: ultrafast and comprehensive peptide identification in mass spectrometry-based proteomics. Nature methods 14, 513–520 (2017). 10.1038/nmeth.4256

93 Polasky, D. A., Yu, F., Teo, G. C. & Nesvizhskii, A. I. Fast and comprehensive N-and O-glycoproteomics analysis with MSFragger-Glyco. Nature methods 17, 1125–1132 (2020). 10.1038/s41592-020-0967-9

94 Holden, M. T. et al. The genome of Burkholderia cenocepacia J2315, an epidemic pathogen of cystic fibrosis patients. Journal of bacteriology 191, 261–277 (2009). 10.1128/JB.01230-08

95 Wickham, H. ggplot2: Elegant Graphics for Data Analysis. (Springer-VerlagNew York, 2016).

96 Brademan, D. R., Riley, N. M., Kwiecien, N. W. & Coon, J. J. Interactive Peptide Spectral Annotator: A Versatile Web-based Tool for Proteomic Applications. Molecular & cellular proteomics: MCP 18, S193–S201 (2019). 10.1074/mcp.TIR118.001209

97 Varga, J. J. et al. Draft Genome Sequences of Burkholderia cenocepacia ET12 Lineage Strains K56-2 and BC7. Genome Announc 1 (2013). 10.1128/genomeA.00841-13

98 Cox, J. et al. Accurate proteome-wide label-free quantification by delayed normalization and maximal peptide ratio extraction, termed MaxLFQ. Molecular & cellular proteomics: MCP 13, 2513–2526 (2014). 10.1074/mcp.M113.031591

99 Tyanova, S. et al. The Perseus computational platform for comprehensive analysis of (prote)omics data. Nature methods 13, 731–740 (2016). 10.1038/nmeth.3901

100 Perez-Riverol, Y. et al. The PRIDE database and related tools and resources in 2019: improving support for quantification data. Nucleic Acids Res 47, D442–D450 (2019). 10.1093/nar/gky1106

101 Vizcaino, J. A. et al. 2016 update of the PRIDE database and its related tools. Nucleic Acids Res 44, D447–456 (2016). 10.1093/nar/gkv1145

102 Ludwig, C. et al. Data-independent acquisition-based SWATH-MS for quantitative proteomics: a tutorial. Mol Syst Biol 14, e8126 (2018). 10.15252/msb.20178126

103 Pino, L. K., Just, S. C., MacCoss, M. J. & Searle, B. C. Acquiring and Analyzing Data Independent Acquisition Proteomics Experiments without Spectrum Libraries. Molecular & cellular proteomics: MCP 19, 1088–1103 (2020). 10.1074/mcp.P119.001913

104 Mitchell, A. M., Srikumar, T. & Silhavy, T. J. Cyclic Enterobacterial Common Antigen Maintains the Outer Membrane Permeability Barrier of Escherichia coli in a Manner Controlled by YhdP. mBio 9 (2018). 10.1128/mBio.01321-18

105 Jorgenson, M. A. et al. Simultaneously inhibiting undecaprenyl phosphate production and peptidoglycan synthases promotes rapid lysis in Escherichia coli. Molecular microbiology 112, 233–248 (2019). 10.1111/mmi.14265

106 D’Elia, M. A. et al. Lesions in teichoic acid biosynthesis in Staphylococcus aureus lead to a lethal gain of function in the otherwise dispensable pathway. Journal of bacteriology 188, 4183–4189 (2006). 10.1128/JB.00197-06

107 D’Elia, M. A., Millar, K. E., Beveridge, T. J. & Brown, E. D. Wall teichoic acid polymers are dispensable for cell viability in Bacillus subtilis. Journal of bacteriology 188, 8313–8316 (2006). 10.1128/JB.01336-06

108 Lewis, J. M. & Scott, N. E. CRISPRi-Mediated Silencing of Burkholderia O-Linked Glycosylation Systems Enables the Depletion of Glycosylation Yet Results in Modest Proteome Impacts. Journal of proteome research 22, 1762–1778 (2023). 10.1021/acs.jproteome.2c00790

109 Lewis, J. M., Jebeli, L., Coulon, P. M. L., Lay, C. E. & Scott, N. E. Glycoproteomic and proteomic analysis of Burkholderia cenocepacia reveals glycosylation events within FliF and MotB are dispensable for motility. Microbiol Spectr 12, e0034624 (2024). 10.1128/spectrum.00346-24

110 Tkalec, K. I. et al. Glycan-Tailored Glycoproteomic Analysis Reveals Serine is the Sole Residue Subjected to O-Linked Glycosylation in Acinetobacter baumannii. Journal of proteome research 23, 2474–2494 (2024). 10.1021/acs.jproteome.4c00148

111 Scott, N. E. et al. Diversity within the O-linked protein glycosylation systems of acinetobacter species. Molecular & cellular proteomics: MCP 13, 2354–2370 (2014). 10.1074/mcp.M114.038315

112 Iwashkiw, J. A. et al. Identification of a general O-linked protein glycosylation system in Acinetobacter baumannii and its role in virulence and biofilm formation. PLoS pathogens 8, e1002758 (2012). 10.1371/journal.ppat.1002758

113 Naess, L. M. et al. Genetic, Functional, and Immunogenic Analyses of the O-Linked Protein Glycosylation System in Neisseria meningitidis Serogroup A ST-7 Isolates. Journal of bacteriology 205, e0045822 (2023). 10.1128/jb.00458-22

114 Kuhn, H. M., Meier-Dieter, U. & Mayer, H. ECA, the enterobacterial common antigen. FEMS microbiology reviews 4, 195–222 (1988). 10.1111/j.1574-6968.1988.tb02743.x

115 Rai, A. K. & Mitchell, A. M. Enterobacterial Common Antigen: Synthesis and Function of an Enigmatic Molecule. mBio 11 (2020). 10.1128/mBio.01914-20

116 Kajimura, J., Rahman, A. & Rick, P. D. Assembly of cyclic enterobacterial common antigen in Escherichia coli K-12. Journal of bacteriology 187, 6917–6927 (2005). 10.1128/JB.187.20.6917-6927.2005

117 Mohamed, Y. F. & Valvano, M. A. A Burkholderia cenocepacia MurJ (MviN) homolog is essential for cell wall peptidoglycan synthesis and bacterial viability. Glycobiology 24, 564–576 (2014). 10.1093/glycob/cwu025

118 Cain, A. K. et al. A decade of advances in transposon-insertion sequencing. Nat Rev Genet 21, 526–540 (2020). 10.1038/s41576-020-0244-x

119 Mateus, A. et al. Transcriptional and Post-Transcriptional Polar Effects in Bacterial Gene Deletion Libraries. mSystems 6, e0081321 (2021). 10.1128/mSystems.00813-21

120 Hutchison, C. A., 3rd et al. Polar Effects of Transposon Insertion into a Minimal Bacterial Genome. Journal of bacteriology 201 (2019). 10.1128/JB.00185-19

121 Sass, A., Everaert, A., Van Acker, H., Van den Driessche, F. & Coenye, T. Targeting the Nonmevalonate Pathway in Burkholderia cenocepacia Increases Susceptibility to Certain beta-Lactam Antibiotics. Antimicrobial agents and chemotherapy 62 (2018). 10.1128/AAC.02607-17

122 Zhao, L., Chang, W. C., Xiao, Y., Liu, H. W. & Liu, P. Methylerythritol phosphate pathway of isoprenoid biosynthesis. Annu Rev Biochem 82, 497–530 (2013). 10.1146/annurev-biochem-052010-100934

123 Moya, B. et al. Beta-lactam resistance response triggered by inactivation of a nonessential penicillin-binding protein. PLoS pathogens 5, e1000353 (2009). 10.1371/journal.ppat.1000353

124 Zamorano, L. et al. The Pseudomonas aeruginosa CreBC two-component system plays a major role in the response to beta-lactams, fitness, biofilm growth, and global regulation. Antimicrobial agents and chemotherapy 58, 5084–5095 (2014). 10.1128/AAC.02556-14

125 Huang, H. H. et al. Expression and Functions of CreD, an Inner Membrane Protein in Stenotrophomonas maltophilia. PloS one 10, e0145009 (2015). 10.1371/journal.pone.0145009

126 Reid, C. W., Stupak, J., Szymanski, C. M. & Li, J. Analysis of bacterial lipid-linked oligosaccharide intermediates using porous graphitic carbon liquid chromatography-electrospray ionization mass spectrometry: heterogeneity in the polyisoprenyl carrier revealed. Anal Chem 81, 8472–8478 (2009). 10.1021/ac9013622

127 Horzempa, J., Dean, C. R., Goldberg, J. B. & Castric, P. Pseudomonas aeruginosa 1244 pilin glycosylation: glycan substrate recognition. Journal of bacteriology 188, 4244–4252 (2006). 10.1128/JB.00273-06

128 Castric, P., Cassels, F. J. & Carlson, R. W. Structural characterization of the Pseudomonas aeruginosa 1244 pilin glycan. The Journal of biological chemistry 276, 26479–26485 (2001). 10.1074/jbc.M102685200

129 Lees-Miller, R. G. et al. A common pathway for O-linked protein-glycosylation and synthesis of capsule in Acinetobacter baumannii. Molecular microbiology 89, 816–830 (2013). 10.1111/mmi.12300

130 Knoot, C. J. et al. Discovery and characterization of a new class of O-linking oligosaccharyltransferases from the Moraxellaceae family. Glycobiology (2022). 10.1093/glycob/cwac070

