## Supplementary document for "The Late-Stage Steps of *Burkholderia cenocepacia* Protein *O*-Linked Glycan Biosynthesis Are Conditionally Essential"

**Table of Contents**

| **Title** | **Page** |
| --- | --- |
| **Supplementary Table 1: Strain list** | **4** |
| **Supplementary Table 2: Plasmid list** | **5** |
| **Supplementary Table 3: Primer list** | **8** |
| **Supplementary Table 4: Proteomic datasets** | **14** |
| **Supplementary Table 5: Antimicrobial Minimal inhibitory concentrations (MIC).** | **17** |
| **Supplementary Tables 6 – 22: Supplementary Proteomic tables** | **18** |
| **Supplementary Figure 1: Generation of *B. cenocepacia* Δ*ogcI* Tn7-*ogcI.* A) Integration of miniTn7-rha-*ogc*I into *B. cenocepacia* Δ*ogcI*.** | **22** |
| **Supplementary Figure 2: Proteomic/glycoproteomic validation of *B. cenocepacia* Tn7-*ogcI*.** | **23** |
| **Supplementary Figure 3: Deletion of *ogcX*, *ogcA*, *ogcAB*, and *ogcB* from the Tmp^S^ parental strain.** | **24** |
| **Supplementary Figure 4. Plate-based growth assays of Δ*ogcI*Δ*ogcX* Tn7-*ogcI*, Δ*ogcI*Δ*ogcA* Tn7-*ogcI*, and Δ*ogcI* Tn7-*ogcI*.** | **25** |
| **Supplementary Figure 5. Complementation of *B. cenocepacia* Δ*ogcI* Δ*ogcA* Tn7-*ogcI.*** | **26** |
| **Supplementary Figure 6. Viability of *B. cenocepacia* Δ*ogcI*Δ*ogcA* Tn7-*ogcI* and Δ*ogcI*Δ*ogcX* Tn7-ogcI containing control plasmids in response to induction.** | **27** |
| **Supplementary Figure 7. Proteomic analysis of the protein of the *ogc* within Δ*ogcI*Δ*ogcX* Tn7-*ogcI*, Δ*ogcI*Δ*ogcA* Tn7-*ogcI*, Δ*ogcI*Δ*ogcB* Tn7-*ogcI* and Δ*ogcI* Tn7-*ogcI* strains.** | **28** |
| **Supplementary Figure 8. Proteomic analysis of *B. cenocepacia* Δ*ogcI* Tn7-*ogcI* strains.** | **29** |
| **Supplementary Figure 9. Comparison of proteomic alterations observed within *B. cenocepacia* Δ*ogc*IΔ*ogc*X Tn7-*ogc*I and Δ*ogc*IΔ*ogc*A Tn7-ogcI compared to *B. cenocepacia* glycosylation-null strains.** | **30** |
| **Supplementary Figure 10. Plate-based growth assays of Δ*ogcI* Tn7-*ogcI,* Δ*ogcI*Δ*ogcX* Tn7-*ogcI*, Δ*ogcI*Δ*ogcA* Tn7-*ogcI*, Δ*ogcI*Δ*ogcAB* Tn7-*ogcI* and Δ*ogcI*Δ*ogcB* Tn7-*ogcI* strains in the presence of membrane / osmotic stress agents.** | **31** |
| **Supplementary Figure 11. Viable count for Hoechst and NPN uptake assay.** | **32** |
| **Supplementary Figure 12. Hoechst and NPN uptake assays of complemented Δ*ogc*IΔ*ogc*A Tn7-*ogc*I and Δ*ogc*IΔ*ogc*X Tn7-*ogc*I.** | **33** |
| **Supplementary Figure 13. Antibiotic susceptibility assay of glycosylation inducible *B. cenocepacia* strains, with and without 1% Rhamnose induction.** | **34** |
| **Supplementary Figure 14. *B. cenocepacia* Δ*ogcI*Δ*ogcB* Tn7-*ogcI* Glycopeptides.** | **35** |
| **Supplementary Figure 15. Proteomics analysis of pSCrhaB2-*ogcAB in* Δ*ogcAB*.** | **36** |
| **Supplementary Figure 16. Proteomics analysis of pSCrhaB2-*ogcB* in Δ*ogcB*.** | **37** |
| **Supplementary Figure 17. Proteomics analysis of pSCrhaB2-*ogcI* in Δ*ogcI*.** | **38** |
| **Supplementary Figure 18. Proteomic analysis of Δ*ogcI*Δ*ogcA* Tn7-*ogcI* and Δ*ogcI*Δ*ogcX* Tn7-*ogcI* in response to glycosylation initiation.** | **39** |
| **Supplementary Figure 19. Spot plate assays of *E. coli pir 2* containing the expression vectors pSCrhaB2, pSCrhaB2-*ogcI (*p*ogcI)* and pSCrhaB2-*ogcB* (p*ogcB*).** | **40** |
| **References.** | **41** |

**Supplementary Document Table 1: Strain list**

| **Strain** | **Description** | **Source** |
| --- | --- | --- |
| ***E. coli*** | | |
| DH5α | F^−^ Φ80*lac*ZΔM15 Δ(*lac*ZYA-*argF*) U169 *rec*A1 *end*A1 *hsd*R17(r_K_^–^, m_K_^+^) *pho*A *sup*E44 *thi*-1 *gyr*A96 *rel*A1 λ^–^ | Invitrogen |
| PIR2 | F^−^ ∆*lac*169 *rpo*S(am) *rob*A1 *cre*C510 *hsd*R514 *end*A *rec*A1 *uidA*(∆*Mlu*I)::*pir* | Thermo Scientific |
| RHO3 | Δ*asd* *thi*-1 *thr*-1 *leu*B26 *ton*A21 lacY1 *sup*E44 *rec*A; integrated RP4-2 Tcr::Mu Δ*aphA* (λpir+) | ^1^ |
| ***B. cenocepacia*** | | |
| K56-2 WT | CF clinical isolate of the ET12 lineage, closely related to *B. cenocepacia* J2315 | Canadian *B. cepacia* research and referral repository ^2^ |
| K56-2 Δ*ogc* | Unmarked *ogc* (BCAL3114–BCAL3118) deletion | ^3^ |
| K56-2 Δ*ogcAB* | Unmarked *ogcAB* (BCAL3114–BCAL3118) deletion | ^3^ |
| K56-2 Δ*ogcB* | unmarked *ogcB* (BCAL3116) deletion generated using pGPI-SceI-*ogcB*/pDAI-SceI-SacB | This study |
| K56-2 Δ*ogcI* | Unmarked *ogcI* (BCAL3118) deletion generated using pGPI-SceI-*ogcI*/pDAI-SceI-SacB | This study |
| K56-2 Δ*ogcI* Tn7-*ogcI* | Trimethoprim sensitive (Tmp^S^) tetracycline sensitive (Tet^S^), Δ*ogcI* strain with chromosomally integrated *ogcI* under the rhamnose inducible promoter p_Rha_ with *rhaS/R* downstream of *glmS1* (BCAL0611) created using the miniTn7 delivery system. Tmp resistance cassette from miniTn7 element was subsequently removed using pFlpTet, and the strain was cured of pFlpTet. | This study |
| K56-2 Δ*ogcI* Tn7-*ogcI* Δ*ogcX* | K56-2 Δ*ogcI* Tn7-*ogcI* with unmarked *ogcX* (BCAL3114) deletion generated using pGPI-SceI-*ogcX*/pDAI-SceI-SacB | This study |
| K56-2 Δ*ogcI* Tn7-*ogcI* Δ*ogcA* | K56-2 Δ*ogcI* Tn7-*ogcI* with unmarked *ogcA* (BCAL3115) deletion generated using pGPI-SceI-*ogcA*/pDAI-SceI-SacB | This study |
| K56-2 Δ*ogcI* Tn7-*ogcI* Δ*ogcB* | K56-2 Δ*ogcI* Tn7-*ogcI* with unmarked *ogcB* (BCAL3116) deletion generated using pGPI-SceI-*ogcB*/pDAI-SceI-SacB | This study |
| K56-2 Δ*ogcI* Tn7-*ogcI* Δ*ogcAB* | K56-2 Δ*ogcI* Tn7-*ogcI* with unmarked *ogcAB* (BCAL3115-6) deletion generated using pGPI-SceI-*ogcAB*/pDAI-SceI-SacB | This study |

**Supplementary Document Table 2: Plasmid list**

| **Plasmid** | **Description** | **Source** |
| --- | --- | --- |
| pRK2013 | Helper plasmid for conjugation*, ori*_colE1_, RK2 derivative, Kan^R^, *mob*^+^, *tra*^+^ | ^4^ |
| pFlpTet | pFlpe4-derived plasmid, rhamnose-inducible *flp* (encoding flippase), Tet^R^, temperature-sensitive | ^5^ |
| pDAI-SceI-SacB | *ori*_pBBR_, Tet^R^, P_dhfr_, *mob^+^*, expressing I-SceI and SacB | ^6^ |
| pTNS3 | *ori*_R6K_, Amp^R^, expressing *tnsABCD* from *P1* and *P_lac_* (for Tn7 site-specific transposition). Addgene Plasmid #63127 | ^7^ |
| pUC18T-mini-Tn7T-Tp | mini-Tn7 suicide vector, *ori*_R6K_, *oriT* (for mobilisation), Amp^R^/Tmp^R^. Addgene Plasmid #65024 | ^7^ |
| pUC18T-mini-Tn7T-Tp-rha-ogcI | mini-Tn7 element containing rhamnose-inducible ogcI (BCAL3118) generated by subcloning the rhamnose-inducible *ogcI* element from pSCrhaB2-ogcI using *Nsi*I/*Hind*III into pUC18T-mini-Tn7T linearised with *Nsi*I/*Hind*III, Amp^R^ / Tmp^R^, | This study |
| pSCrhaB2 | *ori*_pBBR1_, *rhaR*, *rhaS*, P*_rhaB_*, *mob^+^,* Tmp^R^, Addgene Plasmid #113634 | ^8^ |
| pSCrhaB2-ogcI | *ogcI* (BCAL3118) cloned into pSCrhaB2 under the rhamnose-inducible promoter using Gibson assembly using the PCR product of primers Nsco_0526 / Nsco_0527 into *Nde*I/*Xba*I-linearised pSCrhaB2, Tmp^R^ | This study |
| pSCrhaB2-ogcA-met1 | *ogcA* (BCAL3115) from the assigned GTG start codon (modified to ATG) cloned into pSCrhaB2 under the rhamnose-inducible promoter using Gibson assembly using the PCR product of primers Nsco_0566 / Nsco_0525 into *Nde*I/*Xba*I-linearised pSCrhaB2, Tmp^R^ | This study |
| pSCrhaB2-ogcA-met2 | *ogcA* (BCAL3115) from the putative alternative start codon of *ogcA* located within *ogcX* cloned into pSCrhaB2 under the rhamnose-inducible promoter using Gibson assembly using the PCR product of primers Nsco_0567 / Nsco_0525 into *Nde*I/*Xba*I-linearised pSCrhaB2, Tmp^R^ | This study |
| pSCrhaB2-ogcAB-met1 | *ogcAB* (BCAL3115-6) from the assigned GTG start codon (modified into ATG) of *ogc*A cloned into pSCrhaB2 under the rhamnose-inducible promoter by ligating the *Nde*I/*Xba*I-digested PCR product of primers Nsco_0215 / Nsco_0567 and *Nde*I/*Xba*I-linearised pSCrhaB2, Tmp^R^ | This study |
| pSCrhaB2-ogcAB-met2 | *ogcAB* (BCAL3115-6) from the putative alternative start codon of *ogcA* located within *ogcX* cloned into pSCrhaB2 under the rhamnose-inducible promoter by ligating the *Nde*I/*Xba*I-digested PCR product of primers Nsco_0215 / Nsco_0911 and *Nde*I/*Xba*I-linearised pSCrhaB2, Tmp^R^ | This study |
| pSCrhaB2-ogcB | *ogcB* (BCAL3116) cloned into pSCrhaB2 under the rhamnose-inducible promoter using Gibson assembly using the PCR product of primers Nsco_0214 / Nsco_0215 into *Nde*I/*Xba*I-linearised pSCrhaB2, Tmp^R^ | This study |
| pSCrhaB2-ogc | *ogc* (BCAL3114-BCAL3118) cloned into pSCrhaB2 under the rhamnose-inducible promoter using Gibson assembly using the PCR product of primers Nsco_0919 / Nsco_0920 and *Nde*I/*Xba*I-linearised pSCrhaB2, Tmp^R^ | This study |
| pSCrhaB2-ogcΔogcX | pSCrhaB2 containing the *ogc* cluster under the rhamnose-inducible promoter with *ogcX* (BCAL3114) removed. Generated using Gibson assembly of the PCR products of primers Nsco_0216 / Nsco_1187 and Nsco_0217 / Nsco_1186 and *Nde*I/*Hind*III-linearised pSCrhaB2, Tmp^R^ | This study |
| pSCrhaB2-ogcΔogcA | pSCrhaB2 containing the *ogc* cluster under the rhamnose-inducible promoter with *ogcA* (BCAL3115) removed. Generated using Gibson assembly of the PCR products of primers Nsco_0216 / Nsco_1189 and Nsco_0217 / Nsco_1188 and *Nde*I/*Hind*III-linearised pSCrhaB2, Tmp^R^ | This study |
| pSCrhaB2-ogcΔogcB | pSCrhaB2 containing the *ogc* cluster under the rhamnose-inducible promoter with *ogcB* (BCAL3116) removed. Generated using Gibson assembly of the PCR products of primers Nsco_0216 / Nsco_1191 and Nsco_0217 / Nsco_1190 and *Nde*I/*Hind*III-linearised pSCrhaB2, Tmp^R^ | This study |
| pSCrhaB2-ogcΔogcI | pSCrhaB2 containing the *ogc* cluster under the rhamnose-inducible promoter with *ogcI* (BCAL3118) removed. Generated using Gibson assembly of the PCR products of primers Nsco_0216 / Nsco_1193 and Nsco_0217 / Nsco_1192 and *Nde*I/*Hind*III-linearised pSCrhaB2, Tmp^R^ | This study |
| pGPI-SceI | *ori_R6K_*, *mob*^+^, I-SceI restriction site, Tmp^R^ | ^9^ |
| pGPI-SceI-ogcX | pGPI-SceI containing fragments flanking *ogcX* (BCAL3114) generated using Gibson assembly of the PCR products of primers Nsco_0279 / Nsco_0280 and Nsco_0281 / Nsco_0282 with *Sma*I-linearised pGPI-SceI, Tmp^R^ | This study |
| pGPI-SceI-ogcA | pGPI-SceI containing fragments flanking *ogcA* (BCAL3115) generated using Gibson assembly of the PCR products of primers Nsco_0176 / Nsco_0177 and Nsco_0178 / Nsco_0179 with *Sma*I-linearised pGPI-SceI, Tmp^R^ | This study |
| pGPI-SceI-ogcB | pGPI-SceI containing fragments flanking *ogcB* (BCAL3116) generated using Gibson assembly of the PCR products of primers Nsco_0182 / Nsco_0183 and Nsco_0184 / Nsco_0185 with *Sma*I-linearised pGPI-SceI, Tmp^R^ | This study |
| pGPI-SceI-ogcAB | pGPI-SceI containing fragments flanking *ogcAB* (BCAL3115-6) generated using Gibson assembly of the PCR products of primers Nsco_0213 / Nsco_0185 and Nsco_0194 / Nsco_0212 with *Sma*I-linearised pGPI-SceI, Tmp^R^ | This study |
| pGPI-SceI-ogcI | pGPI-SceI containing fragments flanking *ogcI* (BCAL3118) generated using Gibson assembly of the PCR products of primers Nsco_0271 / Nsco_0272 and Nsco_0273 / Nsco_0274 with *Sma*I-linearised pGPI-SceI, Tmp^R^ | This study |
| pUS250-sfGFP | Derivative of pUS250 (Addgene plasmid # 198322) containing the cumate-inducible system of *Pseudomonas putida* (CymR) regulating the expression of superfolder green fluorescent protein (*sfgfp*) gene, Kan^R^ | Nick Coleman (unpublished) |
| pMLBAD | *ori*_pBBR1_, *araC* P_BAD_, *mob^+^*, Tmp^R^ | ^10^ |
| pCumate-sfGFP | pMLBAD backbone containing *ori*_pBBR1_, *mob^+^*, modified to contain the cumate-inducible system from *P. putida* (CymR) regulating the expression of superfolder green fluorescent protein (*sfgfp*) gene generated using Gibson assembly of the PCR product of primers Nsco_1094 / Nsco_1095 from pUS250-sfGFP and the amplified pMLBAD generated using primers Nsco_1092/Nsco_1093, Tmp^R^ | This study |
| pCumate-ogcX-his | pMLBAD with cumate-inducible system from *P. putida* (CymR) regulating the expression of *ogcX*-His_6_ generated using Gibson assembly of the PCR product of primers Nsco_1453 / Nsco_1458 and the amplified pMLBAD-cumate backbone from pCumate-sfGFP generated using primers Nsco_1455 / Nsco_1457, Tmp^R^ | This study |
| pCumate-BCAL2087 | pMLBAD with cumate-inducible system from *P. putida* (CymR) regulating the expression of *uppS* (BCAL2087) generated using Gibson assembly of the PCR product of primers Nsco_1588 / Nsco_1589 and the amplified pMLBAD-cumate backbone from pCumate-sfGFP generated using primers Nsco_1590 / Nsco_1591, Tmp^R^ | This study |
| pCumate-BCAL2067 | pMLBAD with cumate-inducible system from *Pseudomonas putida* regulating the expression of *uppS* (BCAM2067) generated using Gibson assembly of the PCR product of primers Nsco_1594 / Nsco_1662 and the amplified pMLBAD-cumate backbone from pCumate-sfGFP generated using primers Nsco_1419 / Nsco_1420, Tmp^R^ | This study |

Kan^R^, kanamycin resistance, Tet^R^, tetracycline resistance, Amp^R^, ampicillin resistance, Tmp^R^, trimethoprim resistance

**Supplementary Document Table 3: Primer list**

| **Primer** | **Sequence** | **Description / Purpose** |
| --- | --- | --- |
| mini-Tn7 | | |
| Nsco_0190 | cgaaccgaacaggcttatgt | Forward screening primers to assess insertion into pUC18T-mini-Tn7T-Tp or mini-Tn7 integration into chromosome |
| Nsco_0191 | ctgtgggcggacaaaatagt | Reverse screening primers to assess of insertion into pUC18T-mini-Tn7T-Tp or mini-Tn7 integration into chromosome |
| Nsco_0193 | cacagcataactggactgatttc | Internal Tn7 right end primer for screening mini-Tn7 integration ^7^ |
| Nsco_0784 | cataagcctgttcggttcgt | Binds between Tn7L and Tn7R within mini-Tn7 element. Paired with Nsco_0158 or Nsco_0201 verifies the mini-Tn7 integration into chromosome |
| Nsco_0157 | cggtcgagttagcacaggat | *B. cenocepacia* *glmS1* (*BCAL0611*) Att site forward screening primers |
| Nsco_0158 | caactgctcgcgtatcacac | *B. cenocepacia* *glmS1* (*BCAL0611*) Att site reverse screening primers |
| Nsco_0200 | gtggagcaccatttcgtgag | *B. cenocepacia* *glmS2 (BCAM0478)* Att site forward screening primers |
| Nsco_0201 | tgataagccgaggaatttgg | *B. cenocepacia* *glmS2 (BCAM0478)* Att site reverse screening primers |
| 6135 | ggattcgacatgggtcaaag | Forward Screening primer for the Tmp^R^ cassette within mini-Tn7 |
| Nsco_0931 | gctcgaattagcttcaaaagcgc | Reverse Screening primer for the Tmp^R^ cassette within mini-Tn7 |
| Mutation construct (pGPI-SceI) construction and mutant screening | | |
| Nsco_0279 | gcatgcgatatcgagctctccctcttgaagccgttgtagtcg | Upstream *ogcX* forward primer for creation of pGPI-SceI-ogcX |
| Nsco_0280 | gtcatgttcgcagcttgaaattagatctgggtgagccgttc | Upstream *ogcX* reverse primer for creation of pGPI-SceI-ogcX |
| Nsco_0281 | gaacggctcacccagatctaatttcaagctgcgaacatgac | Downstream *ogcX* forward primer for creation of pGPI-SceI-ogcX |
| Nsco_0282 | cggataacaatttgtggaattcccgaaatgcagcaggttcttgg | Downstream *ogcX* reverse primer for creation of pGPI-SceI-ogcX |
| Nsco_0194 | gcatgcgatatcgagctctcccaatcctcgtcccgaccat | Upstream *ogcAB* forward primer for creation of pGPI-SceI-ogcAB |
| Nsco_0212 | cgatgcttccatcagcgtctagccggcatcagcacggccac | Upstream *ogcAB* reverse primer for creation of pGPI-SceI-ogcAB |
| Nsco_0213 | gtggccgtgctgatgccggctagacgctgatggaagcatcg | Downstream *ogcAB* forward primer for creation of pGPI-SceI-ogcAB |
| Nsco_0182 | gcatgcgatatcgagctctcccgtgcacgtgctgatcgtc | Upstream *ogcB* forward primer for creation of pGPI-SceI-ogcB |
| Nsco_0183 | cgatgcttccatcagcgtctaaggtggcggacatgaaaa | Upstream *ogcB* reverse primer for creation of pGPI-SceI-ogcB |
| Nsco_0184 | ttttcatgtccgccaccttagacgctgatggaagcatcg | Downstream *ogcB* forward primer for creation of pGPI-SceI-ogcB |
| Nsco_0185 | cggataacaatttgtggaattcccgcgatcagcttcgactgg | Downstream *ogcB* reverse primer for creation of pGPI-SceI-ogcB/pGPI-SceI-ogcAB |
| Nsco_0176 | gcatgcgatatcgagctctcccgttcgacaagccgctgtt | Upstream *ogcA* forward primer for creation of pGPI-SceI-ogcA |
| Nsco_0177 | gtcacgaaatgcagcaggtctagtcgtcgtgcccgttgtag | Upstream *ogcA* reverse primer for creation of pGPI-SceI-ogcA |
| Nsco_0178 | ctacaacgggcacgacgactagacctgctgcatttcgtgac | Downstream *ogcA* forward primer for creation of pGPI-SceI-ogcA |
| Nsco_0179 | cggataacaatttgtggaattcccgttcgagatcgacgccttc | Downstream *ogcA* reverse primer for creation of pGPI-SceI-ogcA |
| Nsco_0271 | gcatgcgatatcgagctctcccccgtgctcaaggtcatgc | Upstream *ogcI* forward primer for creation of pGPI-SceI-ogcI |
| Nsco_0272 | gtatgccgccagaacagcctatagcagggagacgatgaagc | Upstream *ogcI* reverse primer for creation of pGPI-SceI-ogcI |
| Nsco_0273 | gcttcatcgtctccctgctataggctgttctggcggcatac | Downstream *ogcI* forward primer for creation of pGPI-SceI-ogcI |
| Nsco_0274 | cggataacaatttgtggaattccccctgtacggtttctcggaag | Downstream *ogcI* reverse primer for creation of pGPI-SceI-ogcI |
| 6108 | taacggttgtggacaacaagccaggg | Forward primer for screening insertion into pGPI-SceI and integration of pGPI-SceI into the chromosomes ^11^ |
| 6109 | gccctacacaaattgggagatatatc | Reverse primer for screening insertion into pGPI-SceI and integration of pGPI-SceI into the chromosomes ^11^ |
| Nsco_0275 | tgactcgcatgatcgaactc | Forward primer to verify *ogcI* deletion |
| Nsco_0276 | cgtcacttcgtgctgatctc | Reverse primer to verify *ogcI* deletion |
| Nsco_0180 | atgcgaatacgcttctcctg | Forward primer to verify *ogcA* deletion |
| Nsco_0181 | agggtgataccggttaacga | Reverse primer to verify *ogcA* deletion |
| Nsco_0186 | tttcaagctgcgaacatgac | Forward primer to verify *ogcB* deletion |
| Nsco_0187 | atacggcatcaggttgttcg | Reverse primer to verify *ogcB* deletion |
| Nsco_0283 | gatctgctcgccgtagattg | Forward primer to verify *ogcX* deletion |
| Nsco_0284 | atcagcccgtggcgatac | Reverse primer to verify *ogcX* deletion |
| 6689 | accacgccacgaatgtcata | Forward primer to verify *ogc* deletion |
| 6690 | cgaacatcatgaagctgacc | Reverse primer to verify *ogc* deletion |
| Nsco_0780 | ggcaactatcgggcaaagtaccgc | Forward primer to verify *ogcA* deletion, binds internally within *ogcA* |
| Nsco_0781 | gcggtactttgcccgatagttgcc | Reverse primer to verify *ogcA* deletion, binds internally within *ogcA* |
| 7162 | gatctgctcgccgtagattg | Forward primer to verify the removal of the *ogc* |
| 7163 | gcgagaagctttacgaggaa | Reverse primer to verify the removal of the *ogc* |
| Construction of pSCrhaB2/pCumate expression vectors | | |
| Nsco_0216 | cgaattcaggcgctttttag | pSCrhaB2-ogc screening forward primer |
| Nsco_0217 | acggcgtttcacttctgagt | pSCrhaB2-ogc screening reverse primer |
| Nsco_0698 | ctttccctggttgccaatggccc | pSCrhaB2-ogc screening forward primer alternative 1 |
| Nsco_0699 | cggcgtttcacttctgagttcggc | pSCrhaB2-ogc screening reverse primer alternative 2 |
| Nsco_0777 | tgagcatcacatcaccacaattcagc | pSCrhaB2-ogc screening forward primer alternative 2 |
| Nsco_0778 | ccgccaggcaaattctgttttatcagac | pSCrhaB2-ogc screening reverse primer alternative 3 |
| Nsco_0526 | cgtaatgaaattcagcaggatcacatatgctcagcttcgcgtccgg | Forward primer containing *Nde*I for *ogcI* to insert into pSCrhaB2 |
| Nsco_0527 | tgcctgcaggtcgactctagagtatcgctcaacggctctgt | Reverse primer containing *Xba*I for *ogcI* to insert into pSCrhaB2 |
| Nsco_0566 | cgtaatgaaattcagcaggatcacatAtggccgtgctgatgccggcctacaacg | Forward primer containing *Nde*I to insert *ogcA* met1 for insertion into pSCrhaB2 |
| Nsco_0567 | cgtaatgaaattcagcaggatcacatatgacgtcccctgcttgcccgacc | Forward primer containing *Nde*I to insert *ogcA*-met2 for insertion into pSCrhaB2 |
| Nsco_0525 | tgcctgcaggtcgactctagagcgtcgaaattgccgaac | Reverse primer containing *Xba*I for amplification of *ogcA* and insertion into pSCrhaB2 |
| Nsco_0911 | cgtaatgaaattcagcaggatcacatatggctgctcgcgctggccatcg | Forward primer containing *Nde*I site upstream of the alterative start codon *of ogcA* for amplification of *ogcAB* and insertion into pSCrhaB2 |
| Nsco_0214 | aaaaacatatgtccgccacctccccgctgcgc | Forward primer containing *Nde*I site for amplification of *ogcB* and insertion into pSCrhaB2 |
| Nsco_0215 | aaaaatctagagctgatgcgctccttcagagg | Reverse primer containing *Xba*I for amplification of *ogcB* and insertion into pSCrhaB2 |
| Nsco_0919 | gaaattcagcaggatcacatatgctgaagcgcttcggcaacccgg | Forward primer for amplification of the *ogc* cluster for insertion into pSCrhaB2 |
| Nsco_0920 | gcatgcctgcaggtcgactctaggggtgacgtggctccaggccggaac | Reverse primer for amplification of the *ogc* cluster for insertion into pSCrhaB2 |
| Nsco_1092 | ctgaaatttgcttcggggtcattatagggtctgataaaacagaatttgcctgg | Forward primer to amplify the pMLBAD vector and allow integration of CymR- sfGFP |
| Nsco_1093 | cctttttctttaaaaccgaaaagattaccgatgggagatcctaagatatcgc | Reverse primer to amplify the pMLBAD vector and allow integration of CymR- sfGFP |
| Nsco_1094 | ccaggcaaattctgttttatcagaccctataatgaccccgaagcaaatttcag | Forward primer to amplify CymR-sfGFP for insertion into pMLBAD |
| Nsco_1095 | gcgatatcttaggatctcccatcggtaatcttttcggttttaaagaaaaagg | Reverse primer to amplify CymR-sfGFP for insertion into pMLBAD |
| Nsco_1455 | ccgggttgccgaagcgcttcagcattatcttacctccttaatttgatttc | Forward primer to amplify pCumate vector backbone and the insertion *ogcX* |
| Nsco_1457 | ttacatccgtttcaagctgcgaacacaccaccaccaccaccactgatgatactagtagcggccgctgcagc | Forward primer to amplify pCumate vector backbone and the insertion *ogcX*-His_6_ |
| Nsco_1453 | gaaatcaaattaaggaggtaagataatgctgaagcgcttcggcaacccgg | Forward primer to amplify ogcX for insertion into pCumate |
| Nsco_1458 | gctgcagcggccgctactagtatcatcagtggtggtggtggtggtgtgttcgcagcttgaaacggatgtaa | Reverse primer to amplify *ogc*X with a His_6_-tag for insertion into pCumate |
| Nsco_1590 | cggtagagctggtataggtcattatcttacctccttaatttgatttc | Forward primer to amplify pCumate vector backbone and the insertion of BCAL2087 (UppS1) |
| Nsco_1591 | gcagaacgccgactccctttcatgctgatgatactagtagcggccgctgcagc | Reverse primer to amplify pCumate vector backbone and the insertion of BCAL2087 (UppS1) |
| Nsco_1588 | gaaatcaaattaaggaggtaagataatgacctataccagctctaccg | Forward primer to amplify BCAL2087 (UppS1) for insertion into pCumate |
| Nsco_1589 | gctgcagcggccgctactagtatcatcagcatgaaagggagtcggcgttctgc | Reverse primer to amplify BCAL2087 (UppS1) for insertion into pCumate |
| Nsco_1419 | cattatcttacctccttaatttgatttc | Forward primer to amplify pCumate vector backbone and the insertion of BCAM2067 (UppS2) |
| Nsco_1420 | tgatactagtagcggccgctgcagca | Reverse primer to amplify pCumate vector backbone and the insertion of BCAM2067 (UppS2) |
| Nsco_1594 | gaaatcaaattaaggaggtaagataatgactcaagagctgattctgcgcg | Forward primer to amplify BCAM2067 (UppS2) for insertion into pCumate |
| Nsco_1662 | gctgcagcggccgctactagtatcaaacgggtgtcatgaaggacggtcc | Reverse primer to amplify BCAM2067 (UppS2) for insertion into pCumate |
| Nsco_1502 | cccagaatgttaccatcctctt | Forward screening primer to verify insertion of sfGFP into pCumate |
| Nsco_1256 | agatctgccatgagacccaa | Reverse screening primer to verify insertion of sfGFP into pCumate |
| Nsco_1473 | ttcggtgatctgttcgtaaagc | Forward sequencing primer to verify insertion into pCumate |
| Nsco_1308 | gccttgaccgaaacggaggaat | Reverse sequencing primer to verify insertion into pCumate |
| pSCrhaB2 mutagenesis | | |
| Nsco_1186 | cgtaatgaaattcagcaggatcacattgacgtcccctgcttgcccgaccccgc | Forward primer for deletion of *ogcX* from pSCrhaB2-*ogc* |
| Nsco_1187 | gcggggtcgggcaagcaggggacgtcaatgtgatcctgctgaatttcattacg | Reverse primer for deletion of *ogcX* from pSCrhaB2-*ogc* |
| Nsco_1188 | cgctcgacgacgtggccgtgctgtgacgtgcatccggcccgttcggcaatttcg | Forward primer for deletion of *ogcA* from pSCrhaB2-*ogc* |
| Nsco_1189 | cgaaattgccgaacgggccggatgcacgtcacagcacggccacgtcgtcgagcg | Reverse primer for deletion of *ogcA* from pSCrhaB2-*ogc* |
| Nsco_1190 | cgacgctttattttttctttttttcggaattcgacgaacagcaggtcgtcgaacg | Forward primer for deletion of *ogcB* from pSCrhaB2-*ogc* |
| Nsco_1191 | cgttcgacgacctgctgttcgtcgaattccgaaaaaaagaaaaaataaagcgtcg | Reverse primer for deletion of *ogc*B from pSCrhaB2-*ogc* |
| Nsco_1192 | ccgctgtccattttccgagcggtccctgagcgatacgctcgcttccgaacaaaaaag | Forward primer for deletion of *ogcI* from pSCrhaB2-*ogc* |
| Nsco_1193 | cttttttgttcggaagcgagcgtatcgctcagggaccgctcggaaaatggacagcgg | Reverse primer for deletion of *ogc*I from pSCrhaB2-*ogc* |

**Supplementary Table 4: Proteomic Dataset**

| **Pride accession number (Review login details)** | **MS instrument** | **Number of Biological groups, replicates and total datafiles** | **Description of dataset** |
| --- | --- | --- | --- |
| PXD054841  Username: Password: vtOYlh2QeMmH | Orbitrap Fusion Lumos | 2 biological groups, 4 replicates total of 8 datafiles | DDA experiment assessing glycosylation and expression of OgcB from pSCrhaB2-OgcB or pSCrhaB2 within K56-2 Δ*ogc*B at stationary phase within LB with 0.05% rhamnose. |
| PXD054867  Username: Password: 1KCRw6fckXC0 | Orbitrap Fusion Lumos | 2 biological groups, 4 replicates total of 8 datafiles | DDA experiment assessing glycosylation and expression of OgcI from pSCrhaB2-OgcI or pSCrhaB2 within K56-2 Δ*ogcI* at stationary phase within LB with 0.05% rhamnose. |
| PXD055163  Username: Password: xQ0vECsjoX74 | Orbitrap Fusion Lumos | 6 biological groups, 4 replicates total of 24 datafiles | DDA experiment assessing glycosylation and expression of OgcAB from pSCrhaB2-OgcAB or pSCrhaB2 within K56-2 Δ*ogcAB* at stationary phase within LB with 1% rhamnose. |
| PXD054929  Username: Password: QFVtjaWskLkP | Orbitrap Fusion Lumos equipped with a FAIMS Pro interface | 8 biological groups, 4 replicates total of 32 datafiles | DDA experiment assessing glycosylation within K56-2 Δ*ogc* and K56-2 WT containing pSCrhaB2-*ogc* or pSCrhaB2 at stationary phase within LB with 1% rhamnose. |
| PXD054956  Username: Password: irFS6N8GHplR | Orbitrap Fusion Lumos | 7 biological groups, 4 replicates total of 28 datafiles | DDA experiment assessing glycosylation and expression of *ogcA* from pSCrhaB2-*ogcA*; pSCrhaB2-*ogcA*_alt_; within K56-2 Δ*ogcI*Δ*ogcA* Tn7-*ogc*I and K56-2 Δ*ogcI* Tn7-*ogc*I at stationary phase within LB with 1% rhamnose with pSCrhaB2 containing strains used as negative controls. |
| PXD056440  **Username:**  **Password:** 4e7AjHcW3n8v | Orbitrap Fusion Lumos | 10 biological groups, 4 replicates total of 40 datafiles | DDA experiment assessing glycosylation and expression of *ogcX* from pCumate-*ogcX-his* within K56-2 Δ*ogcI*Δ*ogcX* Tn7-*ogc*I and K56-2 Δ*ogcI* Tn7-*ogc*I at stationary phase within LB with 1% rhamnose/100 μM cumate with  pCumate-sfGFP containing strains used as negative controls. |
| PXD054923  Username: Password: ZiiuDcoa2nre | Orbitrap Fusion Lumos equipped with a FAIMS Pro interface | 6 biological groups, 4 replicates total of 24 datafiles | DDA experiment assessing glycosylation at stationary phase within LB of K56-2 Δ*ogcI* Tn7-*ogc*I (non-induced) compared to K56-2 Δ*ogcI* Tn7-*ogc*I; K56-2 Δ*ogc*IΔ*ogcA* Tn7-*ogc*I; K56-2 Δ*ogcI*Δ*ogcAB* Tn7-*ogc*I; K56-2 Δ*ogcI*Δ*ogcB* Tn7-*ogc*I and K56-2 Δ*ogcI*Δ*ogcX* Tn7-*ogcI* induced with 0.1% rhamnose. |
| PXD054949  Username: Password: imU103bFkKkO | Orbitrap Fusion Lumos | 8 biological groups, 4 replicates total of 32 datafiles | DIA experiment assessing proteome changes at stationary phase within LB of K56-2 Δ*ogcI* Tn7-*ogcI*; K56-2 Δ*ogcI*Δ*ogcA* Tn7-*ogcI*; K56-2 Δ*ogcI*Δ*ogcB* Tn7-*ogc*I and K56-2 Δ*ogcI*Δ*ogcX* Tn7-*ogc*I (non-induced) compared to K56-2 Δ*ogc*I Tn7-*ogcI*; K56-2 Δ*ogc*IΔ*ogcA* Tn7-*ogcI*; K56-2 Δ*ogc*IΔ*ogcB* Tn7-*ogc*I and K56-2 Δ*ogcI*Δ*ogcX* Tn7-*ogcI* induced with 0.1% rhamnose. |
| PXD055576  Username: Password: PLRffXCkJY41 | Orbitrap Fusion Lumos equipped with a FAIMS Pro interface | 3 biological groups, 4 replicates total of 12 datafiles | DDA experiment assessing glycosylation at stationary phase within LB of K56-2 Δ*ogcI* Tn7-*ogcI* (non-induced) compared to K56-2 Δ*ogcI* Tn7-*ogcI* with 0.05% and 0.1% rhamnose. |
| PXD059283  Username: Password: 8F2GdarAPyoS | Orbitrap Fusion Lumos | 8 biological groups, 4 replicates total of 32 datafiles | DDA experiment assessing the impact of OgcX overexpression from pCumate-*ogcX-his* on glycan patterns within  K56-2 Δ*ogcI* Tn7-*ogcI*; K56-2 Δ*ogcI*Δ*ogcA* Tn7-*ogcI*; K56-2 Δ*ogcI*Δ*ogcB* Tn7-*ogc*I and K56-2 Δ*ogcI*Δ*ogcX* Tn7-*ogc*I  at stationary phase in LB with and without 1% rhamnose/100 μM cumate. |

**Supplementary Table 5: Antimicrobial Minimal inhibitory concentrations (MIC).**

| **Strains** | **Rhamnose** | **Tetracycline** | **Trimethoprim** | **Rifampicin** | **Ceftazidime** | **Chlorhexidine** |
| --- | --- | --- | --- | --- | --- | --- |
| Δ*ogcI* Tn7-*ogcI* | 0% | 32-64 | 16 | 64 | 32 | 16-32 |
| Δ*ogcI* Tn7-*ogcI* | 1% | 32-64 | 32 | 64 | 32 | 16-32 |
| Δ*ogcI*Δ*ogcX* Tn7-*ogcI* | 0% | 32-64 | 16 | 64 | 32 | 16-32 |
| Δ*ogcI*Δ*ogcX* Tn7-*ogcI* | 1% | 32-64 | 16 | 64 | 4 | 16-32 |
| Δ*ogcI*Δ*ogcA* Tn7-*ogcI* | 0% | 32-64 | 16 | 16-64 | 64 | 16-32 |
| Δ*ogcI*Δ*ogcA* Tn7-*ogcI* | 1% | 32-64 | 16 | 64 | 4 | 16-32 |
| Δ*ogcI*Δ*ogcB* Tn7-*ogcI* | 0% | 32-64 | 16 | 64 | 32 | 16-32 |
| Δ*ogcI*Δ*ogcB* Tn7-*ogcI* | 1% | 32-64 | 32 | 64 | 64 | 16-32 |

The antimicrobial susceptibility of bacterial strains was determined via broth dilution methods in accordance with Clinical and Laboratory Standards Institute (CLSI) guidelines. Strains were cultured in LB broth to an OD_600_ of 0.5–0.6, followed by a 1:100 dilution in cation-adjusted Mueller–Hinton broth (CAMHB) with and without the addition of rhamnose. Serial twofold dilutions of antibiotics—trimethoprim, tetracycline, chlorhexidine, ceftazidime, and rifampicin—were prepared in CAMHB within flat-bottom microplates. Bacterial inoculum was added to each well, with one well per plate reserved as a growth control (no antibiotics) and another as a sterility control (no inoculum). Plates were incubated at 37°C for 20 hours. Post-incubation, microbial growth was assessed visually, and optical density was measured at 600 nm to quantify bacterial growth.

**Supplementary Proteomics tables**

**Supplementary Table 6. DDA peptide-spectrum matches (PSMs) summary of *B. cenocepacia* WT and Δ*ogc* containing pSCrhaB2 and pSCrhaB2-*ogc****.* The MSfragger PSM search summary for the proteome analysis of *B. cenocepacia* WT and Δ*ogc* strains containing pSCrhaB2 and pSCrhaB2-*ogc* are provided. For each modification type identified within the proteome the total number of PSMs assigned are tabulated.

**Supplementary Table 7. DDA protein level LFQ analysis of *B. cenocepacia* WT and Δ*ogc* containing pSCrhaB2 and pSCrhaB2-*ogc*.** The Perseus processed MSfragger search results for the protein analysis of four biological replicates of strains WT and Δ*ogc* containing the plasmids pSCrhaB2 and pSCrhaB2-*ogc* are provided. For each identified protein, the log2 LFQ protein values, the T-test information including the -log_10_(*p*-value), differences in the mean between the groups and if the resulting *p*-values is below the multiple hypothesis corrected *p*-value are provided. For each protein the top peptide probability, total number of MS/MS events for the corresponding protein as well as both the imputed and non-imputed data is provided.

**Supplementary Table 8. DDA PSM summary of *B. cenocepacia* Δ*ogcI* Tn7-*ogcI* strains with and without induction*.*** The MSfragger PSMs search summary for the proteome analysis of *B. cenocepacia* Δ*ogcI* Tn7-*ogcI* with and without induction for *B. cenocepacia* Δ*ogcI*Δ*ogcX* Tn7-*ogcI, B. cenocepacia* Δ*ogcI*Δ*ogcA* Tn7-*ogcI, B. cenocepacia* Δ*ogcI*Δ*ogcB* Tn7-*ogcI and B. cenocepacia* Δ*ogcI*Δ*ogcAB* Tn7-*ogcI* is provided. For each modification type identified within the proteome the total number of PSMs assigned are tabulated.

**Supplementary Table 9. DDA protein level LFQ analysis of *B. cenocepacia* Δ*ogcI* Tn7-*ogcI* and *B. cenocepacia* Δ*ogcI*Δ*ogcA* Tn7-*ogcI* containing pSCrhaB2, pSCrhaB2-*ogcA*-Met1, or pSCrhaB2-*ogcA*-Met2.** The Perseus processed MSfragger search results for the protein analysis of four biological replicates of strains *B. cenocepacia* Δ*ogcI* Tn7-*ogcI* and *B. cenocepacia* Δ*ogcI*Δ*ogcA* Tn7-*ogcI* containing the plasmids pSCrhaB2, pSCrhaB2-*ogcA*-Met1, or pSCrhaB2-*ogcA*-Met2 without and with 1% rhamnose (with the exception of *B. cenocepacia* Δ*ogcI*Δ*ogcA* Tn7-*ogcI* pSCrhaB2 due to lack of viability). For each identified protein, the log2 LFQ protein values, the T-test information including the -log_10_(*p*-value), differences in the mean between the groups and if the resulting *p*-values is below the multiple hypothesis corrected *p*-value are provided. For each protein the top peptide probability, total number of MS/MS events for the corresponding protein as well as both the imputed and non-imputed data is provided.

**Supplementary Table 10. DDA PSM summary of *B. cenocepacia* Δ*ogcI* Tn7-*ogcI* and *B. cenocepacia* Δ*ogcI*Δ*ogcA* Tn7-*ogcI* containing pSCrhaB2, pSCrhaB2-*ogcA*-Met1, or pSCrhaB2-*ogcA*-Met2 strains with and without induction*.*** The MSfragger PSM search summary for the proteome analysis of *B. cenocepacia* Δ*ogcI* Tn7-*ogcI* and *B. cenocepacia* Δ*ogcI*Δ*ogcA* Tn7-*ogcI* containing pSCrhaB2, pSCrhaB2-*ogcA*-Met1, or pSCrhaB2-*ogcA*-Met2 strains is provided. For each modification type identified within the proteome the total number of PSMs assigned are tabulated.

**Supplementary Table 11. DDA protein level LFQ analysis of *B. cenocepacia* Δ*ogcI* Tn7-*ogcI* and *B. cenocepacia* Δ*ogcI*Δ*ogcX* Tn7-*ogcI* containing pCumate-sfGFP and pCumate-*ogcX-his*.** The Perseus processed MSfragger search results for the protein analysis of four biological replicates of strains *B. cenocepacia* Δ*ogcI* Tn7-*ogcI* and *B. cenocepacia* Δ*ogcI*Δ*ogcX* Tn7-*ogcI* containing the plasmids pCumate-sfGFP and pCumate-*ogcX-his* without and with 1% rhamnose/100 μM cumate. For each identified protein, the log2 LFQ protein values, the T-test information including the -log_10_(*p*-value), differences in the mean between the groups and if the resulting *p*-values is below the multiple hypothesis corrected *p*-value are provided. For each protein the top peptide probability, total number of MS/MS events for the corresponding protein as well as both the imputed and non-imputed data is provided.

**Supplementary Table 12. DDA PSM summary of *B. cenocepacia* Δ*ogcI* Tn7-*ogcI* and *B. cenocepacia* Δ*ogcI*Δ*ogcX* Tn7-*ogcI* containing pCumate-sfGFP and pCumate-*ogcX-his* with and without induction*.*** The MSfragger PSMs search summary for the proteome analysis of *B. cenocepacia* Δ*ogcI* Tn7-*ogcI* and *B. cenocepacia* Δ*ogcI*Δ*ogcX* Tn7-*ogcI* containing pCumate-sfGFP and pCumate-*ogcX-his* strains are provided. For each modification type identified within the proteome the total number of PSMs assigned are tabulated.

**Supplementary Table 13. DIA Proteomic analysis of Δ*ogcI* Tn7-ogcI, Δ*ogcI*Δ*ogcA* Tn7-*ogcI*, Δ*ogcI*Δ*ogcB* Tn7-*ogcI*, and Δ*ogcI*Δ*ogcX* Tn7-*ogcI* strains with and without rhamnose induction (1% rhamnose).** The Spectronaut search results for protein level analysis of biological replicates (n=4) grown +/- the addition of Rhamnose of Δ*ogcI* Tn7-ogcI, Δ*ogcI*Δ*ogcA* Tn7-*ogcI*, Δ*ogcI*Δ*ogcB* Tn7-*ogcI*, and Δ*ogcI*Δ*ogcX* Tn7-*ogcI*. Imputed and non-imputed Perseus processed data with statistical analysis is provided. For each identified protein, the log_2_ LFQ protein values, T-‑test information including the -log_10_(*p*-value), difference in the mean between the groups and if the resulting *p*-values are below 0.05 and the multiple hypothesis corrected *p*‑values (permutation-based false discovery rate of 0.05) are provided. Categorical information associated with protein accessions, gene name, and GO terms are provided in addition to if proteins were identified by a single PSM within a given experiment as well as the total number of precursors assigned for each protein within each replicate.

**Supplementary Table 14. Enrichment analysis of proteome changes observed within Δ*ogcI*Δ*ogcX* Tn7-*ogcI* and Δ*ogcI*Δ*ogcA* Tn7-*ogcI* with compared to without induction*.*** Fisher exact tests outputs assessing the co-occurrence of categorical assignments (statistically significant changes to proteins observed based on GO terms) with and without induction.

**Supplementary Table 15. DDA protein level LFQ analysis of *B. cenocepacia* Δ*ogcI* Tn7-*ogcI* strains containing p*ogcX* with and without induction.** The Perseus processed MSfragger search results for the protein analysis of four biological replicates of *B. cenocepacia* Δ*ogcI* Tn7-*ogcI,* Δ*ogcI*Δ*ogcX* Tn7-*ogcI, B. cenocepacia* Δ*ogcI*Δ*ogcA* Tn7-*ogcI and B. cenocepacia* Δ*ogcI*Δ*ogcB* Tn7-*ogcI* containing pCumate-*ogcX-his* with and without induction (1% Rhamnose, 100 μM cumate) are provided. For each identified protein, the log2 LFQ protein values, the ANOVA test including -log_10_(*p*-value) and if the proteins with *p*-values below the multiple hypothesis corrected *p*-value are provided. For each protein the top peptide probability, total number of MS/MS events for the corresponding protein as well as both the imputed and non-imputed data is provided.

**Supplementary Table 16. DDA PSM summary of *B. cenocepacia* Δ*ogcI* Tn7-*ogcI* strains containing p*ogcX* with and without induction*.*** The MSfragger PSMs search summary for the proteome analysis of *B. cenocepacia* Δ*ogcI* Tn7-*ogcI,* Δ*ogcI*Δ*ogcX* Tn7-*ogcI, B. cenocepacia* Δ*ogcI*Δ*ogcA* Tn7-*ogcI and B. cenocepacia* Δ*ogcI*Δ*ogcB* Tn7-*ogcI* containing pCumate-*ogcX-his* with and without induction is provided. For each modification type identified within the proteome the total number of PSMs assigned are tabulated.

**Supplementary Table 17. DDA protein level LFQ analysis of *B. cenocepacia* Δ*ogcAB* containing pSCrhaB2, pSCrhaB2-*ogcAB*-Met1, or pSCrhaB2-*ogcAB*-Met2.** The Perseus processed MSfragger search results for the protein analysis of four biological replicates of strains *B. cenocepacia* Δ*ogcAB* containing the plasmids pSCrhaB2, pSCrhaB2-*ogcAB*-Met1, or pSCrhaB2-*ogcAB*-Met2 without and with induction. For each identified protein, the log2 LFQ protein values, the T-test information including the -log_10_(*p*-value), differences in the mean between the groups and if the resulting *p*-values is below the multiple hypothesis corrected *p*-value are provided. For each protein the top peptide probability, total number of MS/MS events for the corresponding protein as well as both the imputed and non-imputed data is provided.

**Supplementary Table 18. DDA PSMs summary of *B. cenocepacia* Δ*ogcAB* containing pSCrhaB2, pSCrhaB2-*ogcAB*-Met1, or pSCrhaB2-*ogcAB*-Met2 with and without induction*.*** The MSfragger PSM search summary for whole proteome analysis of *B. cenocepacia* Δ*ogcI* Δ*ogcAB* Tn7-*ogcI* containing pSCrhaB2, pSCrhaB2-*ogcAB*-Met1, or pSCrhaB2-*ogcAB*-Met2 strains are provided. For each modification type identified within the proteome the total number of PSMs assigned are tabulated.

**Supplementary Table 19. DDA protein level LFQ analysis of *B. cenocepacia* Δ*ogcB* containing pSCrhaB2 and pSCrhaB2-*ogcB*.** The Perseus processed MSfragger search results for the protein analysis of four biological replicates of strains *B. cenocepacia* Δ*ogcB* containing the plasmids pSCrhaB2 and pSCrhaB2-*ogcB* grown with 0.05% rhamnose induction. For each identified protein, the log2 LFQ protein values, the T-test information including the -log_10_(*p*-value), differences in the mean between the groups and if the resulting *p*-values is below the multiple hypothesis corrected *p*-value are provided. For each protein the top peptide probability, total number of MS/MS events for the corresponding protein as well as both the imputed and non-imputed data is provided.

**Supplementary Table 20. DDA PSM summary of *B. cenocepacia* Δ*ogcB* containing pSCrhaB2 and pSCrhaB2-*ogcB* with induction*.*** The MSfragger PSM search summary for proteome analysis of *B. cenocepacia* Δ*ogcB* containing pSCrhaB2 and pSCrhaB2-*ogcB* strains are provided. For each modification type identified within the proteome the total number of PSMs assigned are tabulated.

**Supplementary Table 21. DDA protein level LFQ analysis of *B. cenocepacia* Δ*ogcI* containing pSCrhaB2 and pSCrhaB2-*ogcI*.** The Perseus processed MSfragger search results for the protein analysis of four biological replicates of strains *B. cenocepacia* Δ*ogcI* containing the plasmids pSCrhaB2 and pSCrhaB2-*ogcI* grown with 0.05% rhamnose induction. For each identified protein, the log2 LFQ protein values, the T-test information including the -log_10_(*p*-value), differences in the mean between the groups and if the resulting *p*-values is below the multiple hypothesis corrected *p*-value are provided. For each protein the top peptide probability, total number of MS/MS events for the corresponding protein as well as both the imputed and non-imputed data is provided.

**Supplementary Table 22. DDA PSM summary of *B. cenocepacia* Δ*ogcI* containing pSCrhaB2 and pSCrhaB2-*ogcI* with induction*.*** The MSfragger PSM search summary for proteome analysis of *B. cenocepacia* Δ*ogcI* containing pSCrhaB2 and pSCrhaB2-*ogcI* strains are provided. For each modification type identified within the proteome the total number of PSMs assigned are tabulated.

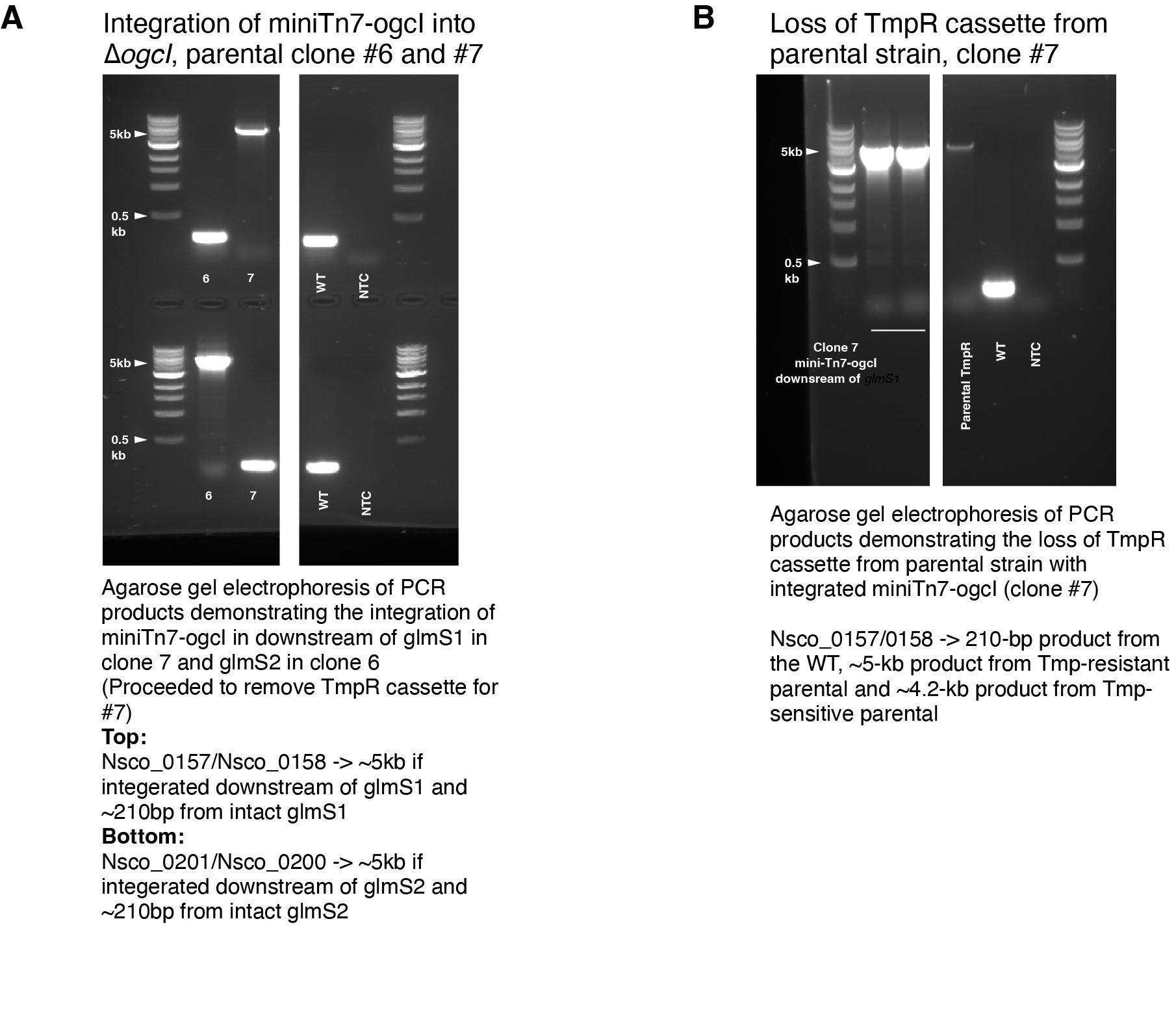

**Supplementary Figure 1. Generation of *B. cenocepacia* Δ*ogcI* Tn7-*ogcI.* A)** Integration of miniTn7-rha-ogcI into *B. cenocepacia* K65-2 Δ*ogcI*. Agarose gel electrophoresis of PCR products using two primer sets (Nsco_0157/Nsco_0158 and Nsco_0201/Nsco_0200) demonstrates the integration of miniTn7-rha-ogcI downstream of *glmS1* in clone #7 and *glmS2* in clone #6. **Top row**: PCR amplification with Nsco_0157/Nsco_0158 yields a ~5 kb product when miniTn7-rha-ogcI integrates downstream of *glmS1*; in the absence of integration, a ~210 bp product is observed. **Bottom row**: PCR amplification with Nsco_0201/Nsco_0200 yields a ~5 kb product when miniTn7-rha-ogcI integrates downstream of *glmS2*; if integration is absent, a ~210 bp product is observed. Clone #7, was selected for subsequent removal of the trimethoprim resistance (Tmp^R^) cassette from the miniTn7 element. **B)** Removal of trimethoprim resistance (Tmp^R^) cassette from *B. cenocepacia* K65-2 Δ*ogcI* Tn7-*ogcI*. Agarose gel electrophoresis of PCR products using primers Nsco_0157 and Nsco_0158 demonstrates the loss of the Tmp^R^ cassette from strain clone #7. PCR amplification with Nsco_0157/Nsco_0158 yields a ~5 kb product from the Tmp^R^ clone #7, a ~4.2 kb product from the Tmp-sensitive (Tmp^S^) clone #7 and a ~210 bp product from the WT strain.

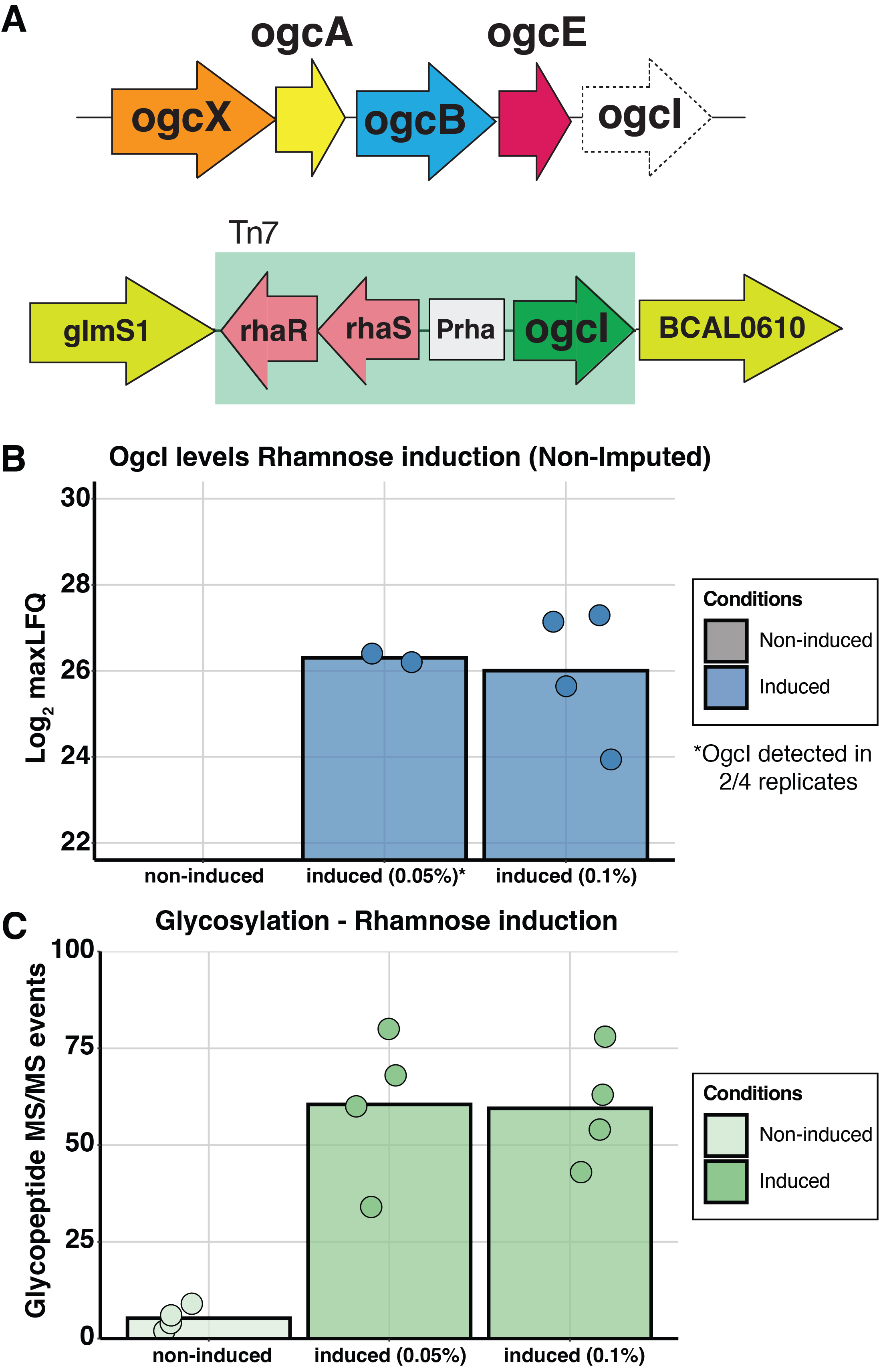

**Supplementary Figure 2 Proteomic/glycoproteomic validation of *B. cenocepacia* Δ*ogcI* Tn7-*ogcI*. A)** Graphic representation of the *ogc* cluster (BCAL3114–BCAL3118) demonstrating the loss of *ogcI* (BCAL3118) and integration of rhamnose-inducible *ogcI* downstream of *glmS1*. Before generating *ogc* mutants, inducible expression and restoration of glycosylation were confirmed by proteomic analysis. Induction of *ogcI* expression by 0.05% and 0.1% rhamnose led to the restoration of both **B)** OgcI and **C)** Glycosylation, as determined by identified glycopeptides, in the strain *B. cenocepacia* Δ*ogcI* Tn7-*ogcI*.

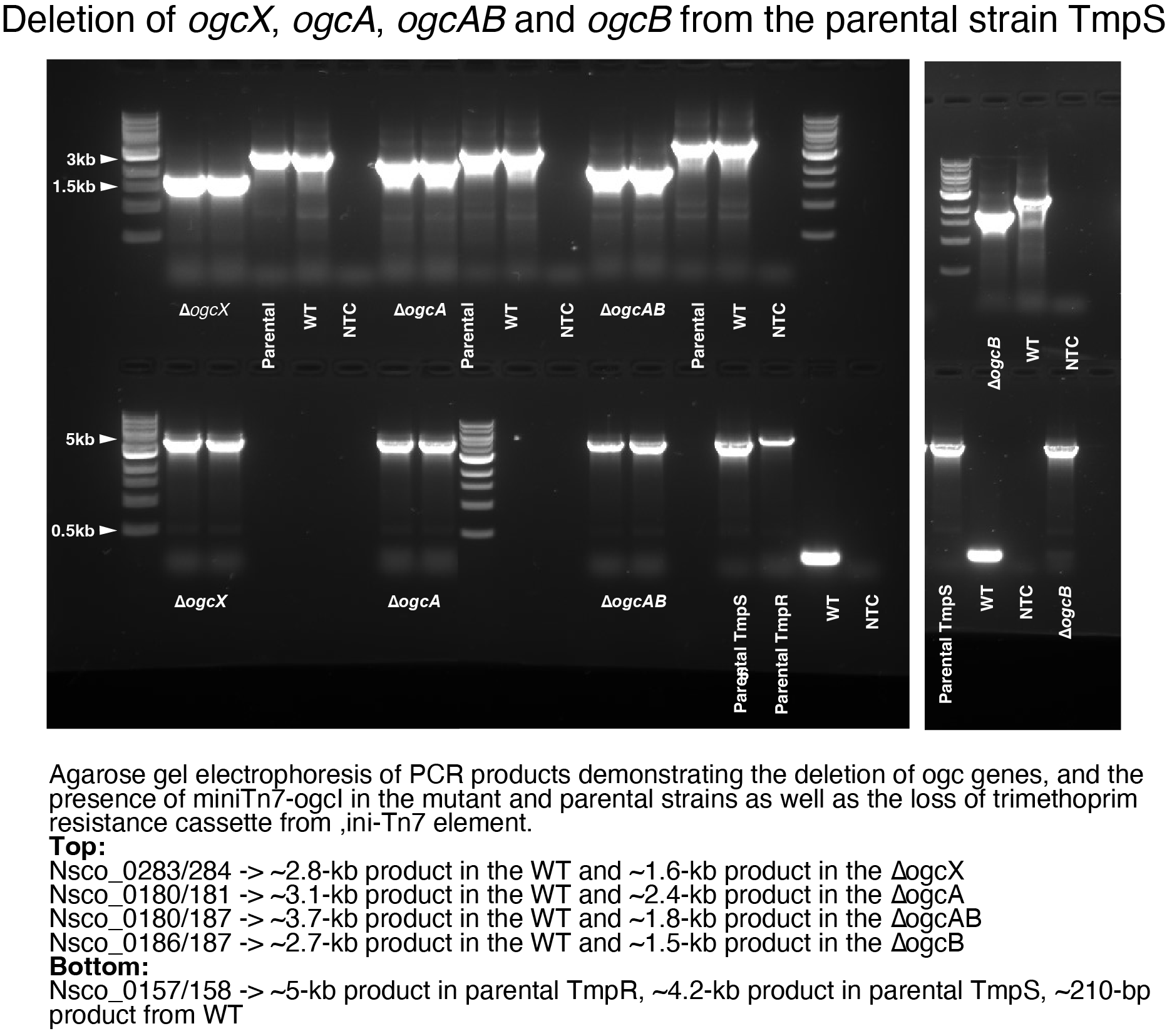

**Supplementary Figure 3. Deletion of *ogcX*, *ogcA*, *ogcAB*, and *ogcB* from the Tmp^S^ *B. cenocepacia* K65-2 Δ*ogcI* Tn7-*ogcI*.** Agarose gel electrophoresis of PCR products demonstrating both the deletion of ogc genes and the presence of miniTn7-rha-*ogcI* in mutant and parental strains, as well as the loss of the trimethoprim resistance (Tmp^R^) cassette from the miniTn7 element. Top row: PCR amplification with Nsco_0283/Nsco_0284 yields a ~2.8 kb product in the wild-type (WT) and a ~1.6 kb product in the mutant lacking *ogcX*. PCR amplification with Nsco_0180/Nsco_0181 yields a ~3.1 kb product in the WT and a ~2.4 kb product in the mutant lacking *ogcA*. PCR amplification with Nsco_0180/Nsco_0187 yields a ~3.7 kb product in the WT and a ~1.8 kb product in the ∆*ogcAB* mutant. PCR amplification with Nsco_0186/Nsco_0187 yields a ~2.7 kb product in the WT and a ~1.5 kb product in the mutant lacking *ogcB*. Bottom row: PCR amplification with Nsco_0157/Nsco_0158 yields a ~5 kb product in the Tmp^R^ *B. cenocepacia* K65-2 Δ*ogcI* Tn7-*ogcI* strain, a ~4.2 kb product in the Tmp^S^ *B. cenocepacia* K65-2 Δ*ogcI* Tn7-*ogcI* strain, and a ~210 bp product from the WT strain.

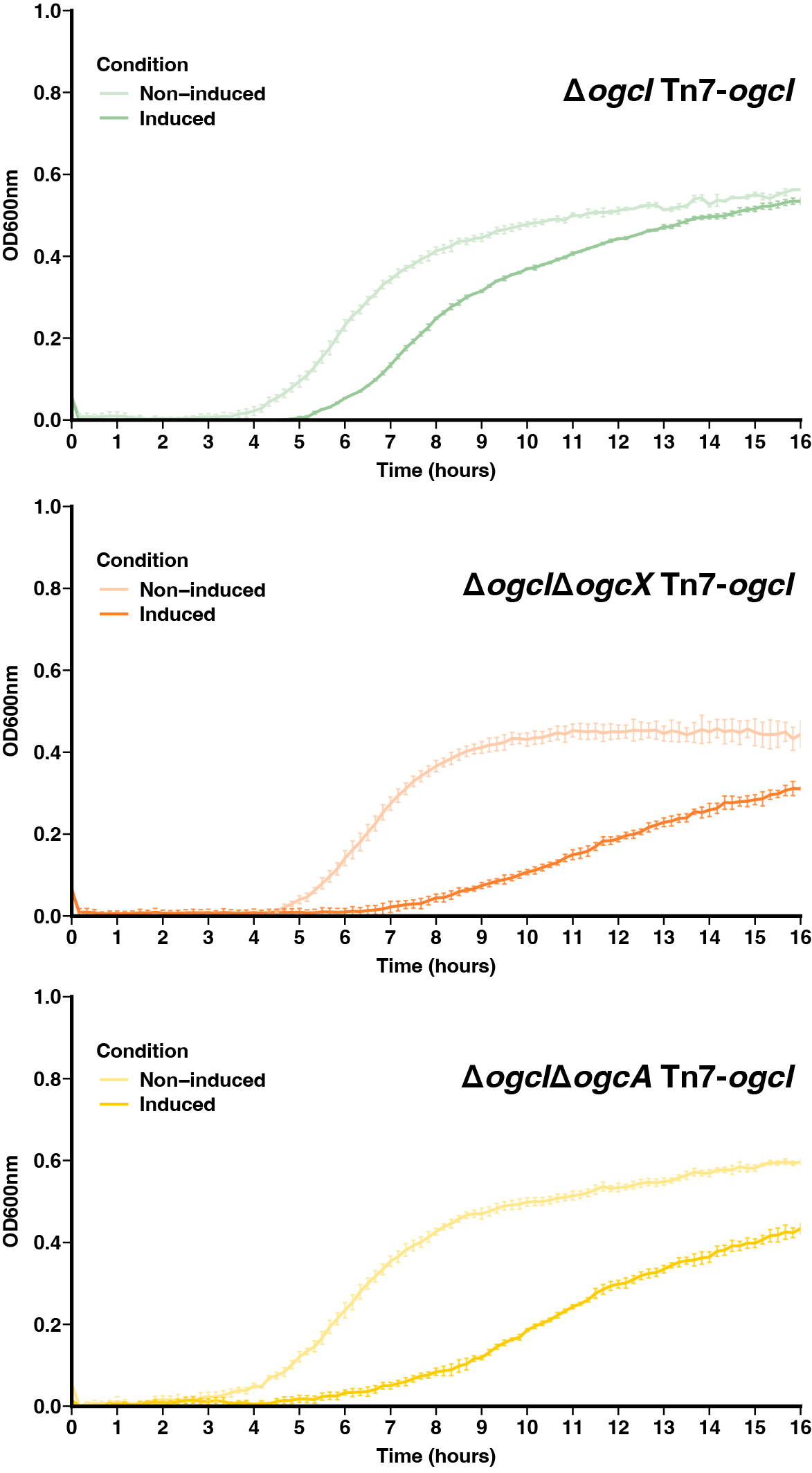

**Supplementary Figure 4. Plate-based growth assays of Δ*ogcI*Δ*ogcX* Tn7-*ogcI*, Δ*ogcI*Δ*ogcA* Tn7-*ogcI*, and Δ*ogcI* Tn7-*ogcI*.** Upon initiation of glycosylation with 1% rhamnose the absence of *ogcX* and *ogcA* leads to a reduced growth rate compared to Δ*ogcI* Tn7-*ogcI*. For Δ*ogcI* Tn7-*ogcI* the presence of 1% rhamnose delayed grow but a comparable rate is observed compared to both Δ*ogcI*Δ*ogcX* Tn7-*ogcI*, Δ*ogcI*Δ*ogcA* Tn7-*ogcI*. Growth curves of the strains were generated by growing bacteria in LB broth in the absence (non-induced) or presence of rhamnose (induced) and measuring OD_600_ using a CLARIOstar microplate reader.

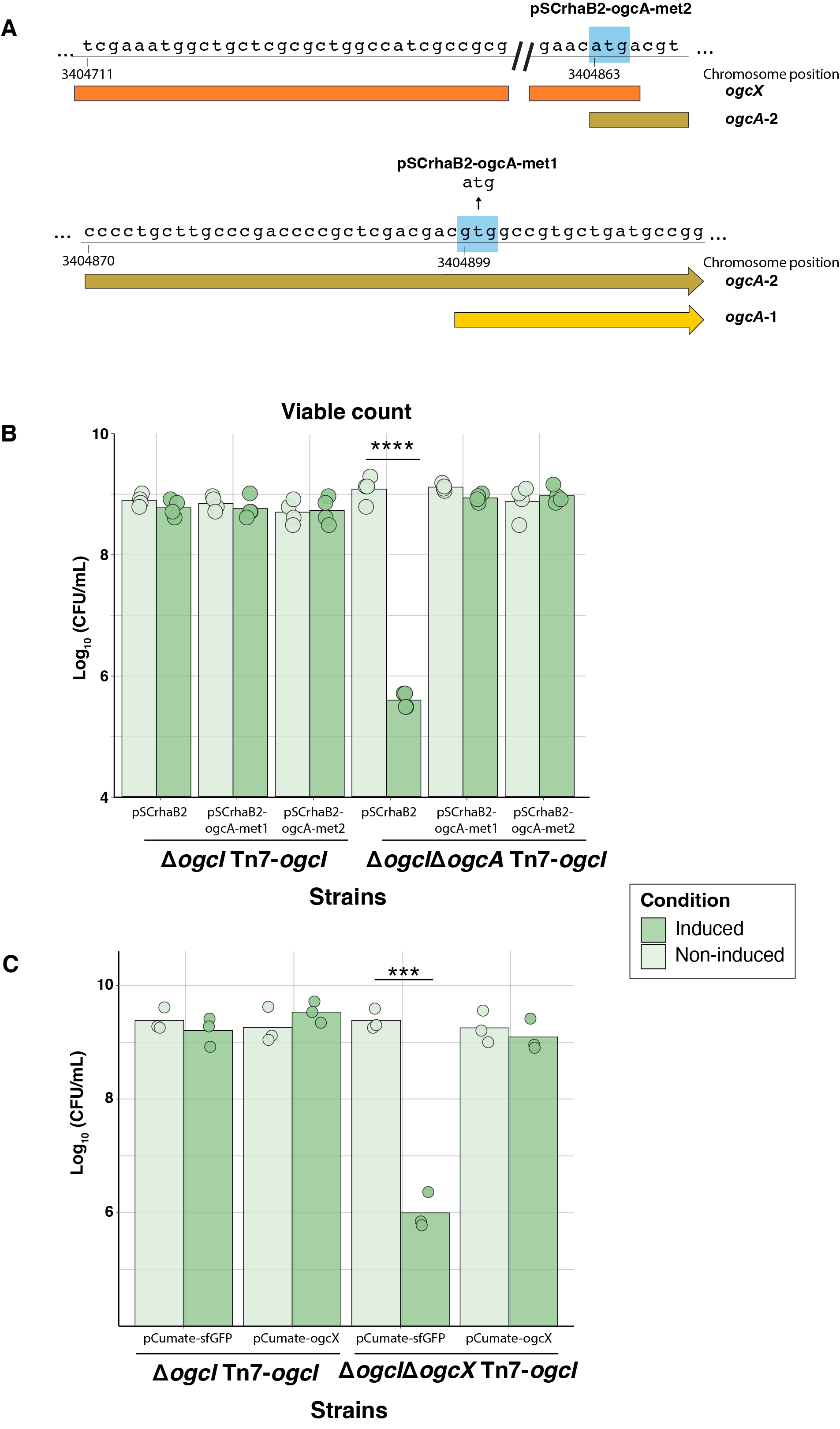

**Supplementary Figure 5. Complementation of *B. cenocepacia* Δ*ogcI* Δ*ogcA* Tn7-*ogcI.*** Graphic representation of putative start sites of *ogcA*. Due to the lack of *ogcA* expression from the annotated start codon, the gene was cloned into the pSCrhaB2 plasmid from two alternative start codons: a GTG (valine) converted to ATG, and a start codon located further upstream were designated as *ogcA*-Met1 and *ogcA*-Met2, respectively.

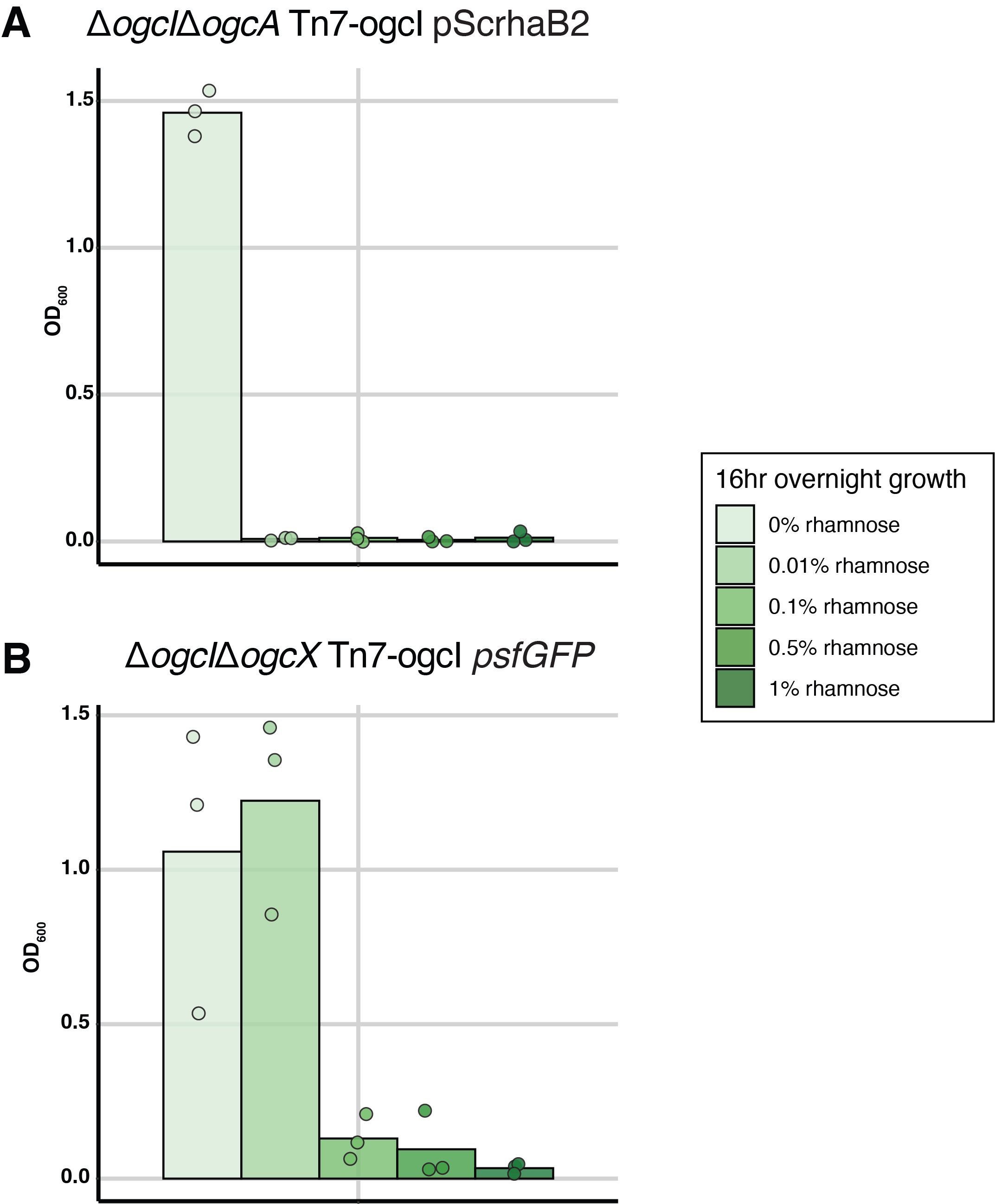

**Supplementary Figure 6. Viability of *B. cenocepacia* Δ*ogcI*Δ*ogcA* Tn7-*ogcI* and Δ*ogcI*Δ*ogcX* Tn7-ogcI containing control plasmids in response to induction. A)** Overnight growth measurements of Δ*ogcI*Δ*ogcA* Tn7-*ogcI* carrying pSCrhaB2 induced under different concentrations of rhamnose reveal that the induction of OgcI under growth conditions to maintain plasmids results in a loss of viability. **B)** Overnight growth measurements of Δ*ogcI*Δ*ogcX* Tn7-*ogcI* carrying *psfGFP* induced under different concentrations of rhamnose reveal that the induction of OgcI under growth conditions to maintain plasmids results in a loss of viability at rhamnose concentrations above 0.01%.

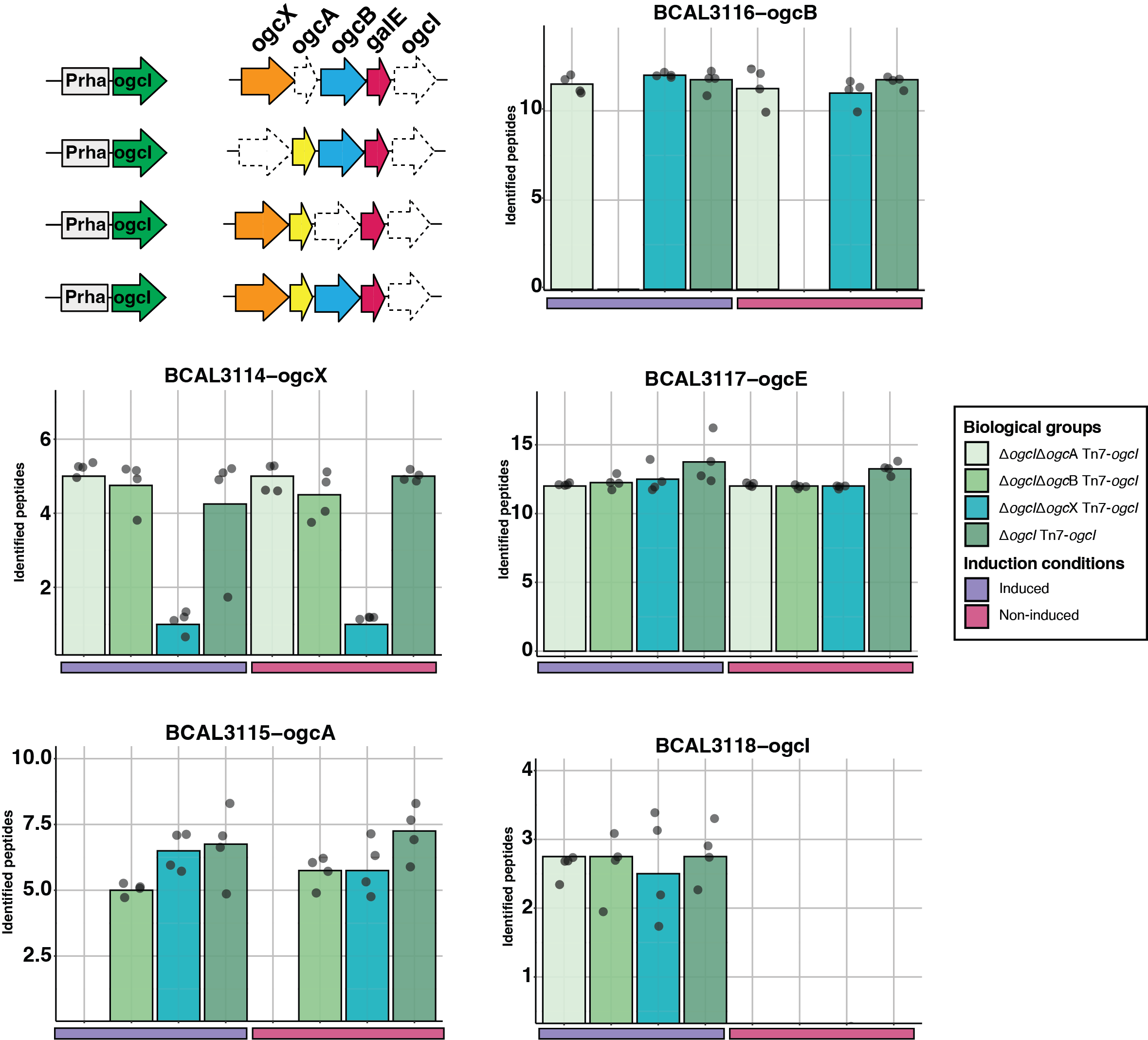

**Supplementary Figure 7. Proteomic analysis of the proteins of the *ogc* within Δ*ogcI*Δ*ogcX* Tn7-*ogcI*, Δ*ogcI*Δ*ogcA* Tn7-*ogcI*, Δ*ogcI*Δ*ogcB* Tn7-*ogcI* and Δ*ogcI* Tn7-*ogcI* strains.** DIA proteomic analysis demonstrates the selective absence of the OGC proteins within mutants and confirms that these mutations are non-polar, with the abundance of undisrupted OGC proteins equivalent to that in the parental strain (Δ*ogcI* Tn7-*ogcI*).

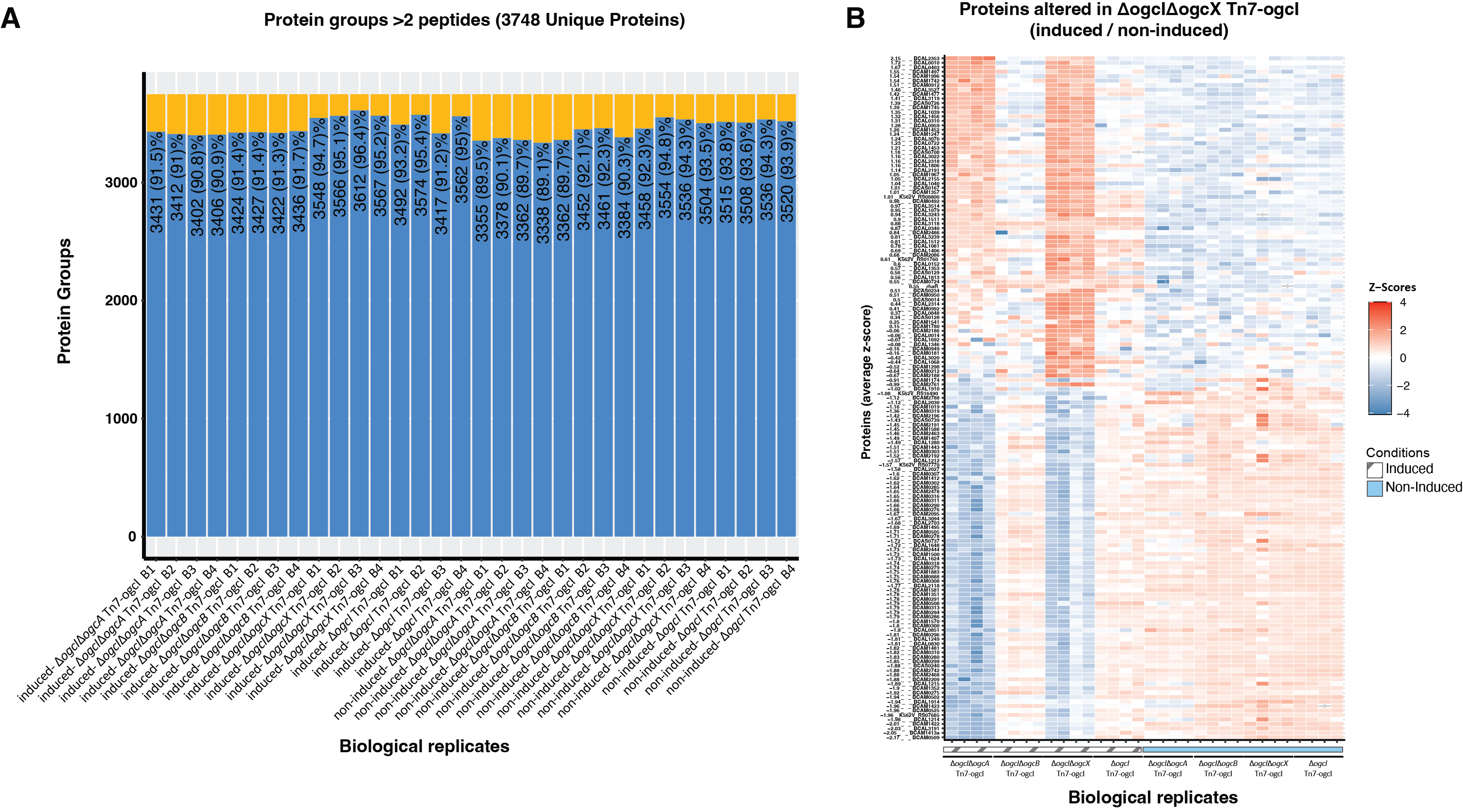

**Supplementary Figure 8. Proteomic analysis of *B. cenocepacia* Δ*ogcI*** **Tn7-*ogcI* strains. A)** Proteomic coverage of *B. cenocepacia* proteins observed across biological replicates demonstrate that >90% of all proteins were identified within samples with at least two unique precursors**. B)** Heatmap of proteomic changes associated with Z-score > ±2 in at least one biological group reveals similar alterations observed within Δ*ogcI*Δ*ogcX* Tn7-*ogcI* and Δ*ogcI*Δ*ogcA* Tn7-*ogcI* upon induction of OgcI.

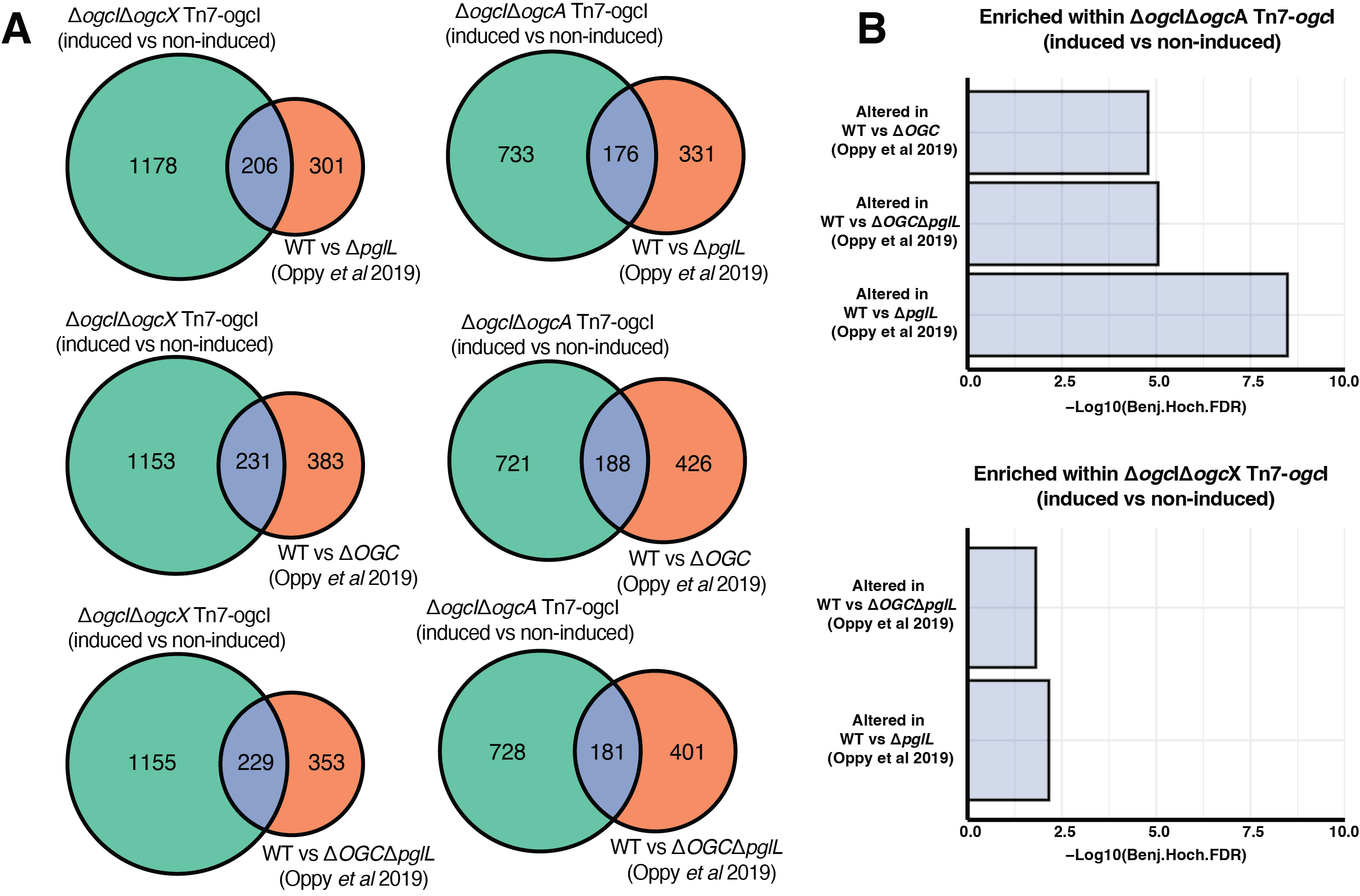

**Supplementary Figure 9. Comparison of proteomic alterations observed within *B. cenocepacia* Δ*ogc*IΔ*ogc*X Tn7-*ogc*I and Δ*ogc*IΔ*ogc*A Tn7-ogcI compared to *B. cenocepacia* glycosylation-null strains. A)** Venn diagrams showing the overlap in proteomic changes observed within Δ*ogc*IΔ*ogc*X Tn7-*ogc*I and Δ*ogc*IΔ*ogc*A Tn7-ogcI upon induction, compared to previously reported proteomic alterations in *B. cenocepacia* glycosylation-null strains Δ*pgl*L, Δ*ogc*, and Δ*pgl*LΔ*ogc*, relative to *B. cenocepacia* K56-2 WT ^3^. **B)** Enrichment analysis of the overlap in altered proteins within Δ*ogc*IΔ*ogc*X Tn7-ogcI and Δ*ogc*IΔ*ogc*A Tn7-ogcI upon induction, compared to previously reported proteomic alterations in the *B. cenocepacia* glycosylation-null strains Δ*pgl*L, Δ*OGC*, and Δ*pgl*LΔ*OGC*, relative to *B. cenocepacia* K56-2 WT, reveals a statistically significant enrichment. These findings support similar proteomic alterations between the induced strains and the glycosylation-null strains.

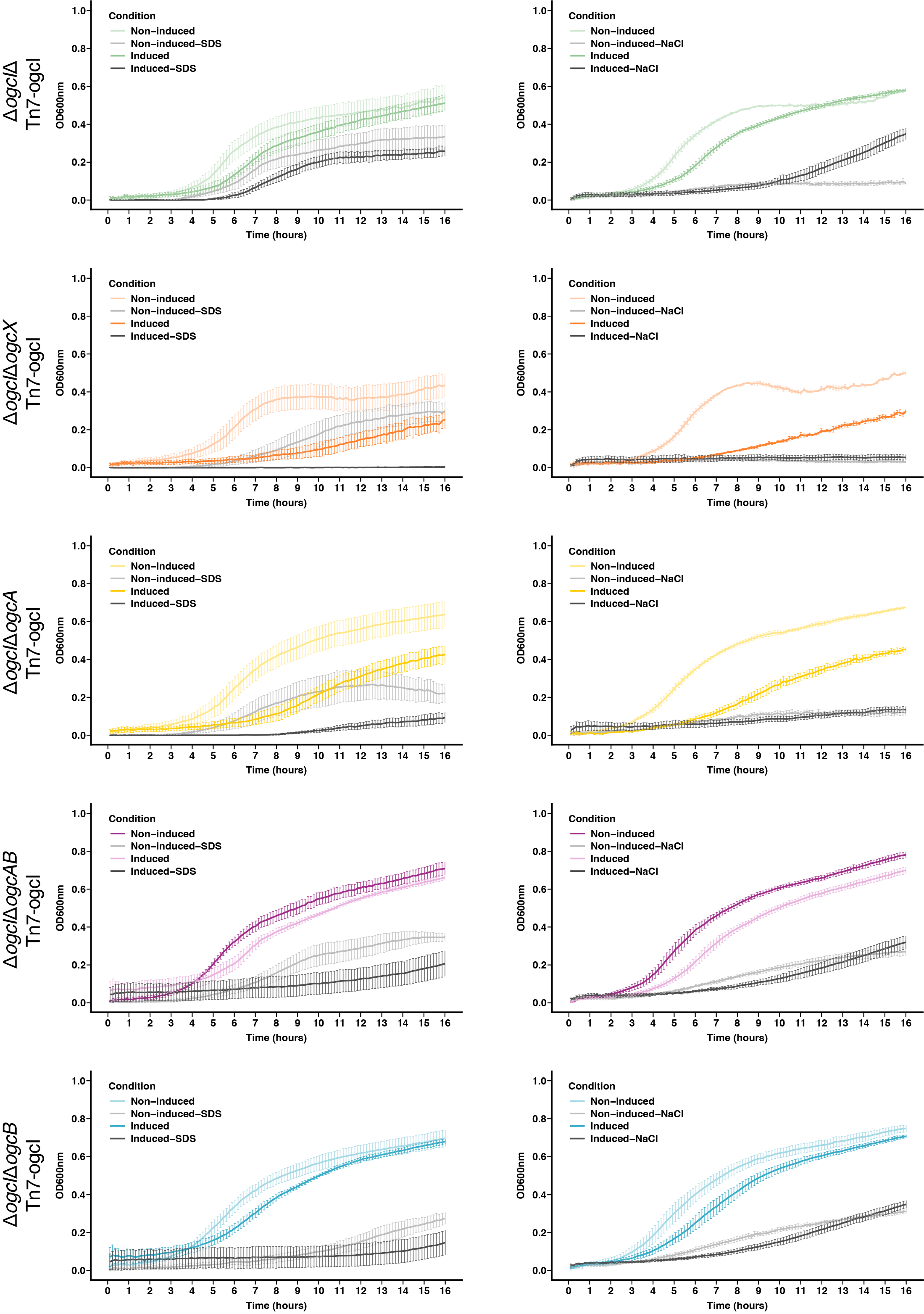

**Supplementary Figure 10. Plate-based growth assays of Δ*ogcI* Tn7-*ogcI,* Δ*ogcI*Δ*ogcX* Tn7-*ogcI*, Δ*ogcI*Δ*ogcA* Tn7-*ogcI*, Δ*ogcI*Δ*ogcAB* Tn7-*ogcI* and Δ*ogcI*Δ*ogcB* Tn7-*ogcI* strains in the presence of membrane / osmotic stress agents.** Growth curves of the strains subjected to membrane stress (0.01% SDS) and osmotic stress (2% NaCl) with and without 1% rhamnose induction. The growth curves demonstrate that the deletion of *ogcX* and *ogcA* results in nearly total loss of growth in the presence of 0.01% SDS when glycosylation is initiated. In the presence of osmotic stress (2% NaCl), both mutants lacking *ogcX* and *ogcA* failed to grow regardless of induction.

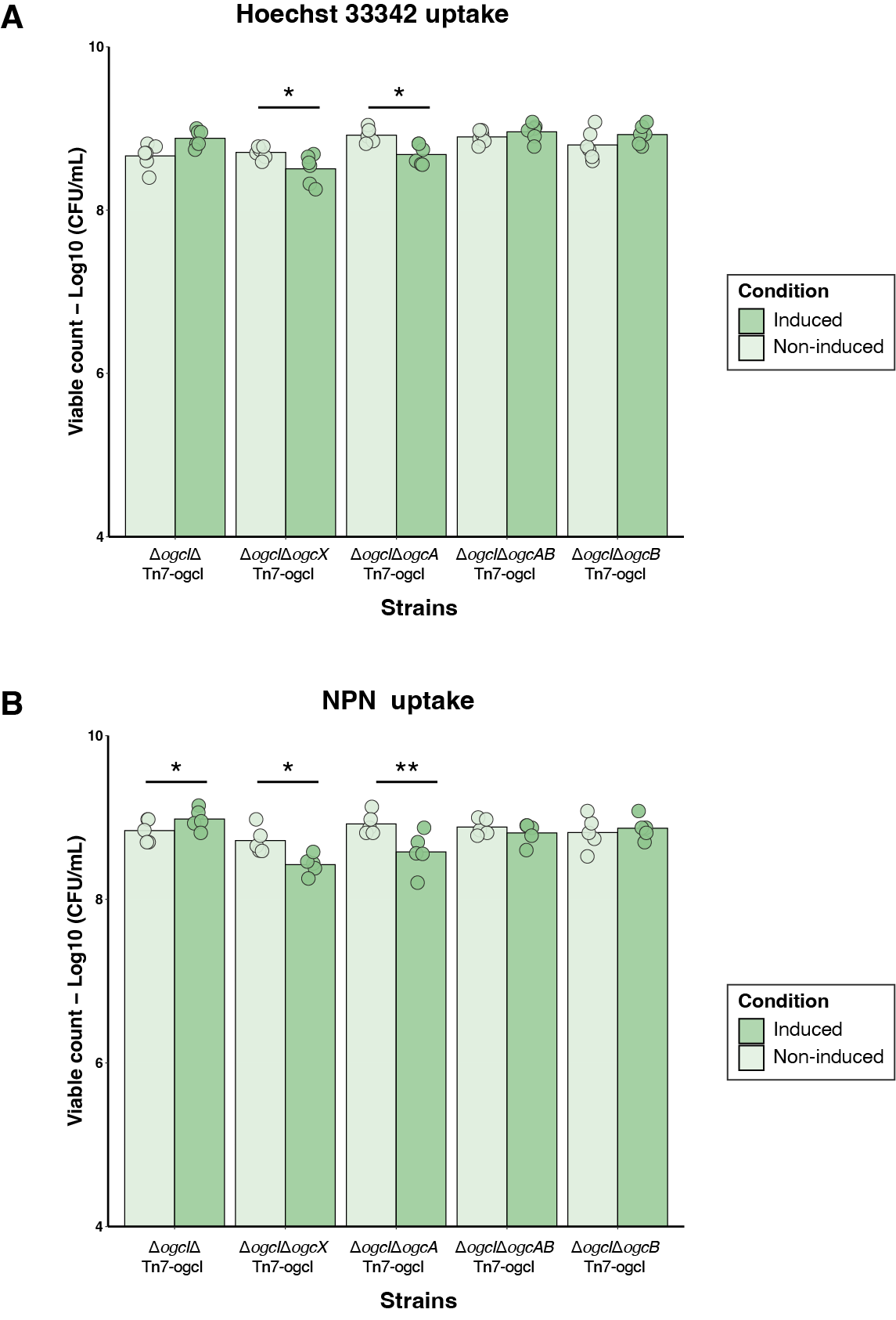

**Supplementary Figure 11. Viable counts of Hoechst and NPN uptake assays.** Viable counts of the parental strain and mutants demonstrate viability of strains prior to the addition of Hoechst 33342 and NPN dyes. Bacterial suspensions were serially diluted and spotted onto LB agar plates, and colonies counted after 48 hours of incubation at 37°C. Due to the presence of sodium azide in the buffer used for washing cells in the NPN assay, bacterial suspensions from the same overnight culture and OD, but resuspended in PBS, were used for determining the viable cell counts.

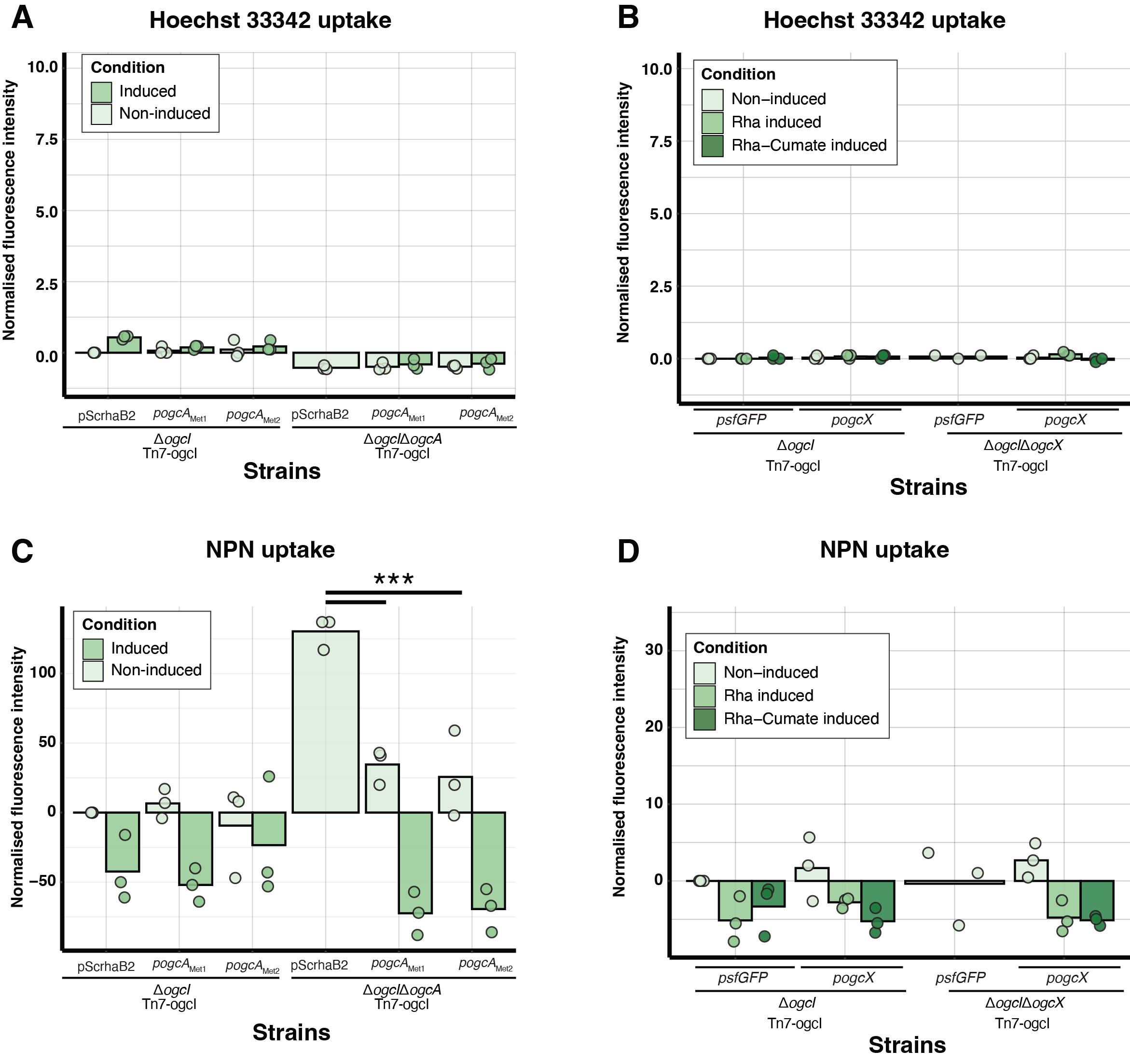

**Supplementary Figure 12. Hoechst and NPN uptake assays of complemented Δ*ogc*IΔ*ogc*A Tn7-*ogc*I and Δ*ogc*IΔ*ogc*X Tn7-*ogc*I. Hoechst 33342 (A/C) and NPN (B/D) uptake assays reveal that complementation of Δ*ogc*IΔ*ogc*A Tn7-*ogc*I and Δ*ogc*IΔ*ogc*X Tn7-*ogc*I reduces dye uptake to levels equivalent to Δ*ogc*I Tn7-*ogc*I containing plasmids (n=3 for Hoechst 33342, n=3 for NPN). The fluorescence intensities have been normalized against bacterial cell counts.**

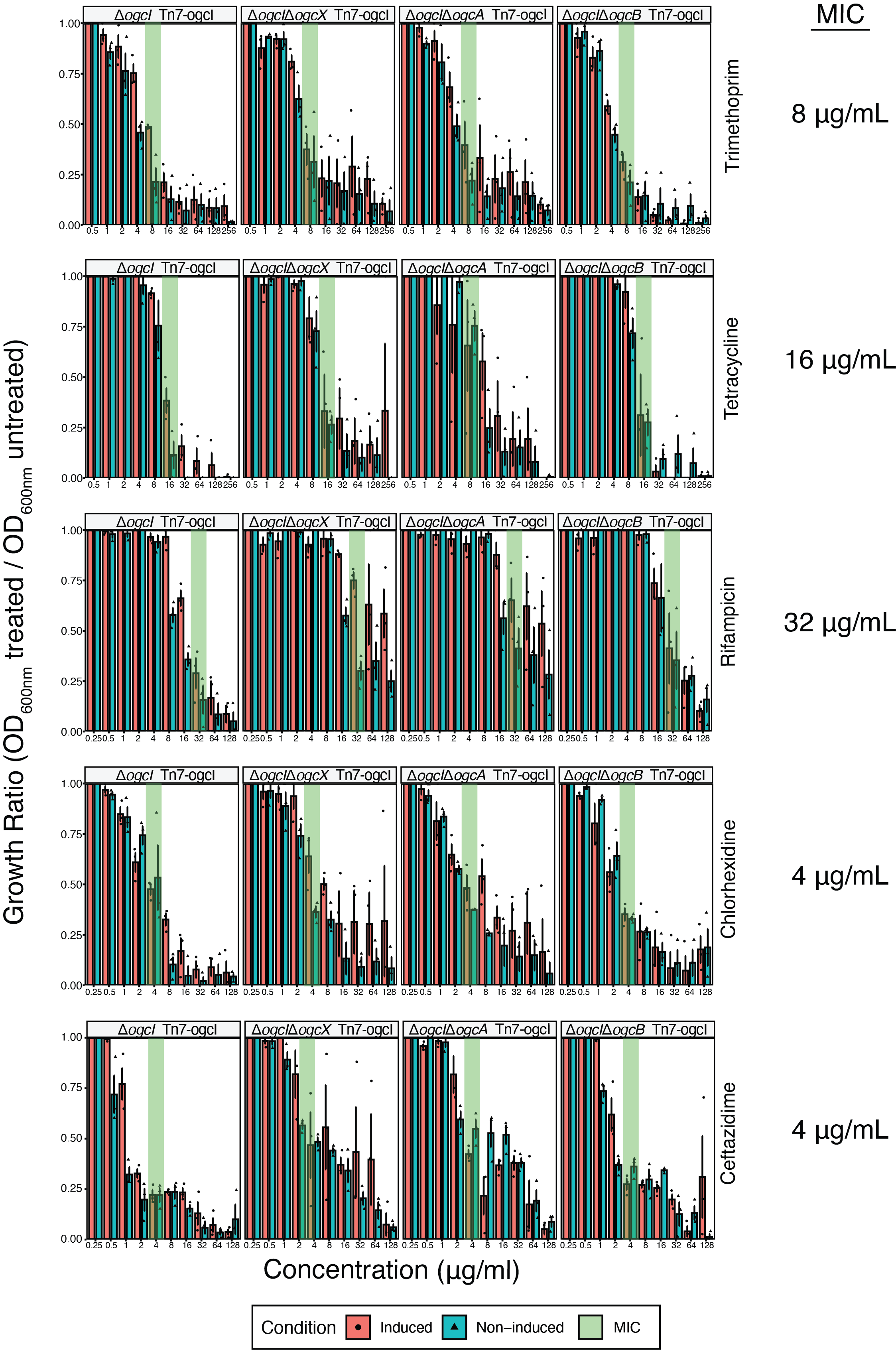

**Supplementary Figure 13. Antibiotic susceptibility assays of glycosylation inducible *B. cenocepacia* strains, with and without 1% rhamnose induction.** The growth of antibiotic-treated bacteria was measured by reading absorbance at 600 nm (OD_600nm_) and normalised to the OD_600nm_ of the lowest antibiotic concentration. Individual values (dots) represent normalised values for three biological replicates, and bars represent the mean of the three replicates.

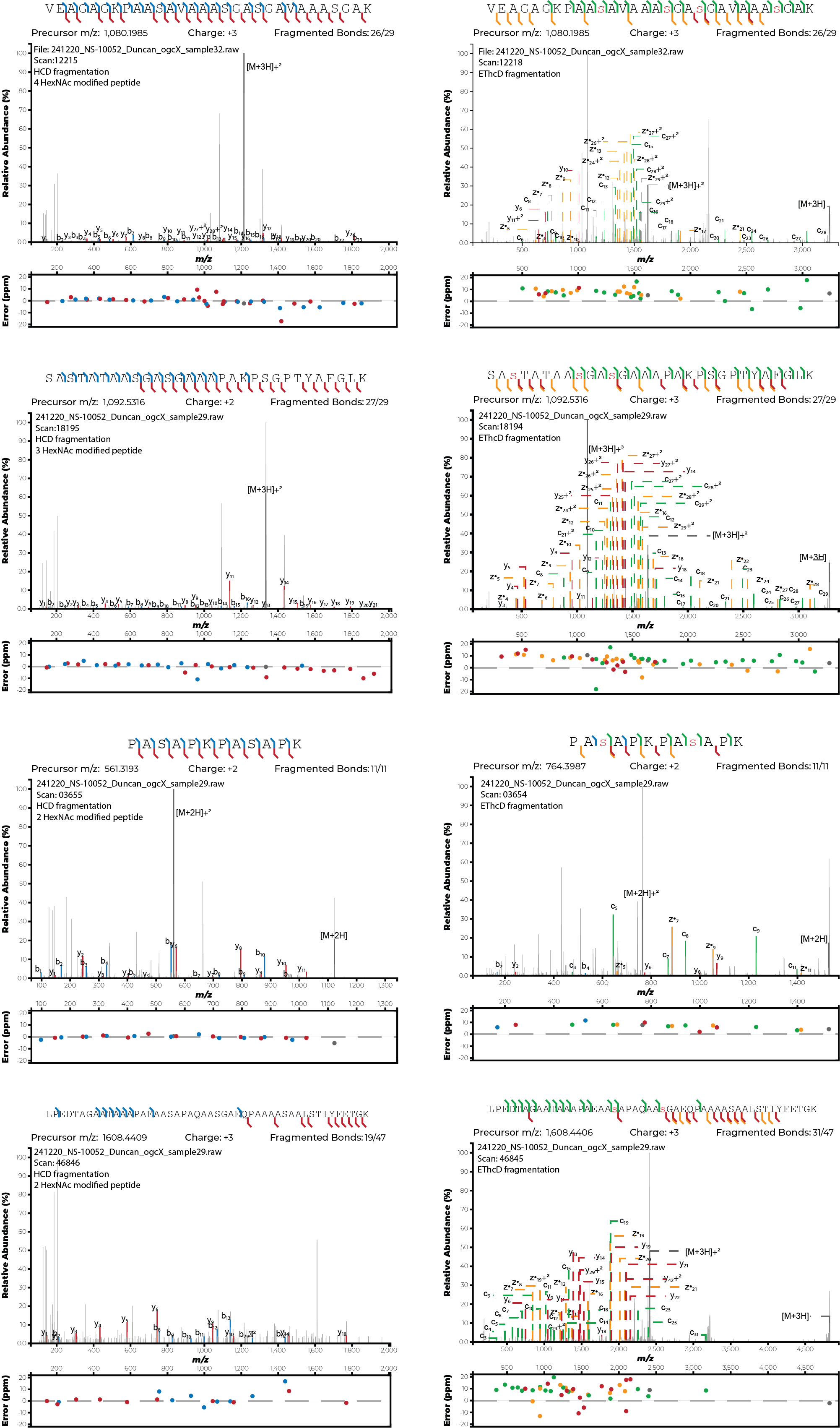

**Supplementary Figure 14. *B. cenocepacia* Δ*ogcI*Δ*ogcB* Tn7-*ogcI* Glycopeptides:** Manual inspection of EThcD and HCD glycopeptide spectra observed within *B. cenocepacia* Δ*ogcI*Δ*ogcB* Tn7-*ogcI* containing p*ogcX* demonstrates that assigned HexNAc2-modified glycopeptides correspond to peptides with multiple single HexNAc residues attached to distinct serine residues**.**

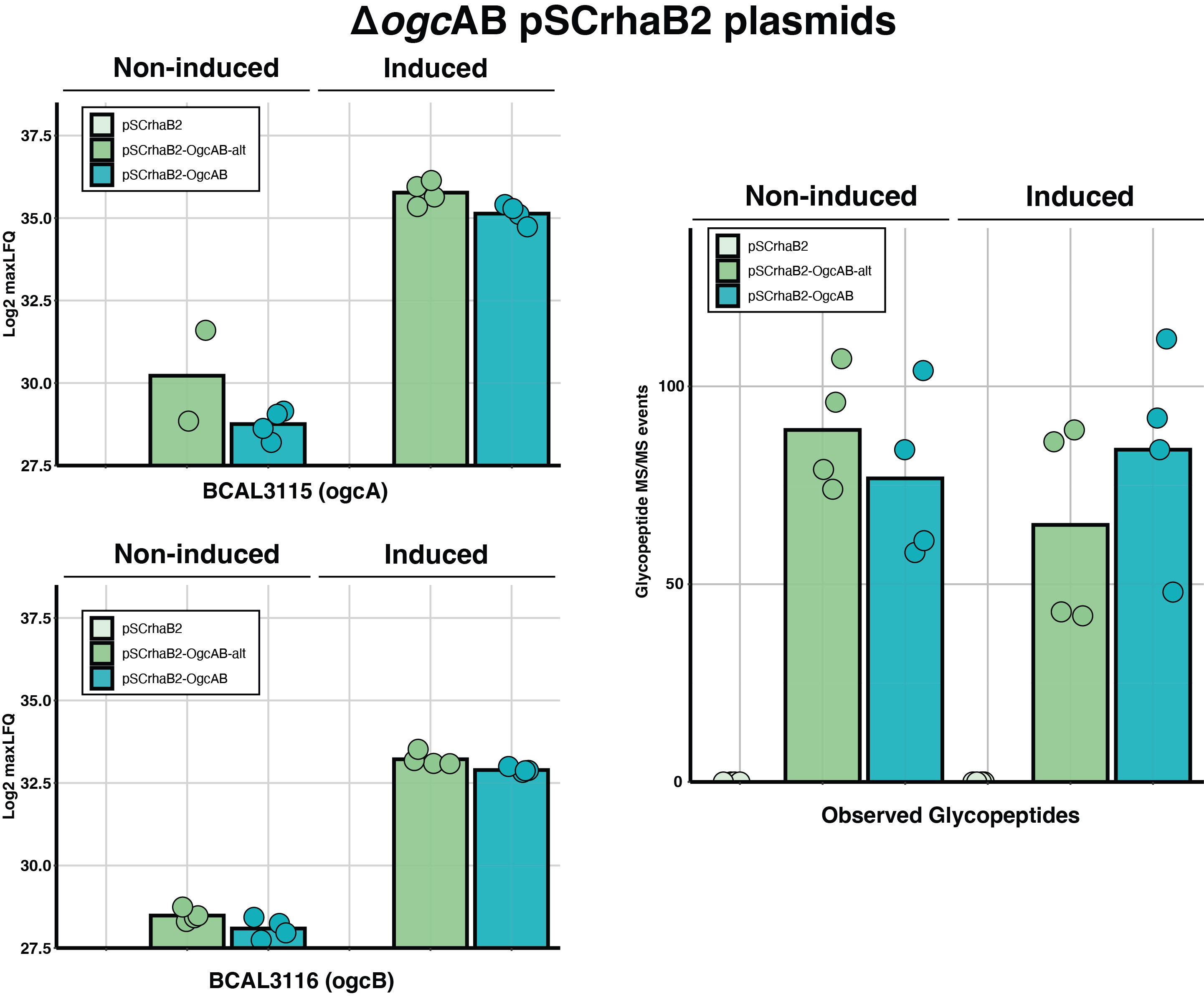

**Supplementary Figure 15. Proteomic analysis of pSCrhaB2-*ogcAB in* Δ*ogcAB*.** Quantification of OgcA and OgcB from whole-cell proteomic analysis of *B. cenocepacia* Δ*ogcAB* strains carrying empty pSCrhaB2 or pSCrhaB2-*ogcAB* constructs from two different start sites confirm restoration of OgcA and OgcB as well as glycosylation. Production of OgcA and OgcB was observed in the non-induced conditions with comparable levels of glycosylation observed to samples induced with 0.05% Rhamnose. Proteomic and glycopeptide data is provided in Supplementary Table 14 and 15.

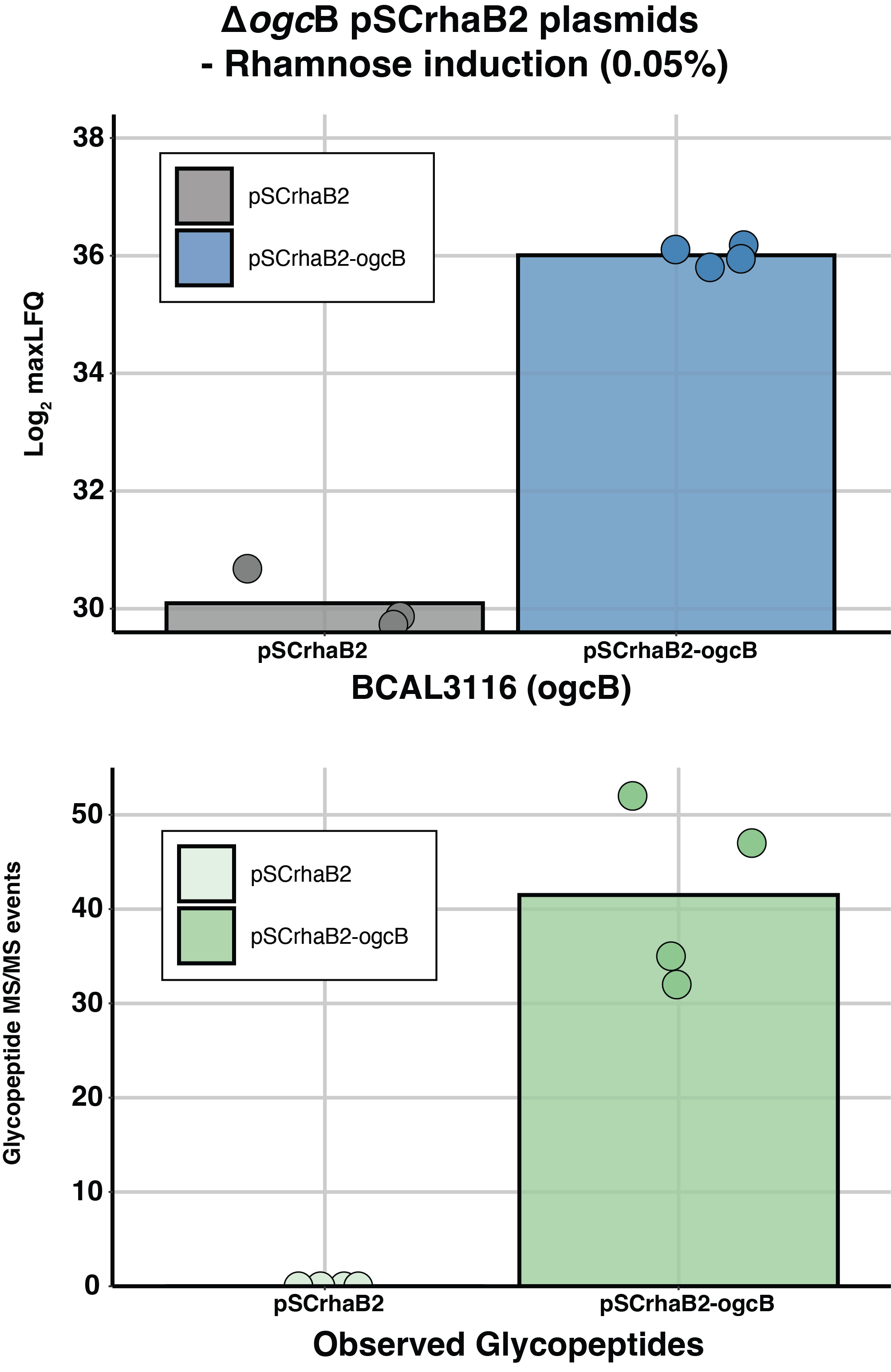

**Supplementary Figure 16. Proteomic analysis of pSCrhaB2-*ogcB* in Δ*ogcB*.** Quantification of OgcB from whole-cell proteomic analysis of *B. cenocepacia* Δ*ogcB* strains carrying empty pSCrhaB2 or pSCrhaB2-*ogcB* grown with 0.05% rhamnose induction. Induction resulted in elevated levels of OgcB within *B. cenocepacia* Δ*ogcB* carrying pSCrhaB2-*ogcB* and the restoration of glycosylation. Proteomic and glycopeptide data is provided in Supplementary Table 16 and 17.

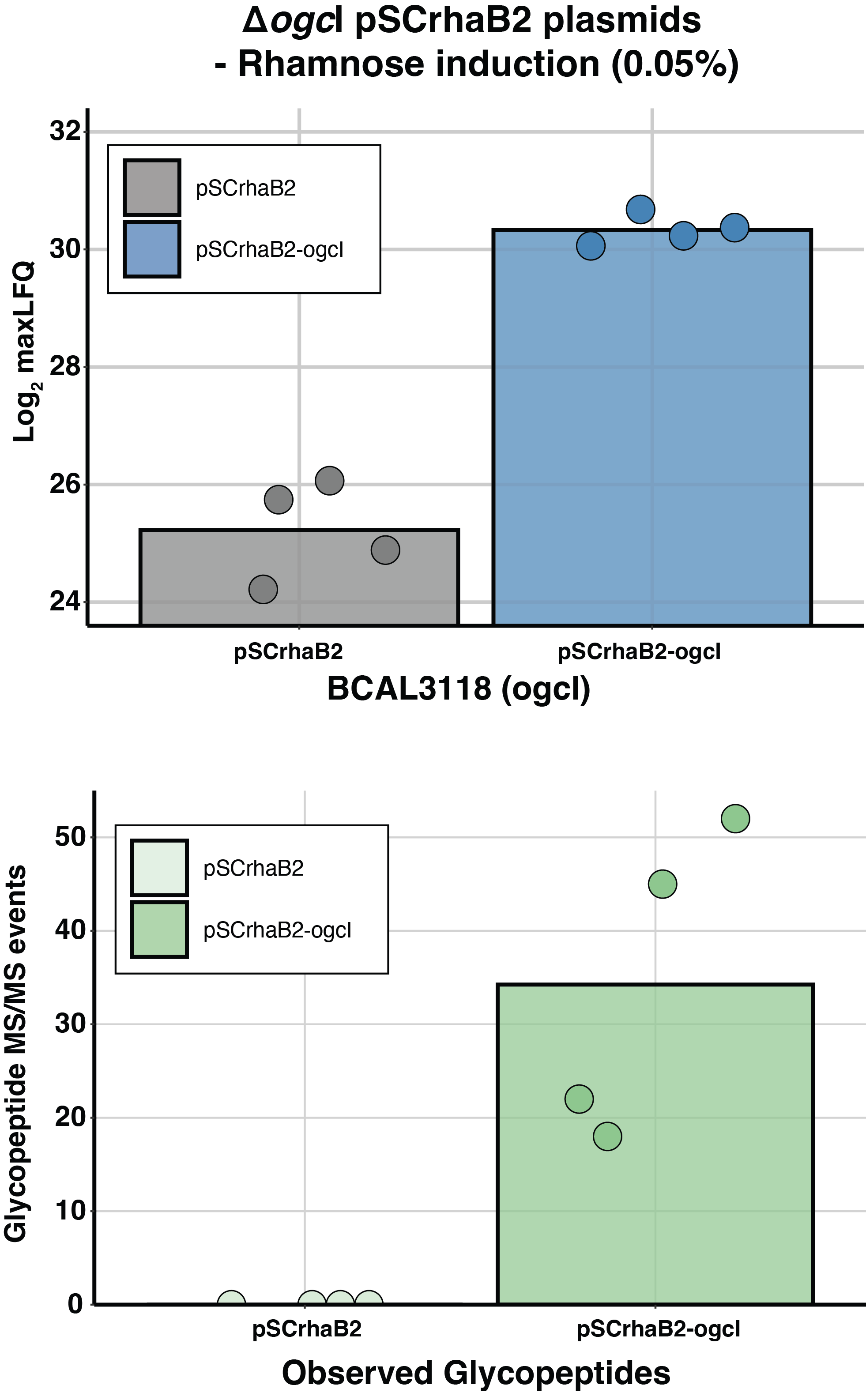

**Supplementary Figure 17. Proteomic analysis of pSCrhaB2-*ogcI* in Δ*ogcI*.** Quantification of OgcI from whole-cell proteomic analysis of *B. cenocepacia* Δ*ogcI* strains carrying empty pSCrhaB2 or pSCrhaB2-*ogcI* grown with 0.05% rhamnose induction. Induction resulted in elevated levels of OgcI within *B. cenocepacia* Δ*ogcI* carrying pSCrhaB2-*ogcI* and the restoration of glycosylation. Proteomic and glycopeptide data is provided in Supplementary Table 18 and 19.

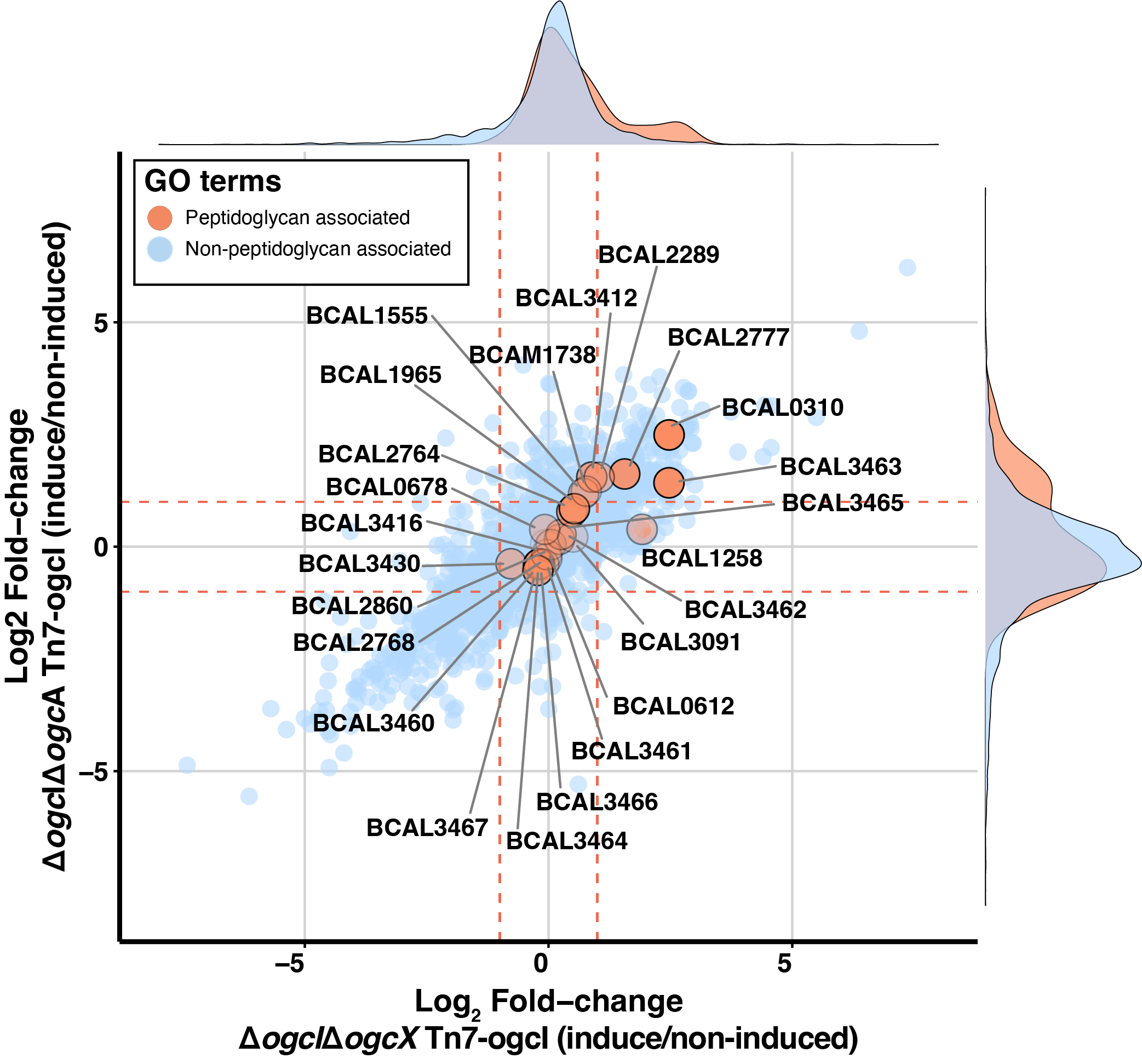

**Supplementary Figure 18. Proteomic analysis of Δ*ogcI*Δ*ogcA* Tn7-*ogcI* and Δ*ogcI*Δ*ogcX* Tn7-*ogcI* in response to glycosylation initiation.** DIA proteomic analysis of Δ*ogcI*Δ*ogcA* Tn7-*ogcI* and Δ*ogcI*Δ*ogcX* Tn7-*ogcI* strains with and without rhamnose induction, with proteins assigned to the GO terms GO:0042834, GO:0009253, GO:0009252, or GO:0000270 as denoted in Supplementary Table 13 highlighted in orange.

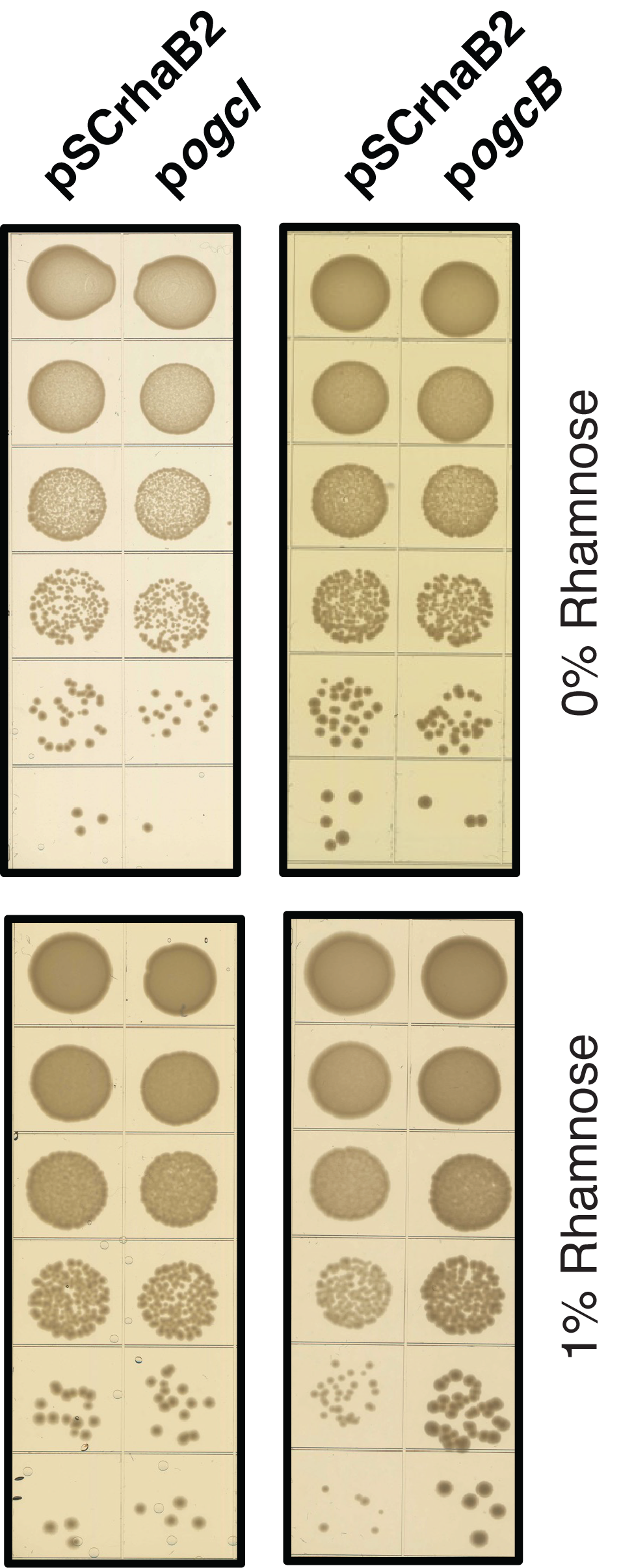

**Supplementary Figure 19. Spot plate assays of *E. coli* PIR2 containing the expression vectors pSCrhaB2, pSCrhaB2-*ogcI (*p*ogcI)* and pSCrhaB2-*ogcB* (p*ogcB*).** Induction of *ogc*I and *ogc*B with 1% rhamnose does not impact the growth of *E. coli.*

**References**

1	Lopez, C. M., Rholl, D. A., Trunck, L. A. & Schweizer, H. P. Versatile dual-technology system for markerless allele replacement in Burkholderia pseudomallei. *Applied and environmental microbiology* **75**, 6496-6503 (2009). <https://doi.org:10.1128/AEM.01669-09>

2	Darling, P., Chan, M., Cox, A. D. & Sokol, P. A. Siderophore production by cystic fibrosis isolates of Burkholderia cepacia. *Infection and immunity* **66**, 874-877 (1998).

3	Oppy, C. C. *et al.* Loss of O-linked protein glycosylation in Burkholderia cenocepacia impairs biofilm formation, siderophore activity and alters transcriptional regulators *mSphere* **4**, e00660-00619 (2019).

4	Figurski, D. H. & Helinski, D. R. Replication of an origin-containing derivative of plasmid RK2 dependent on a plasmid function provided in trans. *Proceedings of the National Academy of Sciences of the United States of America* **76**, 1648-1652 (1979). <https://doi.org:10.1073/pnas.76.4.1648>

5	Garcia, E. C., Anderson, M. S., Hagar, J. A. & Cotter, P. A. Burkholderia BcpA mediates biofilm formation independently of interbacterial contact-dependent growth inhibition. *Molecular microbiology* **89**, 1213-1225 (2013). <https://doi.org:10.1111/mmi.12339>

6	Hamad, M. A., Skeldon, A. M. & Valvano, M. A. Construction of aminoglycoside-sensitive Burkholderia cenocepacia strains for use in studies of intracellular bacteria with the gentamicin protection assay. *Applied and environmental microbiology* **76**, 3170-3176 (2010). <https://doi.org:10.1128/AEM.03024-09>

7	Choi, K. H., DeShazer, D. & Schweizer, H. P. mini-Tn7 insertion in bacteria with multiple glmS-linked attTn7 sites: example Burkholderia mallei ATCC 23344. *Nature protocols* **1**, 162-169 (2006). <https://doi.org:10.1038/nprot.2006.25>

8	Cardona, S. T. & Valvano, M. A. An expression vector containing a rhamnose-inducible promoter provides tightly regulated gene expression in Burkholderia cenocepacia. *Plasmid* **54**, 219-228 (2005). <https://doi.org:10.1016/j.plasmid.2005.03.004>

9	Flannagan, R. S., Linn, T. & Valvano, M. A. A system for the construction of targeted unmarked gene deletions in the genus Burkholderia. *Environ Microbiol* **10**, 1652-1660 (2008). <https://doi.org:10.1111/j.1462-2920.2008.01576.x>

10	Lefebre, M. D. & Valvano, M. A. Construction and evaluation of plasmid vectors optimized for constitutive and regulated gene expression in Burkholderia cepacia complex isolates. *Applied and environmental microbiology* **68**, 5956-5964 (2002).

11	Aubert, D. F., Hamad, M. A. & Valvano, M. A. A markerless deletion method for genetic manipulation of Burkholderia cenocepacia and other multidrug-resistant gram-negative bacteria. *Methods in molecular biology* **1197**, 311-327 (2014). <https://doi.org:10.1007/978-1-4939-1261-2_18>
